# Beyond the Catalytic Serine: Selective Protease Engagement with Covalent Macrocyclic Activity-Based Probes

**DOI:** 10.64898/2026.05.20.726481

**Authors:** Marina Barrueco, Ellie Hyde, Ella Sawtell, Jack V. Mills, Vaia Nikoloudi, Marianne Enget, Maisem Laabei, Markus Lakemeyer, Scott Lovell

## Abstract

Activity-based probes (ABPs) are widely used to profile serine protease activity – enzymes central to diverse physiological and pathological processes – but most rely on covalent modification of the conserved catalytic serine residue, often resulting in poor selectivity across related proteases.

Here, we introduce covalent macrocyclic activity-based probes (cmABPs) that selectively target non-catalytic residues within serine protease active sites. By combining phage display with systematic electrophile scanning, we identify macrocyclic scaffolds that position sulfur(VI) fluoride (SuFEx) electrophiles to covalently engage alternative nucleophiles such as lysine and tyrosine.

Applied to plasma kallikrein, this approach yielded a macrocyclic scaffold that was converted into covalent probes via fluorosulfate scanning. Remarkably, small changes in electrophile structure produced large, tuneable differences in covalent kinetics, with benzenesulfonyl fluoride derivative **23** achieving rapid and complete protein modification. Biochemical and mass spectrometry analyses confirmed selective modification of an active-site lysine by **23**, along with robust performance in complex biological samples. Extension to urokinase plasminogen activator further demonstrates the generality of this strategy.

More broadly, this work establishes electrophile scanning within macrocyclic scaffolds as a general approach for tuning covalent reactivity and provides a blueprint for designing selective probes that move beyond catalytic-residue targeting.

## Introduction

Activity-based probes (ABPs) have emerged as powerful chemical tools for profiling enzyme activities directly within complex biological systems. ABPs form irreversible covalent adducts with active enzymes, enabling selective visualization, enrichment, and quantification of enzymatic activities rather than protein abundances.^1^ Serine proteases, which regulate diverse physiological processes including coagulation, fibrinolysis, immunity, and cancer progression, have been among the most intensively studied enzyme classes using ABP technologies.^2-4^ Canonical serine protease ABPs typically consist of a short peptide recognition element coupled to an electrophilic warhead – most commonly a diaryl or mixed alkyl aryl phosphonate – that covalently modifies the catalytic serine residue within the active site (**Fig. 1a**).^5-7^ These reagents have enabled broad surveys of serine protease activity across tissues and disease states and have become foundational tools in chemical biology and proteomics.

**Figure 1:**
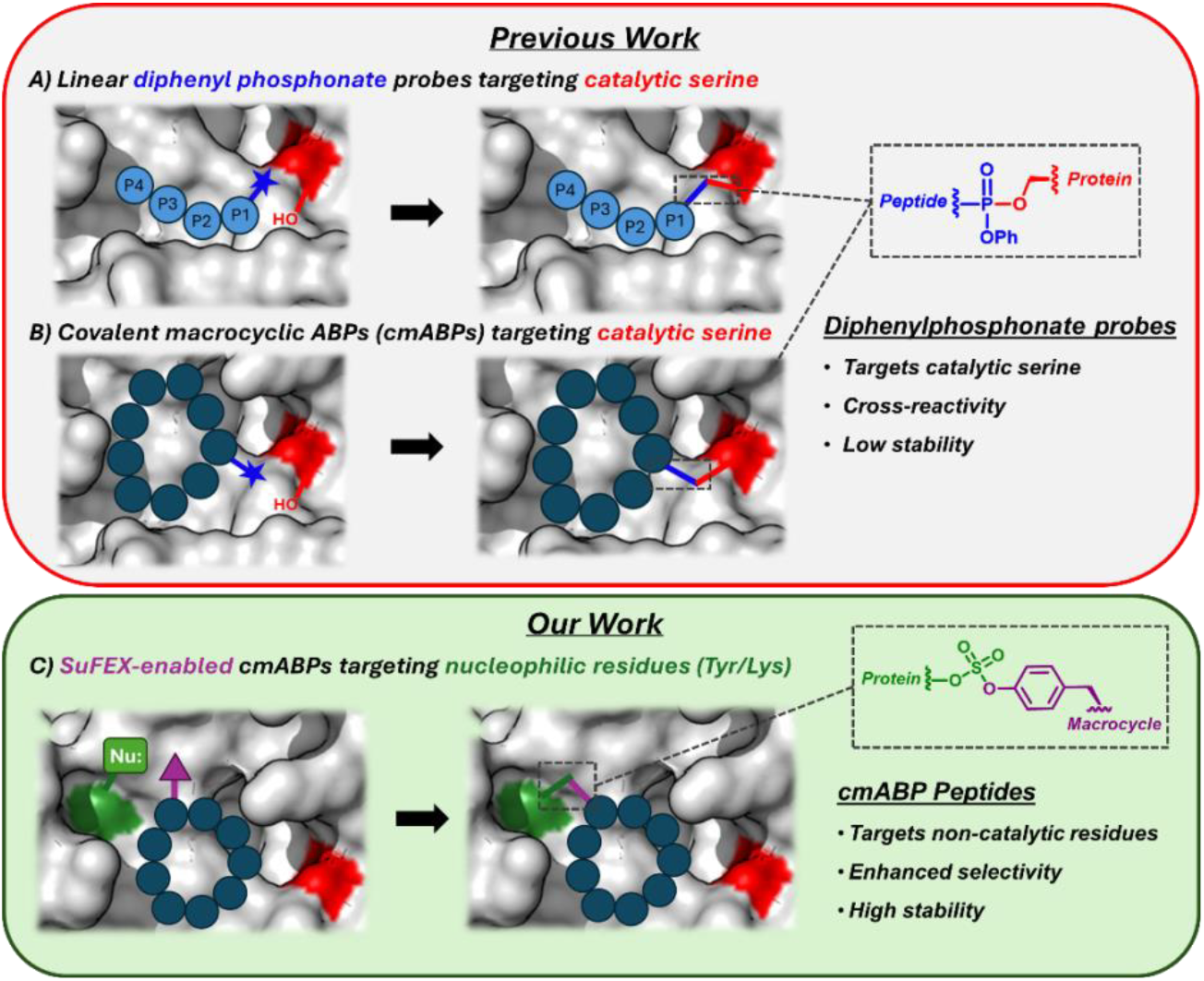
Conceptual framework for serine protease activity-based probe development. (**a**) Tetrapeptide-based probes bearing phosphonate warheads that covalently modify the catalytic serine. (**b**) Covalent macrocyclic ABPs (cmABPs) that enhance potency and selectivity through preorganization and precise placement of serine-directed electrophiles. (**c**) SuFEx-enabled cmABPs that selectively engage serine proteases via non-catalytic active-site residues.

Despite their utility, traditional tetrapeptide-based serine protease ABPs suffer from several notable limitations. First, selectivity remains a persistent challenge: the highly conserved S1-S4 substrate-binding pockets across serine protease families often result in extensive off-target labelling, particularly in proteome-wide applications. Second, phosphonate-based electrophiles, while selective toward catalytic serine residues, can exhibit limited chemical stability and unfavourable pharmacokinetic properties, complicating their use *in vivo*.^8^ In addition, many phosphonate-based probes contain up to two stereogenic centers at the phosphorus atom and adjacent carbon, resulting in complex mixtures of diastereoisomers that are often challenging to separate, purify, and rigorously validate in biochemical and biological assays.^9^ These drawbacks have motivated the development of alternative probe architectures that decouple target recognition and covalent reactivity while improving stability and precision.

Macrocyclization has emerged as a powerful strategy to preorganize binding motifs, reduce entropic penalties upon binding, and enable engagement of extended binding surfaces within enzyme active sites. Recent studies have demonstrated that covalent macrocycles can target proteases by positioning electrophiles with high spatial precision, resulting in potent engagement of catalytic residues (**Fig. 1b**).^10, 11^ However, many of these macrocyclic probes continue to rely on traditional serine-directed phosphonate warheads, thereby reinforcing the paradigm of covalent targeting of the conserved catalytic serine and potentially limiting *in vivo* applicability and proteome-wide selectivity. In contrast, alternative nucleophilic residues – including lysines, tyrosines, and histidines – are frequently positioned within protease active sites, differ across family members, and represent underexploited handles for achieving selective covalent engagement.^12, 13^

In this work, we establish a generalizable strategy to exploit these alternative nucleophilic residues through the development of covalent macrocyclic activity-based probes (cmABPs) that selectively engage serine proteases via non-catalytic active-site modification (**Fig. 1c**). By integrating phage display with systematic electrophile scanning, we identify cmABPs possessing sulfur(VI)-fluoride exchange (SuFEx) electrophiles,^14, 15^ for plasma kallikrein and urokinase plasminogen activator (uPA). Biochemical characterization of our lead plasma kallikrein cmABP, combining site-directed mutagenesis, intact-protein mass spectrometry and chemical proteomics demonstrates covalent modification of an active-site lysine residue rather than the canonical catalytic serine. We show that these probes exhibit high selectivity and can be deployed to measure protease activity in complex biological samples using activity-based protein profiling approaches.

## Results and Discussion

### De novo discovery of plasma kallikrein peptide binders by phage display

As a proof of concept for cmABP development, we initially focused on plasma kallikrein, a trypsin-like serine protease that plays a central role in the kallikrein–kinin system by regulating bradykinin production and vascular permeability, and whose dysregulation is directly implicated in hereditary angioedema and other inflammatory disorders.^16^ Although multiple peptide and small-molecule binders for plasma kallikrein have been reported,^17^ we avoided relying on existing ligands and instead sought to identify a reversibly binding peptide scaffold de novo. By doing so, and without prior knowledge of the detailed binding mode beyond engagement of the active site, we aimed to enable systematic electrophile scanning across the entire sequence and assess whether cmABPs could be discovered starting from unbiased binding information alone.

Accordingly, we employed phage display to identify macrocyclic peptide binders for plasma kallikrein. An AXCX_7_C phage-displayed peptide library (A = Alanine and X = any of the 20 naturally occurring amino acids), was cyclized using 1,3-bis(bromomethyl)benzene (DBMB) to generate constrained macrocyclic architectures (**Fig. 2a**). Three rounds of affinity selection (‘panning’) were performed against biotinylated plasma kallikrein immobilized on streptavidin-coated beads, alongside a no-protein control to account for non-specific enrichment. After three rounds of panning, an approximately 1000-fold enrichment in phage titres was observed for the plasma kallikrein selection relative to the control, consistent with enrichment of specific binders (**Fig. 2b**). Next-generation sequencing of phage DNA from the final round revealed approximately 400 peptide sequences that were enriched at least threefold in the plasma kallikrein selection compared to the no-protein control (**Fig. S1**). Clustering of enriched sequences using GibbsCluster^18^ identified four distinct peptide clusters, all of which converged on a conserved Leu/Phe/Pro–Trp/Phe–Arg motif (**Fig. 2c**). This motif corresponds to a known plasma kallikrein substrate preference, with P1-Arg, P2-Phe/Trp, and P3-Leu/Phe/Pro, providing strong evidence that the selected peptides engage the protease active site.^19^ To identify suitable scaffolds for cmABP development, two representative peptides from each of the four clusters were selected, synthesized, and evaluated for their ability to inhibit plasma kallikrein activity. Of the eight peptides tested, three inhibited plasma kallikrein activity by more than 50% at a concentration of 1 μM, indicating productive engagement of the protease active site (**Fig. 2d**). Among these, peptide **3**, which contains the Leu–Phe–Arg motif and exhibited a K_i_ of 830 nM, was selected as the starting scaffold for subsequent electrophile scanning and cmABP development (**Fig. 2e**).

**Figure 2:**
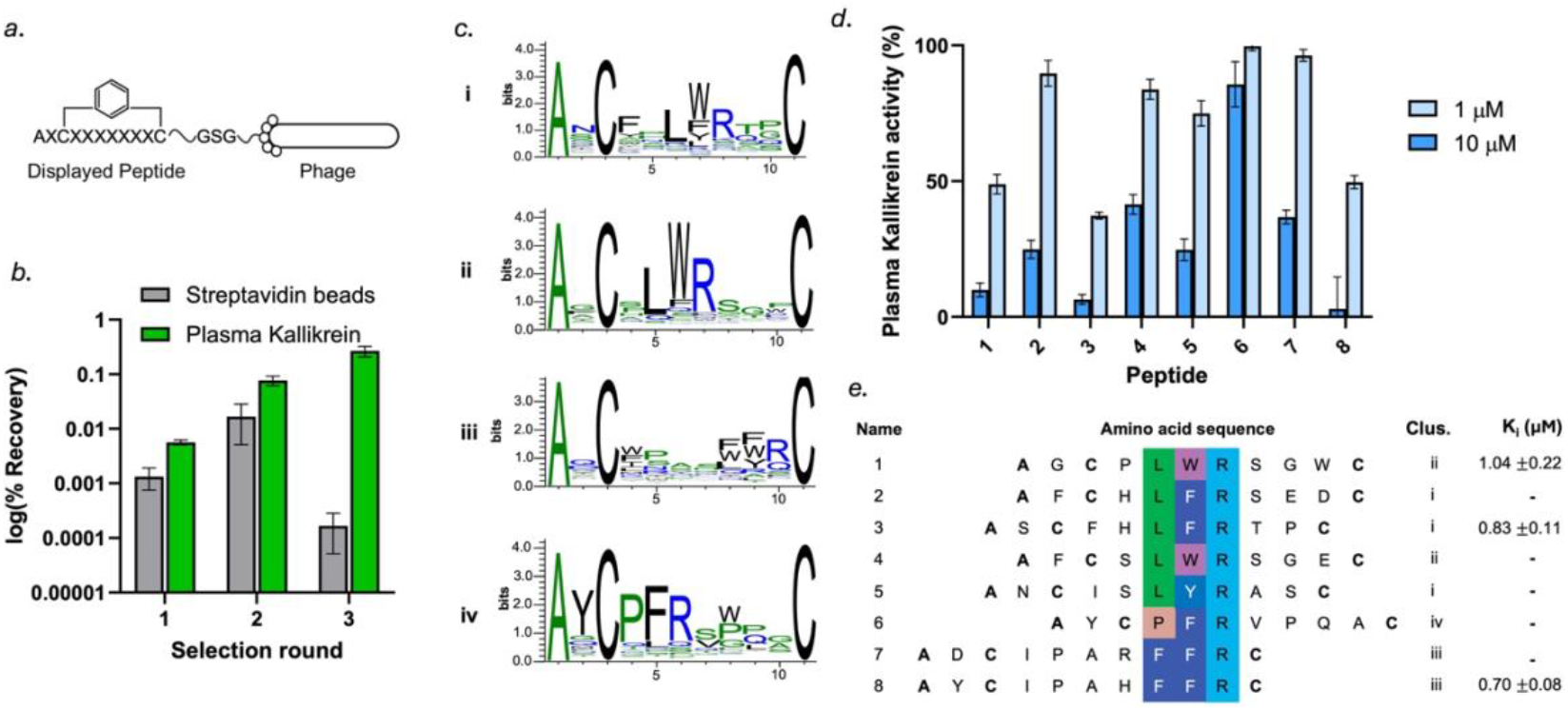
De novo identification of plasma kallikrein peptide binders. (**a**) Structure of the DBMB-cyclised AXCX_7_C phage display library. **b**) Percentage recovery of AXCX_7_C phage display library over three rounds of phage panning against plasma kallikrein or streptavidin beads **c**) Sequence logo representation of enriched macrocyclic peptides following three rounds of phage display against plasma kallikrein. (**d**) Inhibition of plasma kallikrein activity by representative peptides selected from each cluster, measured at 10 and 1 μM concentration. (**e**) Sequences of representative peptides and corresponding K_i_ values, identical amino acids are highlighted in colour.

### Fluorosulfate scanning of plasma kallikrein inhibitor 3

To convert the reversible plasma kallikrein binder into a cmABP, we performed a systematic electrophile scan of peptide **3**, in which fluorosulfate tyrosine (FSY) was introduced at each position along the peptide sequence. A propargyl glycine residue was also incorporated into the sequence in place of the N-terminal alanine residue to enable downstream activity-based protein profiling (ABPP) (**Fig. 3a-b**).^20^ In the absence of detailed structural information defining the precise binding mode of **3**, this comprehensive scanning strategy was designed to identify positions from which the fluorosulfate electrophile could productively engage nucleophilic residues within the plasma kallikrein active site. FSY was selected as the electrophile due to its high plasma stability, low intrinsic reactivity, compatibility with solid-phase peptide synthesis (SPPS), and capacity to form covalent adducts with Tyr, His, and Lys residues.^21-23^

**Figure 3:**
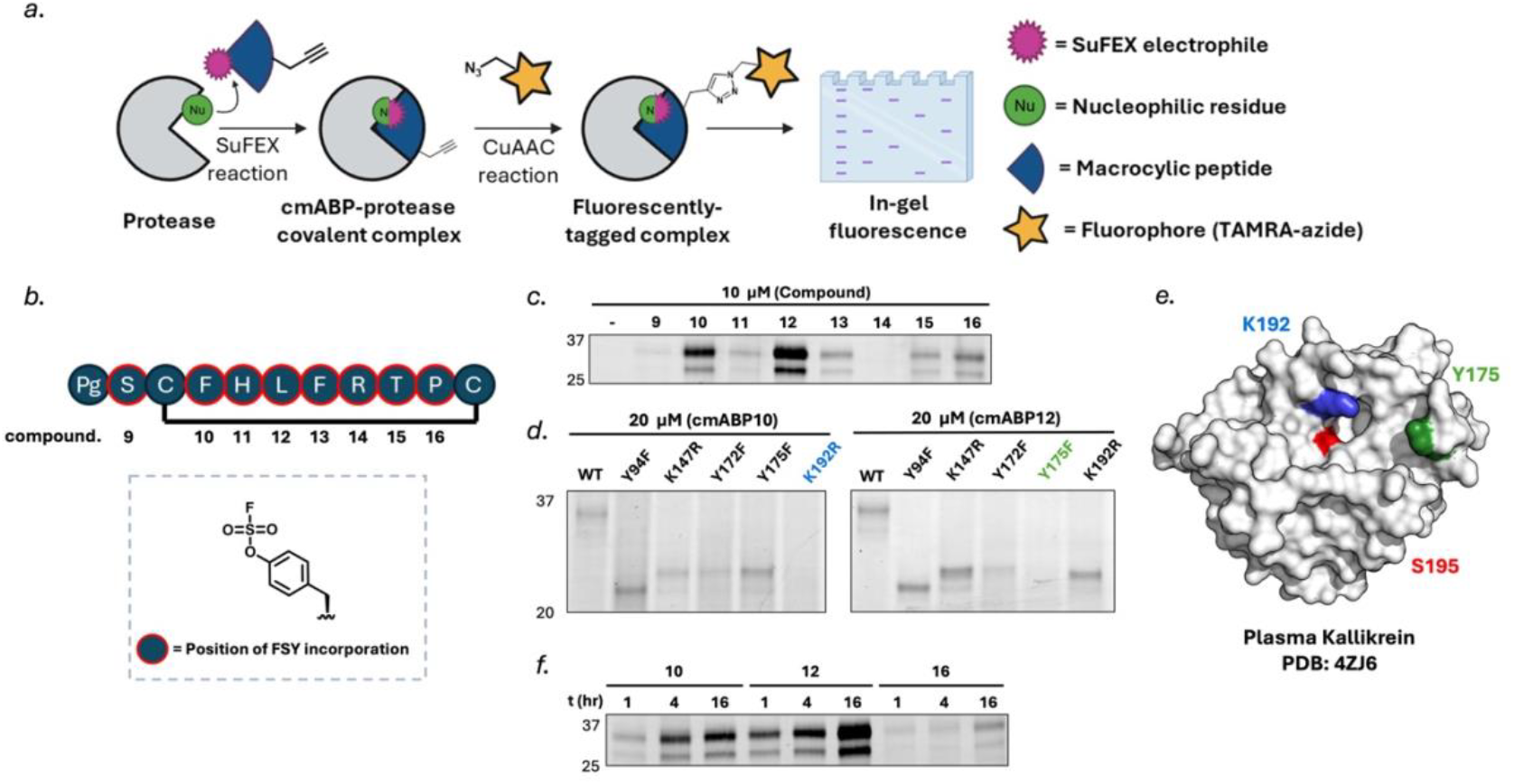
Fluorosulfate scanning of plasma kallikrein peptide inhibitor 3. (**a**) Schematic of the activity-based protein profiling workflow used to detect probe–protease adducts. (**b**) Structures of fluorosulfate-containing cmABP derivatives **9**-**16**. (**c**) In-gel fluorescence analysis of plasma kallikrein labelling (1 µM) following incubation with 20 µM cmABP **9-16** for 16 h at 37 ºC. (**d**) Covalent labelling of plasma kallikrein active-site residue mutants by cmABP derivatives **10** and **12**. (**e**) Crystal structure of Plasma Kallikrein (PDB Accession: 4ZJ6), highlighting the catalytic serine (S195, red) and nucleophilic residues targeted by cmABP10 (K192, blue) and cmABP12 (Y175, green). (**f**) Time-dependent labelling of plasma kallikrein using 20 µM **10, 12** and **16** at 37 ºC.

The eight covalent derivatives of **3** (**9**-**16**) were synthesized using standard Fmoc-based SPPS conditions, with the exception that 2-methylpiperidine was used in place of piperidine during Fmoc deprotection steps to preserve the integrity of the fluorosulfate group (**Scheme S1-2**).^24^ Following chain assembly, linear peptides were cleaved from the resin using a trifluoroacetic acid (TFA) cleavage cocktail and subsequently cyclized in solution using DBMB to afford the desired covalent macrocyclic products (**Scheme S3**).

Each cmABP derivative (5 μM) was incubated with plasma kallikrein (1 μM) for 16 h at 37 °C, followed by ligation of an azide–TAMRA reporter via copper-catalysed azide–alkyne cycloaddition (CuAAC). Formation of probe-protease adducts was visualized by in-gel fluorescence, revealing detectable covalent labelling when FSY was incorporated at positions corresponding to Phe-4 (**10**), Leu-6 (**12**), and to a lesser extent Pro-10 (**16**) (**Fig. 3c**). Two plasma kallikrein bands were consistently observed by in-gel fluorescence, consistent with previously reported heterogeneous glycosylation of the protease.^25^ Substitution at other positions did not result in appreciable labelling, likely reflecting suboptimal electrophile positioning relative to active-site nucleophiles or local microenvironment effects that attenuate nucleophile reactivity.

To identify the site of covalent modification, a panel of plasma kallikrein mutants (H40N, Y94F, K147R, Y172F, Y175F, and K192R) was expressed and evaluated for probe labelling. In all mutants, glycosylation sites were mutated (N377E, N434E, N475E) to generate homogeneous protein for intact mass spectrometry analysis. All mutants retained catalytic activity, as confirmed by labelling with fluorophosphonate–TAMRA (FP-TAMRA) (**Fig. S2**), apart from H40N, suggesting this residue plays a key role in catalysis. Mutational analysis revealed that labelling by **10** was selectively diminished for the K192R mutant, identifying Lys192 as the primary site of covalent modification (**Fig. 3d**, left). In contrast, substitution of Y175 abrogated labelling by **12**, indicating covalent modification of this residue (**Fig. 3d**, right). Y175 is located at the base of the S3 pocket, consistent with binding of the Leu-6 side chain and positioning of the electrophile for covalent engagement. These results demonstrate that distinct electrophile placements within the macrocyclic scaffold enable selective engagement of different non-catalytic nucleophilic residues within the plasma kallikrein active site (**Fig. 3e**).

Time-dependent labelling experiments further differentiated the two probes, with derivative **12** exhibiting more efficient covalent labelling than **10** (**Fig. 3f**). However, enzyme inhibition studies revealed a marked discrepancy between covalent labelling efficiency and binding affinity: after 30-minutes pre-incubation with plasma kallikrein, derivative **12** displayed a K_i_ > 50 μM, indicating poor non-covalent engagement and limited tolerance for electrophile incorporation at this position, whereas derivative **10** retained substantially higher affinity (K_i_ = 1.5 μM) (**Fig. S3**). These data suggest that, despite slower covalent labelling, placement of the fluorosulfate electrophile at the Phe-4 position is better accommodated within the binding pocket, and derivative **10** was therefore selected for further optimization.

### Optimization of covalent engagement through electrophile diversification

Although fluorosulfate scanning identified derivative **10** as a potential cmABP for plasma kallikrein, in-gel fluorescence analysis revealed that covalent modification of the protease was incomplete even after prolonged incubation (16 h), suggesting a relatively slow rate of covalent bond formation. Given that efficient activity-based protein profiling relies not only on selectivity but also on sufficiently rapid covalent engagement under biologically relevant conditions, we sought to further optimize the electrophilic warhead to enhance the rate of covalent modification.

To this end, we systematically evaluated a panel of alternative sulfur(VI) fluoride electrophiles, including substituted fluorosulfates and sulfonyl fluorides, primarily installed at the position corresponding to derivative **10**, while also probing adjacent positions to assess positional tolerance (**Fig. 4a**). These electrophiles span a range of intrinsic reactivities and steric profiles,^14^ enabling modulation of covalent engagement kinetics without altering the underlying peptide recognition scaffold. This strategy was designed to identify cmABPs that achieve more complete and rapid covalent modification of plasma kallikrein, while retaining the favourable selectivity and stability characteristics of the macrocyclic probe architecture.

**Figure 4:**
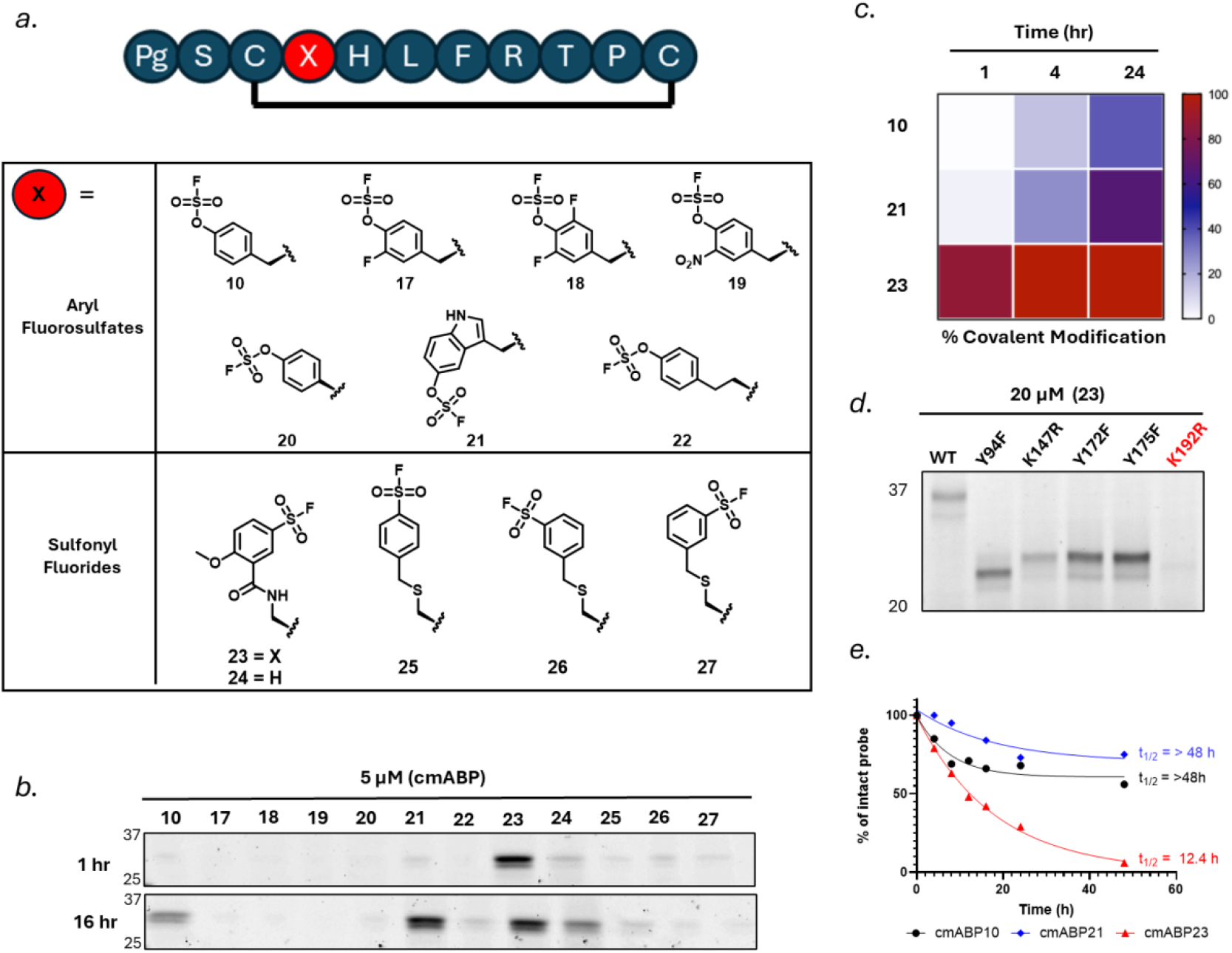
Electrophile diversification of plasma kallikrein cmABP 10. (**a**) Structures of cmABP **10** derivatives bearing fluorosulfate and sulfonyl fluoride electrophiles. (**b**) In-gel fluorescence analysis of plasma kallikrein labelling (1μM) following incubation with 5 µM cmABP derivative for 1 h or 16 h at 37 ºC. (**c**) Intact-protein mass spectrometry analysis of covalent modification kinetics for **10, 21**, and **23**. (**d**) Covalent labelling of plasma kallikrein active-site residue mutants by cmABP **23**. (**e**) Stability of cmABPs **10, 21**, and **23** in PBS (pH 7.4).

Fluorosulfate-containing analogues **17–22** were prepared by incorporating unprotected phenolic residues during Fmoc-based SPPS. Following completion of linear peptide assembly, the resin-bound peptides were treated overnight with 20% piperidine in DMF to remove any acylation of the phenol side chain, after which on-resin conversion to the corresponding fluorosulfates was achieved using 4-(Acetylamino)phenyl]imidodisulfuryl difluoride (AISF) (**Scheme S4**).^26^ Sulfonyl fluoride analogues **23** and **24** were synthesised using MTT-protected diaminopropionic acid (Dap), which after orthogonal deprotection using 3% TFA was functionalised by on-resin amidation with 5-(fluorosulfonyl)-2-methoxybenzoic acid (**Scheme S5**). Finally, sulfonyl fluoride analogues **25–27** were generated via incorporation of STMP-protected Cys into the linear peptide,^27^ followed by orthogonal deprotection using DTT and on-resin alkylation with the corresponding (bromomethyl)benzenesulfonyl fluorides (**Scheme S6**). In all cases, final products were obtained following TFA-mediated cleavage and solution-phase cyclization using DBMB.

Covalent labelling efficiency of the electrophile-diversified cmABPs was evaluated using the in-gel fluorescence assay described above. Analogues bearing ortho-fluoro (**17**), ortho-difluoro (**18**) or ortho-nitro substitutions (**19**), as well as fluorosulfate– phenylglycine (**20**) and homo-fluorosulfate tyrosine (**22**), showed reduced covalent labelling relative to **10**. In contrast, incorporation of 5-fluorosulfate–tryptophan (**21**) resulted in a marked increase in covalent modification of plasma kallikrein. Among the sulfonyl fluoride-containing cmABPs, derivatives **25–27** also displayed reduced covalent labelling, whereas sulfonyl fluoride **23** produced a pronounced increase in labelling efficiency, exceeding that observed for all other electrophiles tested. Installation of the same sulfonyl fluoride electrophile used in **23** at the His-5 position (compound **24**) also resulted in robust covalent labelling, albeit at a reduced level, suggesting that productive engagement of the target nucleophilic residue can also be achieved from an adjacent position within the peptide scaffold (**Fig. 4b**).

To enable a more quantitative comparison of covalent modification kinetics, intact-protein mass spectrometry was performed for derivatives **10, 21**, and **23** (**Fig. 4c**). Plasma kallikrein was incubated with a 10-fold molar excess of each cmABP in PBS (pH 7.4) at 37 °C, and the extent of covalent modification was quantified after 1, 4, and 24 h. Consistent with the in-gel fluorescence data, derivative **10** exhibited slow covalent engagement, with no detectable modification after 1 h, increasing to 15% at 4 h and 38% after 24 h. In contrast, derivative **21** showed enhanced kinetics, with 3% modification at 1 h, 27% at 4 h, and 66% at 24 h. Notably, sulfonyl fluoride **23** displayed rapid and near-complete covalent modification, achieving 87% modification within 1 h and complete modification by 4 h, which was maintained at 24 h.

To confirm the site of covalent modification by **23**, a follow-up mutational analysis was performed using the panel of plasma kallikrein variants described above. Consistent with the results obtained for **10**, substitution of Lys192 (K192R) resulted in a marked loss of labelling by **23**, indicating that this residue remains the primary site of covalent engagement (**Fig. 4d**). This assignment was further supported by tryptic digest followed by LC-MS/MS analysis of plasma kallikrein treated with **23**, which confirmed modification at Lys192 (**Fig. S4-5**).

To assess the chemical stability of the electrophile-diversified cmABPs, derivatives **10, 21**, and **23** were incubated in PBS (pH 7.4), and rates of hydrolysis were determined by monitoring probe depletion using HPLC. Under these conditions, fluorosulfate-containing derivatives **10** and **21** exhibited comparatively high stability, with half-lives of >48 h. Sulfonyl fluoride **23** exhibited moderate aqueous stability with a half-life of ∼12.5 h, lower than that of the corresponding fluorosulfates, yet sufficient to support covalent labelling under the conditions employed, warranting further evaluation in more complex biological environments (**Fig. 4e**).

Taken together, these results establish systematic diversification of sulfur(VI) fluoride electrophiles within a macrocyclic scaffold as a powerful and still largely untapped strategy for controlling covalent engagement. Notably, even a limited fluorosulfate scan delivered an improved covalent binder, with derivative **21** exhibiting enhanced modification kinetics relative to **10**, underscoring the high sensitivity of covalent engagement to electrophile structure. This suggests that broader exploration of SuFEx chemical space could yield cmABPs that approach an optimal balance of covalent reactivity, selectivity, and stability.^28^ Sulfonyl fluoride **23** clearly emerges as the most effective probe identified in this study, achieving rapid and near-complete covalent modification of Lys192 and representing a promising chemical tool for probing plasma kallikrein activity, albeit with remaining questions regarding the chemical stability of the electrophile in complex biological environments. Guided by these insights, we next evaluated the selectivity of derivatives **21** and **23** in more complex biochemical and biological environments.

### Biochemical validation of plasma kallikrein cmABPs

We next evaluated the selectivity of **23** against a panel of serine proteases using complementary biochemical and activity-based assays. Each protease (500 nM) was incubated with **23** (5 μM) for 1 h, followed by CuAAC-mediated ligation of an azide–TAMRA reporter and analysis by in-gel fluorescence. Under these stringent labelling conditions, **23** displayed strong and selective labelling of plasma kallikrein with minimal off-target labelling across the protease panel (**Fig. 5a**). Consistent with these findings, analogous in-gel fluorescence experiments with fluorosulfate **21** also revealed a high degree of selectivity for plasma kallikrein (**Fig. S6**). In contrast, FP-TAMRA exhibited extensive labelling across the protease panel (**Fig. S7**). We also synthesized fluorescent peptidyl diphenylphosphonate probe **28 (Scheme S7-8)**, which possesses the preferred P1–P4 amino acid residues for plasma kallikrein; however, this probe exhibited only modest labelling of plasma kallikrein and substantial off-target labelling of uPA, trypsin, and KLK2 (**Fig. 5b-c**). In enzyme activity assays, incubation with **23** for 30 min at 37 °C resulted in potent inhibition of plasma kallikrein (K_i_ = 31 ± 2 nM), with at least 200-fold selectivity over all other proteases tested (**Fig. 5d**). These findings highlight the advantages of cmABPs, which achieve enhanced selectivity by integrating extended binding interactions and precise electrophile positioning, rather than relying on short linear recognition sequences coupled to serine-directed warheads with high intrinsic reactivity.

**Figure 5:**
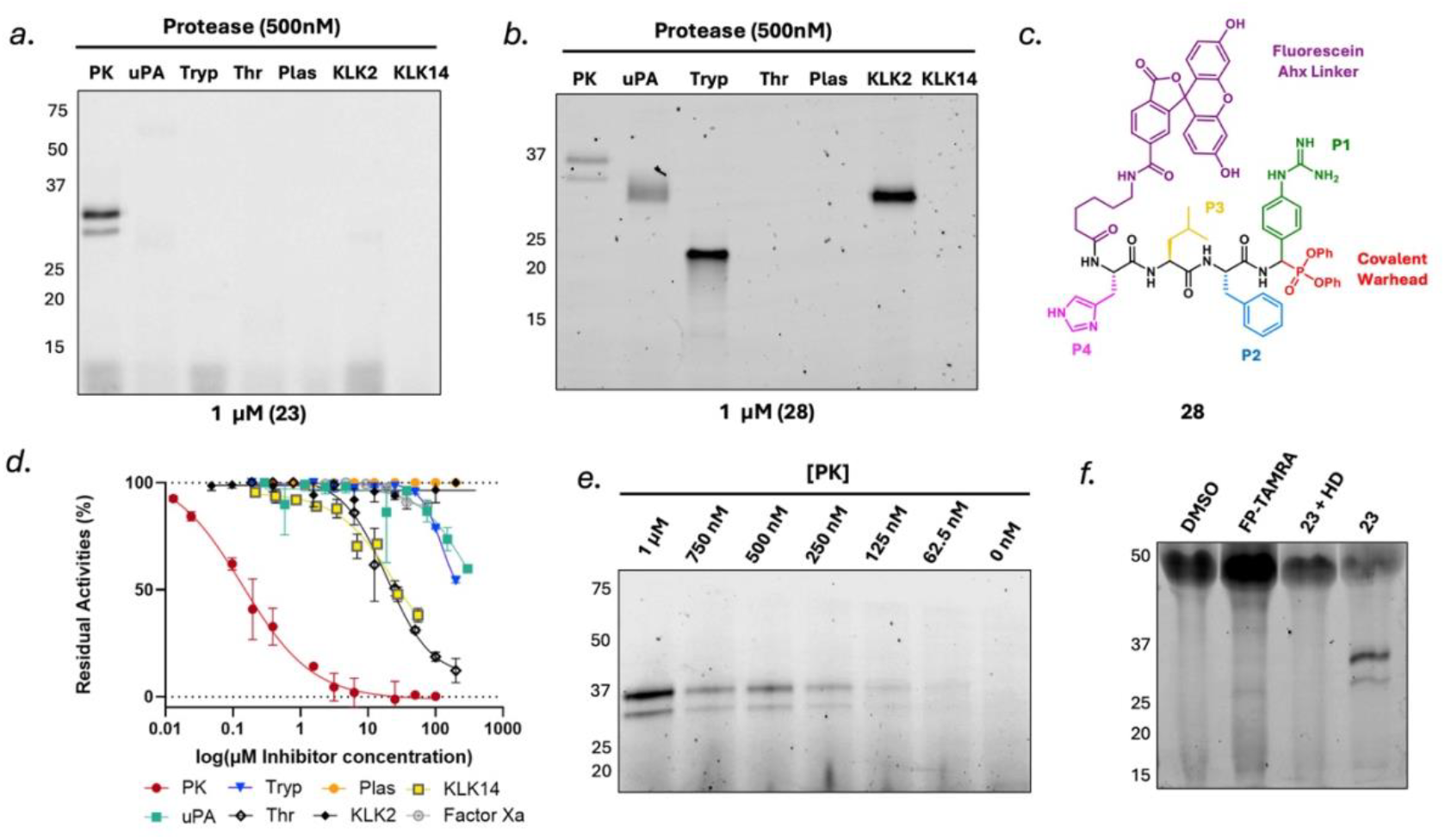
Biochemical validation and selectivity of plasma kallikrein cmABPs. (**a**) In-gel fluorescence analysis of protease labelling by **23** (**b**) In-gel fluorescence analysis of protease labelling by **28**; PK = plasma kallikrein, uPA = urokinase plasminogen activator, Tryp = trypsin, Thr = Thrombin, Plas = Plasmin, KLK = kallikrein (**c**) Structure of diphenyl phosphonate probe **28**; Ahx = 6-amino-hexanoic acid (**d**) Inhibition of protease activity by **23** following 30 min incubation at 37 ºC (**e**) Detection of plasma kallikrein activity in PC3-conditioned media using 1 µM **23** (**f**) Detection of plasma kallikrein activity in human plasma using 1 µM **23** or FP-TAMRA, HD = heat denatured.

To evaluate cmABP performance in a more complex biological context, we next assessed the ability of **23** to selectively detect plasma kallikrein activity in conditioned media from PC3 cells, which have been reported to contain high levels of endogenous proteases, including uPA.^29^ Decreasing amounts of plasma kallikrein were spiked into the conditioned media, spanning concentrations from 1 μM to 62.5 nM, and samples were analysed using the activity-based labelling workflow described above. Selective labelling of plasma kallikrein was detected down to 62.5 nM, corresponding to approximately 0.17% of total protein (**Fig. 5e**), whereas FP-TAMRA produced multiple fluorescent bands (**Fig. S8**). Finally, we evaluated probe performance in human plasma. Incubation with **23** yielded two prominent fluorescent bands at the expected molecular weight of plasma kallikrein (**Fig. 5f**), which were abolished upon heat denaturation, confirming activity-dependent labelling. In contrast, FP-TAMRA showed minimal labelling, consistent with reduced electrophile stability in this environment.

Collectively, these findings demonstrate that **23** is sufficiently stable to support covalent labelling in plasma, positioning it as a functionally robust probe for detecting plasma kallikrein activity in complex biological matrices.

### Extension of cmABP strategy to urokinase plasminogen activator (uPA)

To demonstrate the generality of the cmABP discovery strategy, we next applied this approach to urokinase plasminogen activator (uPA), a trypsin-like serine protease with well-established roles in fibrinolysis and cancer progression.^30^ In contrast to the de novo discovery approach used for plasma kallikrein, we leveraged a previously reported bicyclic peptide inhibitor **29** as a structurally defined starting scaffold, which binds uPA with high potency and selectivity through extensive engagement of the active site (**Fig. 6a**).^31^

**Figure 6:**
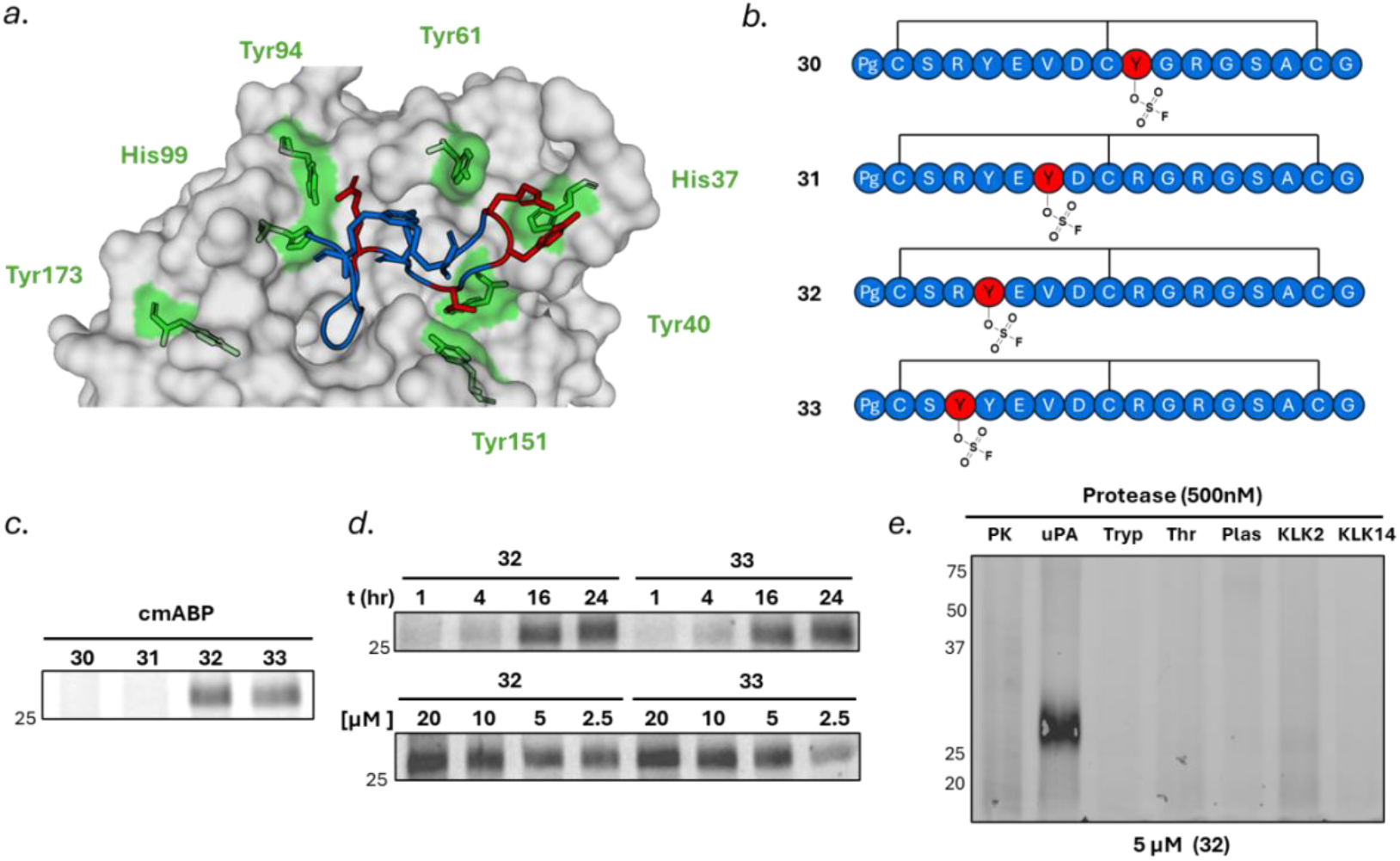
Fluorosulfate scanning of bicyclic peptide inhibitor 1. (**a**) Co-crystal structure of uPA bound to bicyclic peptide inhibitor **29** (blue) (PDB Accession: 4JK6), highlighting the positions selected for FSY substitution (red) and nucleophilic residues within 10 Å (green) (**b**) Structures of cmABP **30-33**, Pg = propargyl glycine (**c**) In-gel fluorescence analysis of uPA labelling following incubation with cmABP derivatives **30–33** (**d**) Time-(top) and concentration-(bottom) dependent labelling of uPA by derivatives **32** and **33** (**e**) Selectivity of cmABP **32** across a panel of structurally related serine proteases; PK = plasma kallikrein, uPA = urokinase plasminogen activator, Tryp = trypsin, Thr = Thrombin, Plas = Plasmin, KLK = kallikrein.

Guided by the co-crystal structure of uPA in complex with **29**, we identified Arg4, Tyr5, Val7, and Arg10 as suitable positions for incorporation of fluorosulfate tyrosine (FSY), based on the proximity and orientation of these residues toward nucleophilic Tyr, His, or Lys side chains within the active site. The corresponding covalent derivatives (**30–33**) were synthesized as above, with a propargyl glycine residue incorporated into each derivative to facilitate gel-based ABPP (**Fig. 6b**). Following chain assembly, linear peptides were cleaved from the resin and cyclized in solution using tribromomethylbenzene (TBMB) to afford the desired covalent bicycles (**Scheme S1**).

Covalent engagement of uPA was evaluated using in-gel fluorescence following incubation of each cmABP derivative (5 μM) with uPA (300 nM) for 16 h at 37 °C and subsequent CuAAC-mediated ligation of an azide–TAMRA reporter. Robust labelling was observed for derivatives **32** and **33**, whereas no detectable labelling was observed for **30** and **31** (**Fig. 6c**). Time- and concentration-dependent labelling experiments further differentiated the active probes, with derivative **32** exhibiting slightly more efficient covalent modification than **33** (**Fig. 6d**). These results indicate that substitution at the Tyr5 position enables productive positioning of the SuFEx electrophile for covalent engagement with a nucleophilic residue (likely His37 or Tyr40) in the uPA active site. However, the relatively slow rate of covalent modification observed for these probes suggests that further optimisation of electrophile structure, for example through systematic electrophile diversification, will be required to enhance covalent engagement kinetics, as demonstrated for plasma kallikrein. Despite these limitations, in-gel fluorescence screening against a panel of structurally related serine proteases revealed that cmABP **32** exhibits exquisite selectivity for uPA (**Fig. 6e**), underscoring the ability of the macrocyclic scaffold to confer high target specificity even in the absence of fully optimised covalent reactivity.

Together, these results demonstrate that covalent macrocyclic probes can be rationally developed to selectively engage serine proteases through non-catalytic active-site residues, and that systematic tuning of sulfur(VI) fluoride electrophiles provides a general strategy to balance covalent reactivity, selectivity, and stability across distinct protease targets.

## Conclusion

In summary, we establish cmABPs as a powerful modality for selectively targeting serine proteases through non-catalytic active-site residues. By integrating phage display or structure-guided ligand identification with systematic electrophile scanning, we demonstrate that covalent engagement can be achieved through precise positioning of sulfur(VI) fluoride warheads toward alternative nucleophilic residues, including lysines and tyrosines. Application of this approach to plasma kallikrein enabled the identification of cmABPs with high selectivity and tuneable covalent kinetics, with electrophile diversification providing a powerful means to modulate the balance between reactivity, selectivity, and chemical stability. Notably, these results imply that covalent engagement is strongly influenced by the trajectory of the SuFEx moiety, where optimised orientation and approach can substantially enhance reactivity (e.g., cmABP **21** vs **10**), underscoring the importance of broader electrophile exploration. Extension to uPA further highlights the generality of this strategy across distinct protease targets, even where covalent reactivity remains to be fully optimized. More broadly, this work establishes macrocycle-enabled covalent targeting as a versatile platform for developing selective chemical tools to interrogate protease activity in complex biological systems.

## Supporting information

Proteomics Data

## Acknowledgments

The authors thank Dr Kate Heesom (Proteomics Facility, University of Bristol) for assistance in acquiring mass spectrometry data. S.L is grateful for funding from the Engineering and Physical Sciences Research Council (UKRI117) and Medical Research Council (UKRI2332). M.LK is grateful for support by the research profile line LIFE of FSU Jena and the Deutsche Forschungsgemeinschaft (DFG, German Research Foundation, Project ID 528114058).

## Author Contributions

Author contributions are assigned using Contributor Role Taxonomy (CRediT)

**M.B:** Data Curation, Investigation, Methodology, Visualisation, Formal Analysis, Writing – original draft, Writing – review and editing. **E.H**: Data Curation, Investigation, Methodology, Visualisation, Writing – review and editing. **E.S:** Data Curation, Investigation, Methodology. **J.M:** Data Curation, Investigation, Methodology. **V.N:** Data Curation, Investigation, Methodology. **M.E:** Data Curation, Investigation, Methodology. **M.LK:** Investigation, Formal Analysis, Methodology. **M.LB** Investigation, Formal Analysis, Methodology. **S.L:** Conceptualisation, Funding Acquisition, Supervision, Writing – original draft, Writing – review and editing.

## References

1. F. Faucher, J.M. Bennett, M. Bogyo and S. Lovell, Strategies for Tuning the Selectivity of Chemical Probes that Target Serine Hydrolases, Cell Chem. Biol., 2020, 27, 937–952

2. P. Kasperkiewicz, Y. Altman, M. D’Angelo, G. S. Salvesen and M. Drag, Toolbox of Fluorescent Probes for Parallel Imaging Reveals Uneven Location of Serine Proteases in Neutrophils, J. Am. Chem. Soc., 2017, 139, 10115–10125

3. S. Modrzycka, S. Kolt, S. G. I. Polderdijk, T. E. Adams, S. Potoczek, J. A. Huntington, P. Kasperkiewicz and M. Drag, Parallel imaging of coagulation pathway proteases activated protein C, thrombin, and factor Xa in human plasma, Chem. Sci., 2022, 13, 6813–6829

4. T. Kryza, N. Bock, S. Lovell, A. Rockstroh, M. L. Lehman, A. Lesner, J. Panchadsaram, M. L. Silva, S. Srinivasan, C.E. Snell, E. D. Williams, L. Fazli, M. Gleave, J. Batra, C. Nelson, E. W. Tate, J. Harris, J. D. Hooper, and J. A. Clements, The molecular function of kallikrein-related peptidase 14 demonstrates a key modulatory role in advanced prostate cancer, Mol. Oncol., 2020, 14, 105–128

5. S. Lovell, L. Zhang, T. Kryza, A. Neodo, N. Bock, E. D. Vita, E. D. Williams, E. Engelsberger, C. Xu, A. T. Bakker, M. Maneiro, R. J. Tanaka, C. L. Bevan, J. A. Clements and E. W. Tate, A Suite of Activity-Based Probes To Dissect the KLK Activome in Drug-Resistant Prostate Cancer, J. Am. Chem. Soc., 2021, 143, 8911–8924

6. S. Ji and S. H. L. Verhelst, Furin-targeting activity-based probes with phosphonate and phosphinate esters as warheads, Org. Biomol. Chem., 2023, 21, 6498–6502

7. S. Serim, P. Baer and S. H. L. Verhelst, Mixed alkyl aryl phosphonate esters as quenched fluorescent activity-based probes for serine proteases, Org. Biomol. Chem., 2015, 13, 2293–2299

8. J. Ides, D. Thomae, L. wyffels, C. Vangestel, J. Messagie, J. Joossens, F. Lardon, P. V. d. Veken, K. Augustyns, S. Stroobants and S. Staelens, Synthesis and in vivo preclinical evaluation of an 18F labeled uPA inhibitor as a potential PET imaging agent, Nucl. Med. Biol., 2014, 41, 477–487

9. J. P. Kahler and S. H. L. Verhelst, Phosphinate esters as novel warheads for activity-based probes targeting serine proteases, RSC Chem. Biol., 2021, 2, 1285–1290

10. T. Lan, C. Peng, X. Yao, R. S. T. Chan, T. Wei, A. Rupanya, A. Radakovic, S. Wang, S. Chen, S. Lovell, S. A. Snyder, M. Bogyo and B. C. Dickinson, Discovery of Thioether-Cyclized Macrocyclic Covalent Inhibitors by mRNA Display, J. Am. Chem. Soc., 2024, 146, 24053–24060

11. S. Chen, S. Lovell, S. Lee, M. Fellner, P. D. Mace and M. Bogyo, Identification of highly selective covalent inhibitors by phage display, Nat. Biotechnol., 2021, 39, 490–498

12. Y. Yang, P. Liu, X. Y. Cui, J. Wu, W. Cai, J. Cheng, C. Lu, X. Fan, Y. Zhang, Z. Liu and Y. Yin, Ribosomal Incorporation of Fluorosulfonyloxy-l-Phenylalanine into Macrocyclic Peptides for De Novo Target-Specific Covalent Binders, J. Am. Chem. Soc., 2025, 147, 27095–27104

13. K. S. V. Horn, D. Wang, D. Medina-Cleghorn, P. S. Lee, C. Bryant, C. Altobelli, P. Jaishankar, K. K. Leung, R. A. Ng, A. J. Ambrose, Y. Tang, M. R. Arkin and A. R. Renslo, Engaging a Non-catalytic Cysteine Residue Drives Potent and Selective Inhibition of Caspase-6, J. Am. Chem. Soc., 2023, 145, 10015–10021

14. K. E. Gilbert, A. Vuorinen, A. Aatkar, P. Pogány, J. Pettinger, E. K. Grant, J. M. Kirkpatrick, K. Rittinger, D. House, G. A. Burley and J. T. Bush, Profiling Sulfur(VI) Fluorides as Reactive Functionalities for Chemical Biology Tools and Expansion of the Ligandable Proteome, ACS Chem. Biol., 2023, 18, 285–295

15. L. Hillebrand, X. J. Liang, R. A. M. Serafim and M. Gehringer, Emerging and Re-emerging Warheads for Targeted Covalent Inhibitors: An Update, J. Med. Chem., 2024, 67, 7688–7758

16. G. Motta, L. Juliano and J. R. Chagas, Human plasma kallikrein: roles in coagulation, fibrinolysis, inflammation pathways, and beyond, Front. Physiol., 2023, 14, 1188816

17. H. Liu, Y. Deng, J. Liu, Z. Wang, X.-Q. Hu, Y. Duan, Y. Chen and Z. Xie, Plasma Kallikrein Inhibitors for Multiple Disorders: Current Advances and Perspectives, J. Med. Chem., 2025, 68, 21012–21034

18. M. Andreatta, B. Alvarez and M. Nielsen, GibbsCluster: unsupervised clustering and alignment of peptide sequences, Nucleic Acids Res., 2017, 45, 458–463

19. A. R. Lima, F. M. Alves, P. F. Angelo, D. Andrade, S. I. Blaber, M. Blaber, L. Juliano and M. A. Juliano, S(1)’ and S(2)’ subsite specificities of human plasma kallikrein and tissue kallikrein 1 for the hydrolysis of peptides derived from the bradykinin domain of human kininogen, Biol. Chem., 2008, 389, 1487–1494.

20. S. Wang, F. F. Faucher, M. Bertolini, H. Kim, B. Yu, L. Cao, K. Roeltgen, S. Lovell, V. Shanker, S. D. Boyd, L. Wang, R. Bartenschlager and M. Bogyo, Identification of Covalent Cyclic Peptide Inhibitors Targeting Protein–Protein Interactions Using Phage Display, J. Am. Chem. Soc., 2025, 147, 7461–7475.

21. P. D. P. Martín-Gago and P. D. C. A. Olsen, Arylfluorosulfate-Based Electrophiles for Covalent Protein Labeling: A New Addition to the Arsenal, Angew. Chem. Int. Ed., 2018, 58, 957–966

22. G. Alboreggia, P. Udompholkul, E. L. Atienza, K. Muzzarelli, Z. Assar and M. Pellecchia, Covalent Targeting of Histidine Residues with Aryl Fluorosulfates: Application to Mcl-1 BH3 Mimetics, J. Med. Chem., 2024, 67, 20214–20223

23. L. H. Jones, Emerging Utility of Fluorosulfate Chemical Probes, ACS Med. Chem. Lett., 2018, 9, 584–586

24. W. Chen, J. Dong, S. Li, Y. Liu, Y. Wang, L. Yoon, P. Wu, K. B. Sharpless and J. W. Kelly, Synthesis of Sulfotyrosine-Containing Peptides by Incorporating Fluorosulfated Tyrosine Using an Fmoc Solid-phase Strategy Angew. Chem. Int. Ed., 2015, 55, 1835–1838

25. J. Tang, C. L. Yu, S. R. Williams, E. Springman, D. Jeffery, P. A. Sprengeler, A. Estevez, J. Sampang, W. Shrader, J. Spencer, W. Young, M. McGrath and B. A. Katz, Expression, Crystallization, and Three-Dimensional Structure of the Catalytic Domain of Human Plasma Kallikrein, J. Biol. Chem., 2005, 280, 41077–41089.

26. H. Zhou, P. Mukherjee, R. Liu, E. Evrard, D. Wang, J. M. Humphrey, T. W. Butler, L. R. Hoth, J. B. Sperry, S. K. Sakata, C. J. Helal and C. W. am Ende, Introduction of a Crystalline, Shelf-Stable Reagent for the Synthesis of Sulfur(VI) Fluorides, Org. Lett., 2018, 20, 812–815.

27. T. M. Postma, M. Giraud and F. Albericio, Trimethoxyphenylthio as a Highly Labile Replacement for tert-Butylthio Cysteine Protection in Fmoc Solid Phase Synthesis, Org. Lett., 2012, 14, 5468–5471

28. A. E. Doherty, C. Oxford, F. Mazumder, K. E. Gilbert, A. Aatkar, E. Mccormick, M. Kabu, P. Pogány, F. Zappacosta, C. S. Greenwood, D. J. Norman, R. E. Peltier-Heap, N. C. O. Tomkinson, G. A. Burley, E. K. Grant, D. House, C. Jamieson, J. T. Bush and J. Pettinger, Next Generation Sulfonyl Fluoride Electrophiles Expand the Scope of Covalent Drug Discovery, ChemRxiv, 2021, DOI: 10.26434/chemrxiv.15000707.v1.

29. Z. Dong, A. D. Saliganan, H. Meng, S. M. Nabha, A. L. Sabbota, S. Sheng, R. D. Bonfil and M. L. Cher, Prostate Cancer Cell-Derived Urokinase-Type Plasminogen Activator Contributes to Intraosseous Tumor Growth and Bone Turnover, Neoplasia, 2008, 10, 439–449

30. K. Dass, A. Ahmad, A. S. Azmi, S. H. Sarkar and F. H. Sarkar, Evolving Role of uPA/uPAR System in Human Cancers, Cancer Treat. Rev., 2008, 34, 122–136

31. A. Angelini, L. Cendron, S. Chen, J. Touati, G. Winter, G. Zanotti and C. Heinis, Bicyclic Peptide Inhibitor Reveals Large Contact Interface with a Protease Target, ACS Chem. Biol., 2012, 7, 817–821.

