## Supplementary-Information for "Beyond the Catalytic Serine: Selective Protease Engagement with Covalent Macrocyclic Activity-Based Probes"

### 1 Materials and Methods

#### 1.1 Chemicals and Reagents

All chemicals and reagents were purchased from commercial suppliers and used without further purification. Rink Amide resin (100-200 mesh) and 2-Chlorotrityl chloride resin (100-200 mesh, 1% DVB) were obtained from Merck. Natural Fmoc-amino acids were obtained from Fluorochem. Amicon 3 kDa and 10 kDa centrifugal filters were obtained from Merck. Zeba spin columns (7 kDa) were obtained from Thermo Fisher Scientific. Purified plasma kallikrein was obtained from Innovative Research. Human urokinase-type plasminogen activator was obtained from AcroBiosystems. Streptavidin- and neutravidin-coated magnetic beads were purchased from Thermo Fisher Scientific. 2YT and agar media were purchased from Merck. All remaining biological products and chemical reagents were obtained from Fluorochem, VWR, Sigma Aldrich, Thermo Fisher Scientific or Merck.

#### 1.2 Equipment List

DNA amplification was performed using a SimpliAmp Thermal Cycler (Fisher). Gel electrophoresis was performed using an mPAGE Mini Gel Tank (Millipore) and PowerPac™ Basic (BioRad). Fluorescent plate-based enzymatic assays were carried out using a VANTAstar Plate reader (BMG Labtech) in kinetic mode for a minimum of 5 min. In-gel fluorescence-based assays were imaged on a BioRad ChemiDoc™ Imaging System. Peptide synthesis was performed on a Multipep 2 (CEM). Flash Chromatography was performed on either a CombiFlash NextGen 300 (Teledyne ISCO) or ECS28P01 Compact Preparative System (ECOM). Sample drying was performed using a Genevac EZ-2 or Edwards Modulyo Freeze Dryer. High resolution mass spectrometry data was collected using an Agilent 1260 Infinity II coupled to a Triple Quadrupole Agilent 6545Q+TOF ESI.

### 2 Biological and Biochemical Methods

#### 2.1 Biotinylation of Human Plasma Kallikrein

Human plasma kallikrein was biotinylated using EZ-Link-Sulfo-NHS-LC-Biotin (Thermo Scientific). Briefly, 0.2 mg of plasma kallikrein (10 μM in PBS, pH 7.4) was incubated with a 10-fold molar excess of freshly prepared EZ-Link-Sulfo-NHS-LC-Biotin for 60 min at RT. Unreacted biotinylation reagent was subsequently removed via size-exclusion chromatography using a disposable Zeba spin desalting column. The final concentration of the biotinylated target was determined using a BCA protein assay.

To confirm successful biotinylation, the target protein was subjected to a magnetic bead capture assay. Biotinylated plasma kallikrein (1 μg) and non-biotinylated plasma kallikrein (1 μg) were incubated with 25 μL of streptavidin-coated magnetic beads for 60 min at RT. The bead fractions (post-boiling) and the unbound supernatant fractions for both samples were then analysed by SDS-PAGE followed by Coomassie Brilliant Blue staining. Efficient biotinylation was visually verified by the presence of the biotinylated protein exclusively in the boiled bead fraction, whereas the non-biotinylated control remained entirely in the supernatant.

#### 2.2 Phage Production

An $AXCX_7C$ peptide phage library was utilised for peptide selection against plasma kallikrein.[1] The phage particles feature a disulfide-free pIII coat protein, to prevent reaction of linker with cysteines on the pIII protein, which significantly lowers the phage infectivity. An aliquot (1 mL) of the phage peptide library in TG1 *E.coli* bacteria was thawed on ice and used to inoculate 0.5L of 2YT rich medium supplemented with 30 μg $ml^{-1}$ chloramphenicol. The culture was incubated for 16 h at 30 °C with vigorous shaking (250 rpm). Bacteria were pelleted by centrifugation at 8,500 rpm for 30 min at 4 °C and the supernatant was collected.

The supernatant was combined with a 25% (v/v) precipitation buffer (20% w/v PEG 8000, 2.5 M NaCl) and incubated for 60 min at 4 °C with gentle swirling every 15 min to maximise phage recovery. Precipitated phage particles were then pelleted by centrifugation at 9,000 rpm for 60 min at 4 °C. The supernatant was discarded and the inner walls of the tube were carefully dried to maximise PEG removal without disturbing the phage pellet. The pellet was resuspended in 10 mL of sterile PBS and centrifuged at 16,000 rpm for 10 min at 4 °C to remove residual cell debris. The cleared supernatant was transferred to a fresh tube containing 25% (v/v) of the PEG precipitation solution and incubated at 4 °C for an additional 60 min. Purified phage particles were collected by precipitation at 9,000 rpm for 60 min at 4 °C. The supernatant was discarded, and the final pellet was resuspended in a 50% (v/v) glycerol in PBS solution to a final 1 mL volume. A final purification step was performed by centrifugation at 18,000 rpm for 20 min at 4 °C in a bench-top centrifuge to remove cell debris. The purified phage stock was stored at -20 °C until further use.

To monitor phage production, serial 10-fold dilutions of the stock were prepared in PBS. Separately, a 5 mL 2YT culture from a single colony of *E.coli* TG1 cells was grown overnight and used to prepare a fresh culture in exponential growth phase ($OD_{600}$ = 0.4). To allow for phage infection, 20 µL of the highly diluted phage samples (e.g., $10^6$, $10^7$, $10^8$) were mixed with 180 µL of the exponential phase TG1 cells. The infection mixtures were incubated for 90 min at 37 °C. Subsequently, 10 µL of each mixture was spotted in triplicate onto 2YT agar plates supplemented with 30 µg $ml^{-1}$ chloramphenicol and incubated overnight at 37 °C. The following day, visible colonies were counted, and the number of phage particles per mL of solution was determined using the following formula:

$$\frac{phage}{mL} = \frac{x}{10} \times 10^{y} \times 10^{3}$$

where x is the average number of colonies across three replicates within a specific titer and y is the absolute dilution factor of the titer yielding countable colonies.

#### 2.3 On-Phage Peptide Cyclisation

Phage-displayed peptides were cyclised using 1,3-Bis(bromomethyl)benzene. First, peptide cysteines were reduced by incubating the phage library with 1 mM TCEP in 50% (v/v) glycerol-PBS for 30 min at RT. The reduced phage particles were then precipitated from the media using the previously described PEG/NaCl buffer; due to the smaller working volume, incubation and centrifugation times were reduced to 30 min. Following supernatant removal, the phage pellet was resuspended in alkylation buffer (50 mM $NH_4HCO_3$, 5 mM EDTA, pH 8.5) to facilitate cysteine alkylation. A pre-made solution of DBMB in MeCN was subsequently added to achieve a final linker concentration of 40 µM and 20% (v/v) MeCN in a total reaction volume of 1 mL. The reaction was incubated with continuous mixing for 60 min at 30 °C.

Following alkylation, a final phage precipitation was performed using PEG/NaCl as described above. The cyclised phage pellet was then resuspended in 50% (v/v) glycerol in PBS for storage at -20 °C, or, if proceeding directly to biopanning experiments, in plasma kallikrein activity buffer (10 mM Tris-Cl, 150 mM NaCl, 10 mM $MgCl_2$, 1 mM $CaCl_2$, pH 7.4) supplemented with 0.3% (v/v) Tween-20 and 3% (w/v) BSA.

#### 2.4 Phage Panning

To enrich for high-affinity peptide binders, three rounds of biopanning were performed against immobilised target protein. Biotinylated plasma kallikrein was immobilised onto either streptavidin- or neutravidin-coated magnetic beads (Cytiva). To prevent the enrichment of matrix-specific background binders, the capture matrix was alternated between rounds (e.g., neutravidin beads were utilised in the third round). To increase selection stringency for successive panning rounds, the quantity of immobilised protein was reduced from 5 µg for the first round to 2 µg for rounds two and three. Each panning round included a parallel negative control, wherein half of the input phage was panned against target-free magnetic beads to account for non-specific binding.

The day prior to biopanning, a 5 mL 2YT culture was inoculated from a single colony of *E.coli* TG1. If the phage-displayed cyclic peptide library stock was stored in 50% (v/v) glycerol, it was first precipitated using the PEG/NaCl protocol and resuspended in panning buffer (plasma kallikrein activity buffer supplemented with 0.3% (v/v) Tween-20 and 3% (w/v) BSA).

Prior to target incubation, the cyclised phage library was subjected to a pre-clearing step to deplete non-specific bead binders. A 30 µL aliquot of a 1:1 mixture of streptavidin and neutravidin magnetic beads was washed three times with 200 µL with plasma kallikrein activity buffer. The phage library was added to these washed beads and incubated for 60 min at RT with continuous mixing.

Following the phage pre-clearing step, the beads were magnetically separated and discarded, while the supernatant containing the pre-cleared phage was retained. This phage library was equally divided: one half was combined with biotinylated plasma kallikrein (2-5 µg) in panning buffer to form the positive selection mixture, while the remaining half was combined with an equal volume of panning buffer without target to serve as the negative control. Both samples were then incubated at 37 °C with slow rotation for 30 min.

Simultaneously, two 25 µL aliquots of the selected magnetic beads (streptavidin or neutravidin) were washed three times with 200 µL of activity buffer, followed by resuspension in 50 µL of panning buffer and blocking for 30 min at RT with rotation. Following the initial in-solution incubation, the pre-blocked magnetic beads were added to each library mixture and incubated for 30 min at RT with slow rotation to capture the biotinylated target complexes.

After the target capture step, the beads were isolated using a magnetic rack and washed ten times with PBS-T (PBS containing 0.1% (v/v) Tween-20). To increase selection stringency, the duration of the individual wash steps was progressively extended across successive panning rounds (See Table 1).

***Supplementary Table 1: Washing steps in the phage panning experiment***

| Panning Round | PBS-T |
|---|---|
| **1st round** | 10 x 2 min |
| **2nd round** | 4 x 2 min, 2 x 10 min, 4 x 2 min |
| **3rd round** | 3 x 2 min, 4 x 20 min, 3 x 2 min |

To minimise the retention of non-specific, plastic-binding phage, the samples were transferred to fresh microcentrifuge tubes after every two washes. After the final wash, the buffer was removed, the beads resuspended in 100 μL elution buffer (50 mM Glycine, pH 2.2) and rotated at RT for 10 min. The beads were then captured and the eluted phage supernatants transferred to new tubes containing 50 μL of neutralisation buffer (1 M Tris-Cl, pH 8.0). A 10 μL aliquot from both the target and negative control panning outputs was reserved for phage titration. The remaining eluted phage from both conditions were used to inoculate two separate 50 mL cultures of *E.coli* TG1 cells in the exponential growth phase ($OD_{600}$ = 0.4) in 2YT media. Following a 90 min infection period at 37 °C, the cells were pelleted by centrifugation at 4,000 rpm for 10 min at 4 °C. The cell pellets were resuspended in 1 mL of 2YT media and plated onto two large 2YT agar plates supplemented with chloramphenicol and incubated overnight at 37 °C. The following day, the amplified cells were harvested by scraping the plates with 2 mL of 2YT media per plate. Glycerol was added to a final concentration of 50% (v/v), and 1 mL aliquots were stored at -80 °C. To initiate the next round of selection, input phage particles were produced from these glycerol stocks following the procedure described in **Section 2.2**. The enrichment factor for each panning round was determined by calculating the ratio of the phage titre recovered from the target (plasma kallikrein) output against the titre from the negative control output.

#### 2.5 Sample Preparation for Next-Generation Sequencing

Following three rounds of biopanning, DNA from both the phage enriched against plasma kallikrein and the negative control was prepared for Next-Generation Sequencing (NGS) analysis. All samples were processed in triplicate. A two-step PCR amplification strategy was employed: the primary PCR utilised a conserved set of five forward and five reverse primers to amplify the peptide-encoding region, while the secondary PCR step utilised unique combinations of forward and reverse primers to append sample-specific indices (barcodes) for multiplexed sequencing analysis (See **Supplementary Table 2**).

***Supplementary Table 2: List of primers used in NGS sample preparation***

| Round | **Fwd/Rvr** | **Primer** | **Sequence** |
|---|---|---|---|
| **1** | **Fwd.** | 1a | TCGTCGGCAGCGTCAGATGTGTATAAGAGACAGCCAGAGCCACCCTCGCTACCG |
| | | 1b | TCGTCGGCAGCGTCAGATGTGTATAAGAGACAG**N**CCAGAGCCACCCTCGCTACCG |
| | | 1c | TCGTCGGCAGCGTCAGATGTGTATAAGAGACAG**NN**CCAGAGCCACCCTCGCTACCG |
| | | 1d | TCGTCGGCAGCGTCAGATGTGTATAAGAGACAG**NNN**CCAGAGCCACCCTCGCTACCG |
| | | 1e | TCGTCGGCAGCGTCAGATGTGTATAAGAGACAG**NNNN**CCAGAGCCACCCTCGCTACCG |
| | **Rvrs.** | 2a | GTCTCGTGGGCTCGGAGATGTGTATAAGAGACAGGCTATGCGGCCCAGCCGGCC |
| | | 2b | GTCTCGTGGGCTCGGAGATGTGTATAAGAGACAGG**N**CTATGCGGCCCAGCCGGCC |
| | | 2c | GTCTCGTGGGCTCGGAGATGTGTATAAGAGACAGG**NN**CTATGCGGCCCAGCCGGCC |
| | | 2d | GTCTCGTGGGCTCGGAGATGTGTATAAGAGACAGG**NNN**CTATGCGGCCCAGCCGGCC |
| | | 2e | GTCTCGTGGGCTCGGAGATGTGTATAAGAGACAGG**NNNN**CTATGCGGCCCAGCCGGCC |
| **2** | **Fwd.** | S505 | AATGATACGGCGACCACCGAGATCTACAC**GTAAGGA**GTCGTCGGCAGCGTC |
| | | S506 | AATGATACGGCGACCACCGAGATCTACAC**ACTGCATA**TCGTCGGCAGCGTC |
| | | S507 | AATGATACGGCGACCACCGAGATCTACAC**AAGGAGTA**TCGTCGGCAGCGTC |
| | **Rvrs.** | N704 | CAAGCAGAAGACGGCATACGAGAT**GCTCAGGA**GTCTCGTGGGCTCGG |
| | | N705 | CAAGCAGAAGACGGCATACGAGAT**AGGAGTCC**GTCTCGTGGGCTCGG |

To isolate template DNA for the primary PCR, three 200 μL aliquots were sampled from the third-round plasma kallikrein panning output glycerol stock and the corresponding third-round negative control glycerol stock. Plasmid DNA was purified from these aliquots using the Monarch Plasmid Miniprep Kit (New England Biolabs) according to the manufacturer's instructions. For each subsequent primary PCR amplification, 200 ng of the purified plasmid DNA was utilised as the template (See **Supplementary Table 3** and **4**).

***Supplementary Table 3: Reagents used in primary PCR for NGS sample preparation.***

| **Reagent** | **Volume** |
|---|---|
| 1a<br>1b<br>1c<br>1d<br>1e | 1 μL<br>1 μL<br>1 μL<br>1 μL<br>1 μL |
| 2a<br>2b<br>2c<br>2d<br>2e | 1 μL<br>1 μL<br>1 μL<br>1 μL<br>1 μL |
| dNTP mix | 1 μL |
| Plasmid DNA | 200 ng |
| 5X HF buffer | 10 μL |
| Phusion Enzyme | 2 μL |
| $dH_2O$ | Made up to 50 μL |

***Supplementary Table 4: Conditions used in primary PCR for NGS sample preparation.***

| Temperature (°C) | Time | Cycles |
|---|---|---|
| 95 | 5 min | 1 |
| 95 | 30 sec | 25 |
| 72 | 45 sec | |
| 72 | 1 min | |
| 72 | 5 min | 1 |
| 4 | ∞ | |

The unpurified amplicon from the primary PCR was utilised directly as the template for the secondary indexing PCR, which was assembled as follows (See Table 5, 6, 7).

***Supplementary Table 5: Reagents used in secondary PCR for NGS sample preparation.***

| Reagent | Volume |
|---|---|
| Fwd. | 1 µL |
| Rvrs. | 1 µL |
| dNTP mix | 1 µL |
| PCR 1 DNA | 2 µL |
| 5X HF buffer | 10 µL |
| Phusion Enzyme | 2 µL |
| $dH_2O$ | Made up to 50 µL |

***Supplementary Table 6: Conditions used in secondary PCR for NGS sample preparation.***

| **Temperature (°C)** | **Time** | **Cycles** |
|---|---|---|
| 95 | 5 min | 1 |
| 95 | 30 sec | 25 |
| 61 | 45 sec | |
| 72 | 1 min | |
| 72 | 5 min | 1 |
| 4 | ∞ | |

***Supplementary Table 7: Specific barcoded-primers used for each NGS sample analysed.***

| **Fwd.** | **Rvrs.** | **Sample** |
|---|---|---|
| S505 | N704 | Control-1 |
| S506 | N704 | Control -2 |
| S507 | N704 | Control -3 |
| S505 | N705 | Protein-1 |
| S506 | N705 | Protein -2 |
| S507 | N705 | Protein-3 |

The secondary PCR products were resolved by electrophoresis on a 2% (w/v) agarose gel in Tris-Borate-EDTA (TBE) buffer at 100 V for 30 min, utilising a Low Molecular Weight DNA Ladder (New England Biolabs) as a reference standard. Amplicons corresponding to the expected size of 270 bp were excised from the gel and purified using the Monarch DNA Gel Extraction Kit (New England Biolabs) according to the manufacturer's instructions. Finally, equimolar amounts of the purified, barcoded amplicons were then pooled together to generate the final library for Next Generation Sequencing (NGS) analysis.

#### 2.6 Next-Generation Sequencing Analysis

Raw sequencing data were first processed to ensure high read quality. Sequences containing excessive errors, ambiguous bases, or lacking the correct peptide sequence were discarded. The remaining high-quality peptide sequences were arranged as count matrices for downstream statistical analysis.

To identify peptides specifically enriched in the plasma kallikrein samples compared to the control group, differential enrichment analysis was conducted using the DESeq2 package in R. Count data were normalised, and statistical significance was calculated using the default DESeq2 pipeline. Peptides were defined as significantly enriched if they demonstrated a $log_2(Fold\ Change) > 1.5$ and an adjusted p-value $\left(p_{adj}\right) < 0.05$.

The significantly enriched peptides identified via DESeq2 were clustered based on sequence similarity using the Gibbs clustering algorithm (GibbsCluster version 2.0). This grouped the related peptide families and determined consensus sequence motifs.

Custom scripts utilised for sequence filtering, quality control, and data analysis are available from the corresponding author upon request.

#### 2.7 Enzyme Assays

The inhibitory activity of the cyclic peptides was evaluated by incubating the target proteases with varying concentrations of the inhibitors and subsequently quantifying residual enzyme activity using specific fluorogenic substrates. Briefly, the peptide inhibitors were pre-incubated with the respective target protein in assay buffer at 37 °C for 30 min. Following the addition of the substrate, enzyme activity was determined by measuring fluorescence intensity at 1 min intervals over a 30 min period. The inhibitory constant ($K_i$) was calculated according to the Cheng–Prusoff equation: $K_i$=IC50/(1+([S]0/$K_M$)), where IC50 is the functional strength of the inhibitor, [S]0 is the total substrate concentration, and $K_M$ is the Michaelis–Menten constant.[2]

**Plasma Kallikrein Assay conditions**

- **Buffer:** 50 mM Tris-HCl (pH 7.4), 150 mM NaCl, 10 mM $MgCl_2$, 1 mM $CaCl_2$, 0.01% (w/v) BSA, 0.01% (v/v) Triton X-100, 5% (v/v) DMSO.
- **Enzyme:** Purified Human Plasma Kallikrein (Innovative Research), 10 nM
- **Substrate:** Cbz-Phe-Arg-AMC (Thermo Fisher Scientific), 50 μM

**Urokinase-Type Plasminogen Activator (uPA) Assay conditions**

- **Buffer:** 50 mM Tris-HCl (pH 7.4), 150 mM NaCl, 10 mM $MgCl_2$, 1 mM $CaCl_2$, 0.01% (w/v) BSA, 0.01% (v/v) Triton X-100, 5% (v/v) DMSO.
- **Enzyme:** Human PLAU/uPA protein (AcroBiostems), 1.5 nM
- **Substrate:** Cbz-Gly-Gly-Arg-AMC (SantaCruz Biotechnologies), 50 μM.

**Trypsin Assay conditions**

- **Buffer:** 50 mM ammonium bicarbonate (pH 8.0).
- **Enzyme:** Sequencing Grade Modified Trypsin (Promega), 50 nM.
- **Substrate:** Cbz-Phe-Arg-AMC, 50 μM.

**Thrombin Assay conditions**

- **Buffer:** 50 mM Tris-HCl (pH 7.5), 10 mM $CaCl_2$, 150 mM NaCl, 0.05% (w/v) Brij-35.
- **Enzyme:** Recombinant Human Coagulation Factor II/Thrombin (R&D Systems), 0.6 nM.
- **Substrate:** BOC-VPR-AMC, 50 µM.

**Plasmin Assay conditions**

- **Buffer:** 0.1 M Tris-HCl (pH 7.5), 0.1 M NaCl.
- **Enzyme:** Human Plasminogen Protein (R&D Systems), 6 nM.
- **Substrate:** SUC-Ala-Phe-Lys-AMC, 50 µM.

**KLK2 Assay conditions**

- **Buffer:** 50 mM Tris-HCl (pH 7.5), 150 mM NaCl, 10 mM $CaCl_2$, 0.05% (w/v) Brij-35.
- **Enzyme:** Recombinant Human Kallikrein 2 (AcroBiosystems),12 nM.
- **Substrate:** Morpholin-Val-Dab-4Pal-Arg-ACC, 10 µM.

**KLK14 Assay conditions**

- **Buffer:** 50 mM Tris-HCl (pH 8.0), 150 mM NaCl, 0.05% (w/v) Brij-35.
- **Enzyme:** Recombinant Human Kallikrein 14 (AcroBiosystems), 4 nM.
- **Substrate:** BOC-VPR-AMC, 100 µM.

#### 2.8 In-gel Fluorescence

The concentrations of the target proteins and covalent probes, as well as the duration of the incubation periods, were optimised for each specific target-probe pair. Control samples were treated with an equivalent volume of DMSO. All primary incubations were performed in PBS (pH 7.4) at 37 °C and in a final reaction volume of 25 µL.

Following the initial incubation, the samples were subjected to copper-catalysed azide-alkyne cycloaddition (CuAAC) for 60 min via the addition of 2.5 µL of a freshly prepared 'click' chemistry cocktail. The final concentrations in the conjugation reactions were 100 μM TAMRA-azide, 1 mM $CuSO_4$, 1 mM TCEP and 100 μM TBTA (Tris((1-benzyl-4-triazolyl)methyl)amine). The reactions were subsequently quenched with 4X SDS Laemmli Loading Buffer with 5% β-Mercaptoethanol and denatured by heating at 95 °C for 10 min. Proteins were resolved via SDS-PAGE, and in-gel TAMRA fluorescence was visualised using the Cy3 excitation/emission channels on a Gel Doc Imaging System (Bio-Rad).

#### 2.9 Intact Protein Mass Spectrometry

To facilitate intact mass analysis, a non-glycosylated variant of the plasma kallikrein protease domain was utilised. GenScript expressed this protein in Sf9 cells followed by purification using a HisTrap column. Specifically, to prevent heterogeneous glycosylation and eliminate unpaired thiols, the native sequence was engineered with the following substitutions: three N-linked glycosylation sites were mutated to glutamic acid (N377E, N434E, and N475E, highlighted in blue), and two specific cysteine residues were mutated to serine (C364S and C484S, highlighted in red).

$^{357}$NTGDNSVSTTKTSTRIVGGTESSWGEWPWQVSLQVKLTAQRHLCGGSLIGHQWVLTAAH
CFDGLPLQDVWRIYSGILELSDITKDTPFSQIKEIIIHQNYKVSEGNHDIALIKLQAPLEYTEFQK
PISLPSKGDTSTIYTNCWVTGWGFSKEKGEIQNILQKVNIPLVTNEECQKRYQDYKITQRMVC
AGYKEGGKDACKGDSGGPLVCKHNGMWRLVGITSWGEGCARREQPGVYTKVAEYMDWIL
EK$^{612}$TQSSDGKAQMQSPA$^{622}$HHHHHH

To evaluate covalent-adduct formation, the engineered non-glycosylated plasma kallikrein (5 μM) was incubated with a 10-fold molar excess of covalent probe in assay buffer (10 mM Tris-HCl, 150 mM NaCl, pH 7.4). The reactions were incubated at 37 °C with continuous rotation for varying time intervals. The extent of covalent modification was analysed by intact-protein mass spectrometry utilising an Agilent Q-TOF LC/MS instrument. Data processing was performed by biomolecular deconvolution using Agilent MassHunter BioConfirm Version 10.0.

#### 2.10 Expression of Plasma Kallikrein Mutants

Target nucleophilic residues for mutagenesis were selected based on their spatial proximity to the active site of the plasma kallikrein protease domain, as determined from the published crystal structure (PDB: 2ANY). The corresponding mutant constructs (mutant residue highlighted in green) were expressed and purified by GenScript using the TurboCHO platform and HisTrap columns, respectively.

**H40N**

$^{357}$NTGDNSVSTTKTSTRIVGGTESSWGEWPWQVSLQVKLTAQR**N**LCGGSLIGHQWVLTAAH
CFDGLPLQDVWRIYSGILELSDITKDTPFSQIKEIIIHQNYKVSEGNHDIALIKLQAPLEYTEFQK
PISLPSKGDTSTIYTNCWVTGWGFSKEKGEIQNILQKVNIPLVTNEECQKRYQDYKITQRMVCA
GYKEGGKDACKGDSGGPLVCKHNGMWRLVGITSWGEGCARREQPGVYTKVAEYMDWILE
K$^{612}$TQSSDGKAQMQSPA$^{622}$HHHHHH

**Y94F**

$^{357}$NTGDNSVSTTKTSTRIVGGTESSWGEWPWQVSLQVKLTAQRHLCGGSLIGHQWVLTAAH
CFDGLPLQDVWRIYSGILELSDITKDTPFSQIKEIIIHQN**F**KVSEGNHDIALIKLQAPLEYTEFQK
PISLPSKGDTSTIYTNCWVTGWGFSKEKGEIQNILQKVNIPLVTNEECQKRYQDYKITQRMVCA
GYKEGGKDACKGDSGGPLVCKHNGMWRLVGITSWGEGCARREQPGVYTKVAEYMDWILE
K$^{612}$TQSSDGKAQMQSPA$^{622}$HHHHHH

**K147R**

$^{357}$NTGDNSVSTTKTSTRIVGGTESSWGEWPWQVSLQVKLTAQRHLCGGSLIGHQWVLTAAH
CFDGLPLQDVWRIYSGILELSDITKDTPFSQIKEIIIHQEYKVSEGNHDIALIKLQAPLEYTEFQK
PISLPSKGDTSTIYTNCWVTGWGFSKE**R**GEIQNILQKVNIPLVTNEECQKRYQDYKITQRMVC
AGYKEGGKDACKGDSGGPLVCKHNGMWRLVGITSWGEGCARREQPGVYTKVAEYMDWIL
EKTQSSDGKAQMQSPA$^{622}$HHHHHH

**Y172F**

$^{357}$NTGDNSVSTTKTSTRIVGGTESSWGEWPWQVSLQVKLTAQRHLCGGSLIGHQWVLTAAH
CFDGLPLQDVWRIYSGILELSDITKDTPFSQIKEIIIHQEYKVSEGNHDIALIKLQAPLEYTEFQK
PISLPSKGDTSTIYTNCWVTGWGFSKEKGEIQNILQKVNIPLVTNEECQKR**F**QDYKITQRMVC
AGYKEGGKDACKGDSGGPLVCKHNGMWRLVGITSWGEGCARREQPGVYTKVAEYMDWIL
EKTQSSDGKAQMQSPA$^{622}$HHHHHH

**Y175F**

$^{357}$NTGDNSVSTTKTSTRI$^{372}$VGGTESSWGEWPWQVSLQVKLTAQRHLCGGSLIGHQWVLTAA
HCFDGLPLQDVWRIYSGILELSDITKDTPFSQIKEIIIHQEYKVSEGNHDIALIKLQAPLEYTEFQ
KPISLPSKGDTSTIYTNCWVTGWGFSKEKGEIQNILQKVNIPLVTNEECQKRYQD**F**KITQRMVC
AGYKEGGKDACKGDSGGPLVCKHNGMWRLVGITSWGEGCARREQPGVYTKVAEYMDWIL
EK$^{622}$TQSSDGKAQMQSPAHHHHHH

**K192R**

$^{357}$NTGDNSVSTTKTSTRIVGGTESSWGEWPWQVSLQVKLTAQRHLCGGSLIGHQWVLTAAH
CFDGLPLQDVWRIYSGILELSDITKDTPFSQIKEIIIHQEYKVSEGNHDIALIKLQAPLEYTEFQK
PISLPSKGDTSTIYTNCWVTGWGFSKEKGEIQNILQKVNIPLVTNEECQKRYQDYKITQRMVCA
GYKEGGKDAC**R**GDSGGPLVCKHNGMWRLVGITSWGEGCARREQPGVYTKVAEYMDWILE
KTQSSDGKAQMQSPA$^{622}$HHHHHH

#### 2.11 Tryptic Digest LC–MS/MS Analysis

**Experimental:** 1 µM plasma kallikrein was incubated with 1 µM **23** or DMSO in 10 mM Tris, 150 mM NaCl, 10 mM MgCl2, 1 mM $CaCl_2$, pH 7.4 for 4 h at 37 °C. The samples were reduced (10 mM TCEP, 55°C for 1h), alkylated (40 mM iodoacetamide, room temperature for 1 h), reduced again (40 mM DTT, room temperature for 1 h) and then digested overnight at 37 °C using 0.5 µg sequencing grade trypsin (Promega). The resulting peptides were desalted using C18 ziptips according to the manufacturer's instructions (Millipore). Eluate from the ziptips was evaporated to dryness, resuspended in 1% formic acid and the peptides analysed using an Ultimate 3000 nano-LC system in line with an Orbitrap Fusion Tribrid mass spectrometer (Thermo Scientific). In brief, peptides in 1% (vol/vol) formic acid were injected onto an Acclaim

PepMap C18 nano-trap column (Thermo Scientific). After washing with 0.5% (vol/vol) acetonitrile 0.1% (vol/vol) formic acid peptides were resolved on a 500 mm × 75 µm Acclaim PepMap C18 reverse phase analytical column (Thermo Scientific) over an 80 min organic gradient, using 4 gradient segments (1-50% solvent B over 55 min., 50-90% B over 0.5 min., held at 90% B for 4.5 min and then reduced to 1% B over 0.5 min.) with a flow rate of 300 nL min−1. Solvent A was 0.1% formic acid and Solvent B was aqueous 80% acetonitrile in 0.1% formic acid. Peptides were ionized by nano-electrospray ionization at 2.2 kV using a stainless-steel emitter with an internal diameter of 30 µm (Thermo Scientific) and a capillary temperature of 275 °C.

All spectra were acquired using an Orbitrap Fusion Tribrid mass spectrometer controlled by Xcalibur 2.1 software (Thermo Scientific) and operated in data-dependent acquisition mode. FTMS1 spectra were collected at a resolution of 120 000 over a scan range (m/z) of 400-2000, with an automatic gain control (AGC) target of 400 000 and a max injection time of 50ms. Precursors were filtered according to charge state (to include charge states 2-7), with monoisotopic peak determination set to peptide and using an intensity range from 5E3 to 1E20. Previously interrogated precursors were excluded using a dynamic window (30s +/-10ppm). The MS2 precursors were isolated with a quadrupole mass filter set to a width of 1.6m/z. ITMS2 spectra were collected with an AGC target of 5000, max injection time of 50ms and HCD collision energy of 35%.

**Data Analysis:** Proteomics data analysis was performed using Fragpipe (v23.1) with MSFragger v4.4.1, IonQuant v1.11.20, diaTracer v2.2.1. For analysis, the "Basic Search" workflow was used with the following modification: Fragment mass tolerance 0.5 Da, Cleavage by Trypsin (KR, but not P), Peptide length 6-50 and mass range 500-5000 Da, fixed modification C=57.02146 (Carbamidomethylation: CAM). For PSM Validation, Percolator was run with the following settings: Min probability 0.5, opts: --only-psms --no-terminate --post-processing-tdc. ProteinProphet was run using the following opts: --maxppmdiff 2000000. MS1 quantification was performed using default settings with a "Min site localization probability" of 0.25.

Variable modification was set for STYKH to the mass of the expected **23** adduct (1540.6050 Da) and its single (1556.5999 Da)- and double (1572.59483 Da)-oxidized forms to account for potential thioether-oxidation. Only the search against the single-oxidized form (1556.5999 Da) resulted in the identification of modified peptides and was thus used for final analysis and PTM Site Localization using PTM Prophet with the following opts: NOSTACK KEEPOLD STATIC FRAGPPMTOL=500 EM=2 NIONS=by MINPROB=0.5 M:15.9949, n:42.0106, S:1556.5999, T:1556.5999,Y :1556.5999, K:1556.5999, H:1556.5999 (Fig. S4).

The data table 'Combined_modified_peptide.tsv' was used for further analysis of the identification of the modified peptide. Contaminants were removed and peptides quantified using intensity (for non-modified peptides) or spectral count (for modified peptides). Due to low probability scores, the exact modified amino acid within the peptides could not be unambiguously assigned.

For further validation, the HPLC-MS-Traces for **23**-modified and **23**-non-modified peptides (DACKGDSGGPLVCK and EGGKDACKGDSGGPLVCK) were analysed using FreeStyle (Thermo Scientfic) and plotted in Graphpad Prism (Fig. S5).

#### 2.12 PC3 Cell Culture

PC3 cells were seeded into a T175 culture flask and cultured in Gibco™ RPMI 1640 Medium supplemented with 10% foetal bovine serum (FBS) and 1% penicillin/streptomycin. Cells were maintained at 37 °C in humidified atmosphere of 5% $CO_2$ until reaching approximately 80% confluency. At 80% confluency, the growth medium was aspirated, and the cell monolayer was washed three times with PBS (pH 7.4) to remove residual serum proteins. The cells were incubated in 15 mL of serum-free Gibco™ RPMI 1640 Medium (without phenol red, supplemented with 1% penicillin/streptomycin) for 24 h at 37 °C, 5% $CO_2$. Following incubation, the conditioned medium was collected and centrifuged at 300 x *g* for 10 min at 4 °C to remove detached cells and debris. The cleared supernatant was collected in 1 mL aliquots, snap-frozen, and stored at -70 °C until downstream use.

#### 2.13 Plasma Kallikrein Detection in Human Plasma

Human plasma was obtained from a single healthy donor using BD vacutainer sodium citrate tubes. Plasma was obtained following centrifugation of blood tubes at 1500 x g for 10 min at 4 °C. Plasma was then aliquoted and stored at -70 °C. The volunteer provided written informed consent, and all methods and experimental protocols were conducted in accordance with and approved by the NHS Research Ethics Committee (REC permit number 25/NI/0035). Prior to incubation with the covalent probes, the plasma was diluted 1:10 in PBS (pH 7.4).

Reaction mixtures (25 µL) were prepared using the diluted plasma, which was spiked with 1 µM blood-purified human plasma kallikrein and 1 µM **23** or FP-TAMRA. To induce contact pathway activation, 2 µg of kaolin (Thermo Fisher Scientific) was added to the plasma samples. The mixtures were incubated at 37 °C for varying time intervals. Following incubation, the samples were conjugated with TAMRA-azide using the CuAAC protocol described above, and protein labelling assessed by in-gel fluorescence.

#### 2.14 cmABP Stability Tests

The hydrolytic stability of cmABPs was evaluated in phosphate-buffered saline (PBS, pH 7.4) at 37 °C. Each covalent peptide was prepared at a final concentration of 250 µM in the presence of Fmoc-Tyr(OtBu)-OH (250 µM), a control which served as an internal standard for quantitative LC-MS analysis.

To capture the degradation kinetics, incubation times were sampled at 0, 4, 8, 12, 16, 24 and 48 h.

***Supplementary Table 8: Timetable for probe stability analysis by LC-MS.***

| **Time (min)** | **Solvent A, %** <br> **($H_2O$ + 0.1 FA)** | **Solvent B %** <br> **(MeCN + 0.1 FA)** |
|---|---|---|
| 0.00 | 95 | 5 |
| 10.00 | 95 | 5 |
| 100.00 | 5 | 95 |

| | | |
|---|---|---|
| 105.00 | 5 | 95 |
| 110.00 | 95 | 5 |
| 120.00 | 95 | 5 |

The relative quantity of the remaining intact peptide at each timepoint was determined by integrating the area under the curve (AUC) of the UV signal at 254 nm and normalising it against the AUC of the internal standard. Data processing was performed using Agilent OpenLab CDS Data Analysis Software Version 2.8. To calculate the hydrolytic half-life ($t_{1/2}$), the normalised percentage of remaining intact peptide was plotted against time and fitted to a one-phase exponential decay model using GraphPad Prism.

### 3 Chemical Synthesis

#### 3.1 Chemical Reagents

***Supplementary Table 9: Unnatural amino acids for synthesis of cmABPs.***

| Cmp. | Amino Acid | Structure | Supplier |
|---|---|---|---|
| **17** | Fmoc-*m*-fluoro-L-tyrosine | 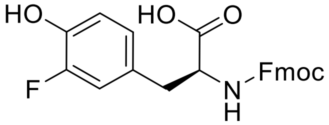 | Merck |
| **18** | Fmoc-3,5-difluoro-L-tyrosine | 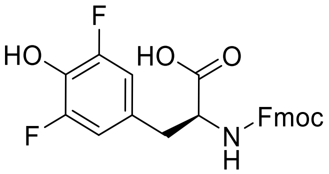 | MedChem Express |
| **19** | Fmoc-3-nitro-L-tyrosine | 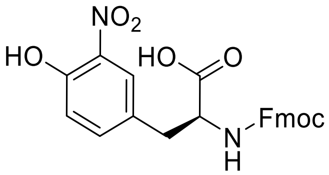 | Fluorochem |
| **20** | Fmoc-4-hydroxyphenylglycine | 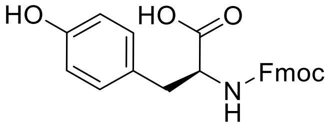 | Merck |
| **21** | Fmoc-5-hydroxy-L-tryptophan | 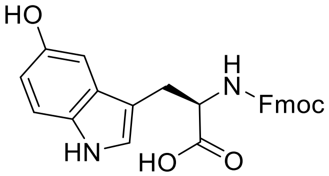 | Merck |

| | | | |
|---|---|---|---|
| **22** | Fmoc-L-homotyrosine | 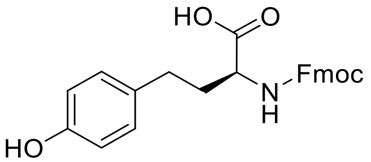 | Santa Cruz. Biotech. |

Fmoc-Dap(Mtt)-OH (Fluorochem), Fmoc-Cys(STmp)-OH (Merck), 5-(fluorosulfonyl)-2-methoxybenzoic acid (Enamine Ltd), AISF (Merck)

#### 3.2 General Procedures

##### 3.2.1 Automated Peptide Synthesis:

Peptide synthesis was performed on a 0.1 mM scale by a Multi Pep 2 peptide synthesizer (CEM) utilising Fmoc SPPS. 150 mg of rink amide resin was pre-swelled in DCM prior to synthesis. The automated double coupling method followed repeated cycles of the following steps. Fmoc deprotection: incubation with 20% 4-methylpiperidine in DMF (1 x 5 min, then 1 x 7 min). Washing: DMF (x 7) after each deprotection. Amino acid double coupling – incubation with amino acid (0.4 mmol, 4 equiv.), HBTU (0.5 mmol, 5 equiv.) and DIPEA (0.5 mmol, 5 equiv.) in DMF (2 x 30 min), RT with shaking (350 rpm). Washing: DMF (x 3) after each amino acid double coupling. Cycles were repeated until the peptide chain was complete, followed by a final Fmoc deprotection and a final wash with ethanol.

##### 3.2.2 Flash Chromatography:

Purification of the crude peptides was carried out using a CombiFlash NextGen 300 flash chromatography system (Teledyne ISCO) or ECS28P01 Compact Preparative System (ECOM). Separations were achieved utilising reversed-phase C18 columns (Teledyne ISCO) of varying capacities (5 g, 15 g, or 30 g), selected according to the scale of the synthesis. For closely eluting products, a ACCQ Prep 150 (Teledyne ISCO) was used with a RediSep Prep C18 column for >30 mg product or an Agilent Eclipse XDB-C18 column for <30 mg product. For all systems, the mobile phase consisted of Milli-Q $H_2O$ (Solvent A) and HPLC-grade MeCN (Solvent B), both supplemented with 0.1% (v/v) formic acid (FA). Peptides were eluted using a standardised 25-min linear gradient from 5% to 95% MeCN. Flow rates were scaled appropriately based on column dimensions following the manufacturer's instructions. Fraction collection was guided by UV absorbance monitoring at 220 nm.

##### 3.2.3 Analytical LC-MS:

Analysis of cmABPs was carried out on an Agilent 1260 Affinity II LC/MS coupled to a single quadrupole Agilent InfinityLab LC/MSD iQ (ESI SQ) using the following elution method: gradient of $H_2O$ and MeCN, supplemented with 0.1% formic acid: 0-10 min 5-95% MeCN with a Jupiter 4 μm Proteo 90 Å, LC C18 column (150 x 4.6 mm). A flow rate of 1 mL/min was used.

##### 3.2.4 Mass-Spectrometry:

High-resolution mass spectrometry was conducted using an Agilent 1260 Infinity II Quat pump HPLC coupled to a triple quadrupole Agilent 6545Q-TOF mass-spectrometer (ESI).

##### 3.2.5 Sample Drying:

Purified peptide fractions were concentrated using a Genevac EZ-2 series centrifugal evaporator The concentrated fractions were subsequently pooled, dissolved in a mixture of MeCN and $H_2O$, and lyophilised to a dry powder using an Edwards Modulyo freeze-dryer.

#### 3.3 Chemical Procedures

##### 3.3.1 Installation of Fluorosulfate Using Fluorosulfate Tyrosine.

For **Compounds 9-16** and **30-33**, SPPS was performed (general procedure in **Section 3.2**). After incorporating Fmoc-fluorosulfate tyrosine, removal of Fmoc groups was achieved by resin treatment with 20% (v/v) 2-methylpiperidine (3 x 5 mL) in DMF for 3 min per treatment.

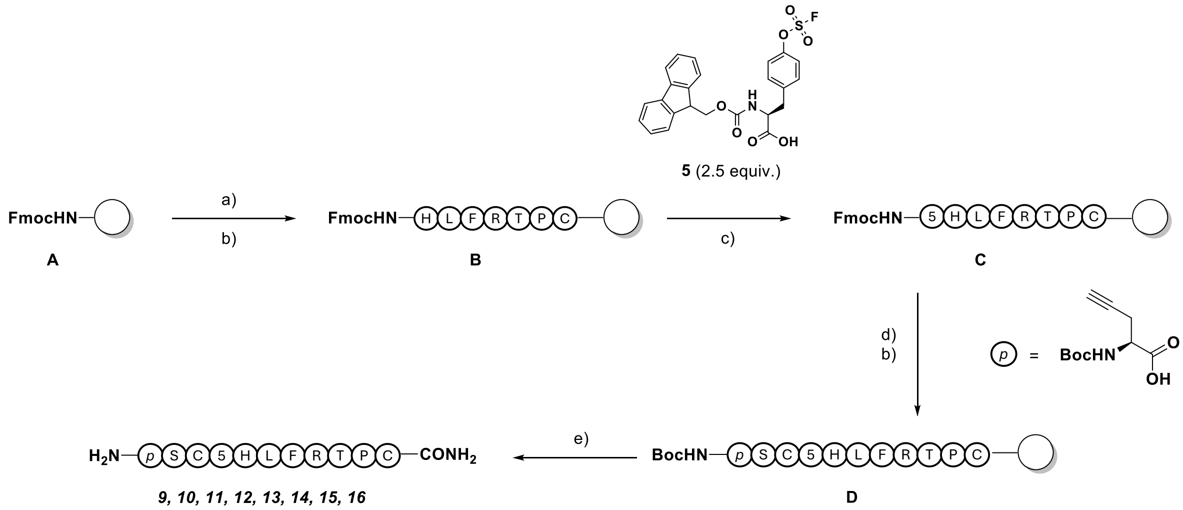

***Supplementary Scheme 1: Solid-phase synthesis of compounds 9-16.*** *a) 20% 4-methyl piperidine, DMF, 3 x 3 min, RT. b) Fmoc-Xaa-OH (5 equiv.), HBTU (5 equiv.), DIPEA (10 equiv.), 1 h, RT. c) Fmoc-5 (2.5 equiv.), HBTU (2.5 equiv.), DIPEA, (5 equiv.), DMF, 90 min, RT. d) 20% 2-methyl piperidine, DMF, 3 x 3 min, RT. e) TFA (82.5%), $H_2O$ (5%), thioanisole (5%), phenol (5%) and DODt (2.5%), 2 h, RT.*

#### 3.3.2 Synthesis of Fmoc-Tyrosine Fluorosulfate

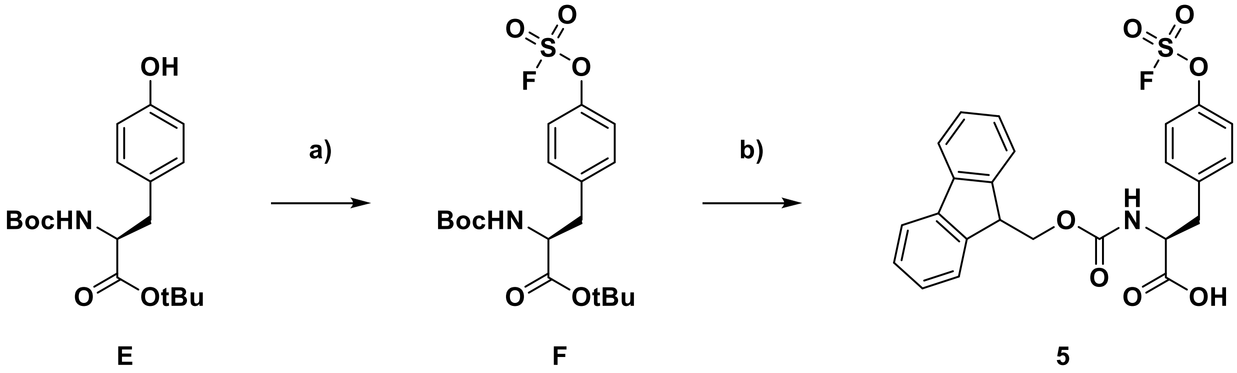

***Supplementary Scheme 2: Synthesis of Fmoc-fluorosulfate tyrosine (5).*** Boc-Tyr(OtBu)-OH, AISF, DBU, THF, RT, 15min. b) i. 50% TFA in DCM, RT, 1 h. ii. Fmoc-succinimide, 1,4-dioxane/$H_2O$, RT, 1 h.

#### tert-Butyl(S)-2-((tert-butoxycarbonyl)amino)-3-(4-((fluorosulfonyl) oxy)phenyl)propanoate (Intermediate F)

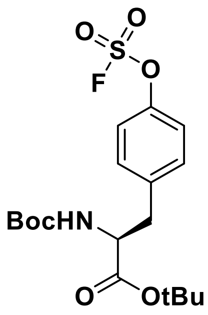

**F**

To a solution of *tert*-butyl (*tert*-butoxycarbonyl)-L-tyrosinate (**Intermediate E, Supplementary Scheme 2**) (1.79 g, 5.30 mmol, 1.0 equiv.) and AISF (2.00 g, 6.36 mmol, 1.2 equiv.) in THF (25 mL) was added 1,8-Diazabicyclo[5.4.0]undec-7-ene (1.74 mL, 11.70 mmol, 2.2 equiv.) (dropwise over 30 sec). Following complete addition, the reaction mixture was stirred for a further 15 min at RT before being diluted with 100 mL EtOAc and washed sequentially with 20 mL of 0.5 M HCl (x 2) and brine (x 1). The organic layer was dried ($MgSO_4$), filtered, and concentrated *in vacuo*. Purification *via* flash column chromatography on silica gel in petroleum ether/EtOAc using a gradient of 0–30% over 15 min followed by a gradient of 30–70% over 10 min afforded **Intermediate F**, as a white solid (1.60 g, 70%).

**$^1$H NMR** (400 MHz, DMSO-$d_6$) δ 7.55 – 7.39 (m, 4H), 7.25 (d, $J$= 8.1 Hz, 1H), 4.07 (td, $J$ = 9.0, 5.9 Hz, 1H), 3.01 (dd, $J$ = 13.8, 5.8 Hz, 1H), 2.91 (dd, $J$ = 13.7, 9.8 Hz, 1H), 1.34 (s, 18H).

#### tert-Butyl 2-((((9H-fluoren-9-yl)methoxy)carbonyl)amino)-3-(4-((fluorosulfonyl)oxy)phenyl)propanoate (5)

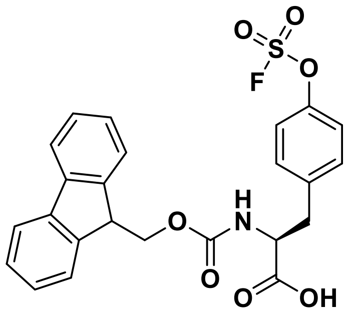

**5**

**Intermediate F** (1.12 g, 4.26 mmol, 1.0 equiv.) was dissolved in a 1:1 (v/v) mixture of DCM/TFA (20 mL) and stirred for 1 h at RT. Upon complete deprotection (monitored by TLC analysis), the reaction mixture was concentrated *in vacuo* and the resulting residue dissolved in 1:1 (v/v) 1,4-dioxane/$H_2O$ (20 mL). The pH was adjusted to 8.5 using $NaHCO_3$ and then Fmoc-OSu (1.43 g, 4.26 mmol, 1.0 equiv.) was added and stirred at RT for 1 h. The reaction mixture was acidified to pH 2 with 1 M HCl and extracted with 50 mL EtOAc (x3). The combined organic layers were washed with 20 mL of 1 M HCl (x 2) and brine (x 1), dried ($MgSO_4$), filtered, and concentrated *in vacuo*. Purification *via* flash column chromatography on silica gel in MeOH/DCM using a gradient of 0–5% afforded **5** as a white solid (1.05 g, 51%).

**$^1$H NMR** (400 MHz, DMSO-$d_6$) δ 12.82 (s, 1H), 7.89 (d, $J$ = 7.5 Hz, 2H), 7.78 (d, $J$ = 8.5 Hz, 1H), 7.64 (dd, $J$ = 7.6, 3.3 Hz, 2H), 7.49 (d, $J$ = 1.9 Hz, 3H), 7.42 (td, $J$ = 7.4, 2.1 Hz, 2H), 7.38 – 7.25 (m, 2H), 5.76 (d, $J$ = 0.7 Hz, 1H), 4.28 – 4.13 (m, 4H), 3.17 (dd, $J$ = 13.9, 4.4 Hz, 1H), 2.94 (dd, $J$ = 13.9, 10.8 Hz, 1H).

##### 3.3.3 DBMB/TBMB Peptide Cyclisation

To a solution of purified linear peptide (30 mg) in DMF (1 mL) was added α,α'-dibromo-*m*-xylene (DBMB) (1.5 equiv.) or 1,3,5-tris(bromomethyl)benzene (TBMB) (1.5 equiv.), followed by DIPEA (4 equiv.). The reaction mixture was stirred at 37 °C for 30 min, and the progress of reaction monitored by LC-MS analysis. Upon completion, the crude reaction was concentrated *in vacuo* and purified by reversed-phase flash column chromatography on C18 silica gel. The purified cyclic peptides were then lyophilised and subsequently prepared in DMSO (10 mM) followed by storage at −80 °C.

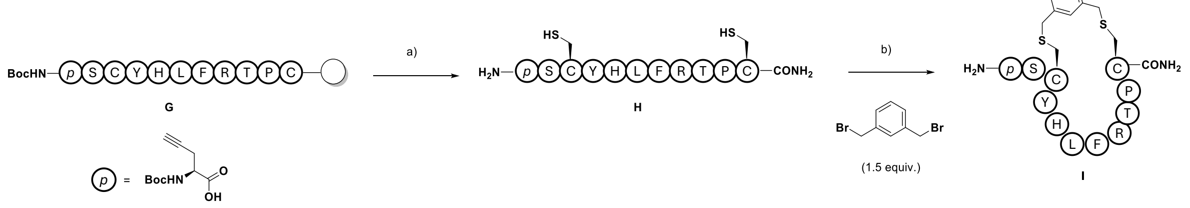

***Supplementary Scheme 3: DBMB cyclisation**. a) TFA (82.5%), $H_2O$ (5%), thioanisole (5%), phenol (5%) and DODt (2.5%), 2 h, RT. b) DBMB or TBMB (1.5 equiv.), DIPEA (4 equiv.), DMF, 30 min, 37 °C. p* = propargyl glycine

##### 3.3.4 Installation of Fluorosulfate Using AISF

For **Compounds 17–22**, incorporation of the Fmoc-Tyr-OH or unnatural amino acids (see **Supplementary Table 8**) was performed using conditions in **Section 3.2.1**, followed by deprotection with 20% (v/v) 4-methylpiperidine in DMF (3 x 5 mL washes, 3 min per treatment). The resin was then washed sequentially with DMF (2 x 5 mL), DCM (2 x 5 mL), and DMF (2 x 5 mL). A solution of 4-(acetylamino)phenylimidodisulfuryl difluoride (3 equiv.) in THF (4 mL) was then added, followed by 1,8-diazabicyclo[5.4.0]undec-7-ene (4 equiv.). The resin was agitated at RT for 30 min before being washed sequentially with DMF (x 2), DCM (x 2), and DMF (x 2). Following resin cleavage, the incorporation of the fluorosulfate group was confirmed by LC/MS analysis.

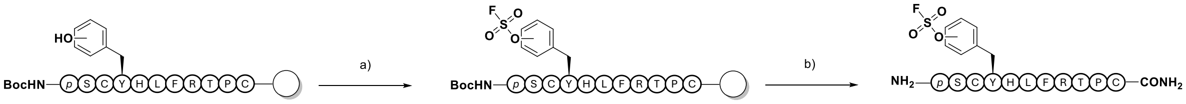

J K *17, 18, 19, 20, 21, 22*

***Supplementary Scheme 4: Solid-phase synthesis of compounds 17-22**. a) AISF (3 equiv.), DBU (4 equiv.), THF, 30 min, RT. b) TFA (82.5%), $H_2O$ (5%), thioanisole (5%), phenol (5%) and DODt (2.5%), RT, 2 h.*

##### 3.3.5 Installation of Sulfonyl Fluoride Using 5-(Fluorosulfonyl)-2-methoxybenzoic acid

For **Compounds 23–24**, selective deprotection of incorporated Dap(Mtt)-OH was performed on resin following chain completion. The Mtt protecting group was removed using 3% (v/v) TFA and 5% (v/v) triisopropylsilane (TIS) in DCM (3 x 5 mL, 15 min per treatment; the resin was washed with 2 × 5 mL DCM between treatments). After deprotection, the resin was washed sequentially with DMF (x 2), DCM (x 2), and DMF (x 2). The resin was then treated with 5-(fluorosulfonyl)-2-methoxybenzoic acid (1.5 equiv.), DIPEA (3 equiv.), and PyBOP (3 equiv.) in 2 mL DMF and agitated for 1 h at RT.

The resin was then washed with DMF (x 2), DCM (x 2), DMF (x 2), and DCM (x 2). Sulfonyl fluoride warhead incorporation was confirmed *via* LC-MS analysis.

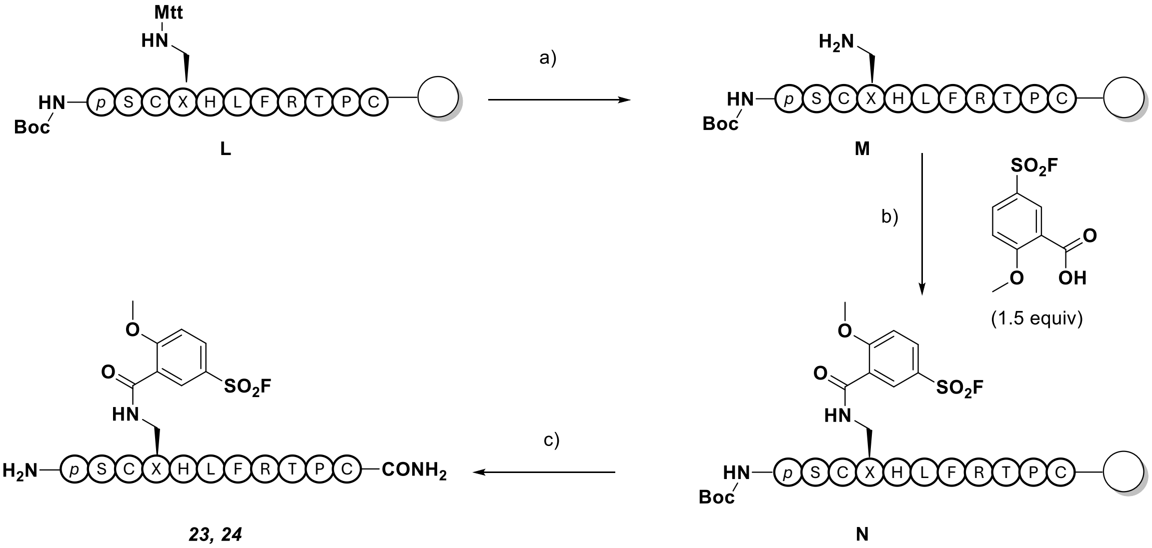

***Supplementary Scheme 5: Solid-phase synthesis of compounds 23-24****. a) 3% (v/v) TFA, 5% (v/v) TIS, DCM, 3 x 15 min, RT. b) 5-(Fluorosulfonyl)-2-methoxybenzoic acid (1.5 equiv.), PyBOP (3 equiv.), DIPEA (3 equiv.), 2 mL DMF, 60 min, RT. c) TFA (82.5%),* $H_2O$ *(5%), thioanisole (5%), phenol (5%) and DODt (2.5%) 2 h, RT.*

#### 3.3.6 Installation of Sulfonyl Fluoride Using 3/4/5-Benzylsulfonyl Fluoride

For **Compounds 25–27**, selective deprotection of incorporated Cys(STmp)-OH was performed on resin by treatment with 5% (w/v) DTT and *N*-methylmorpholine (NMM) (0.1 M) in DMF (5 mL, 3 x 5 min per treatment; the resin was washed with DMF (3 × 5 mL) between each treatment). After completion, the resin was washed with DMF (2), DCM (x 2), and DMF (x 2). The resin was then treated with a solution of substituted (bromomethyl)benzenesulfonyl fluoride (1.2 equiv.) and DIPEA (2.5 equiv.) in DMF (5 mL), followed by agitation at RT for 30 min. Incorporation of the sulfonyl fluoride warhead was confirmed *via* LC-MS analysis.

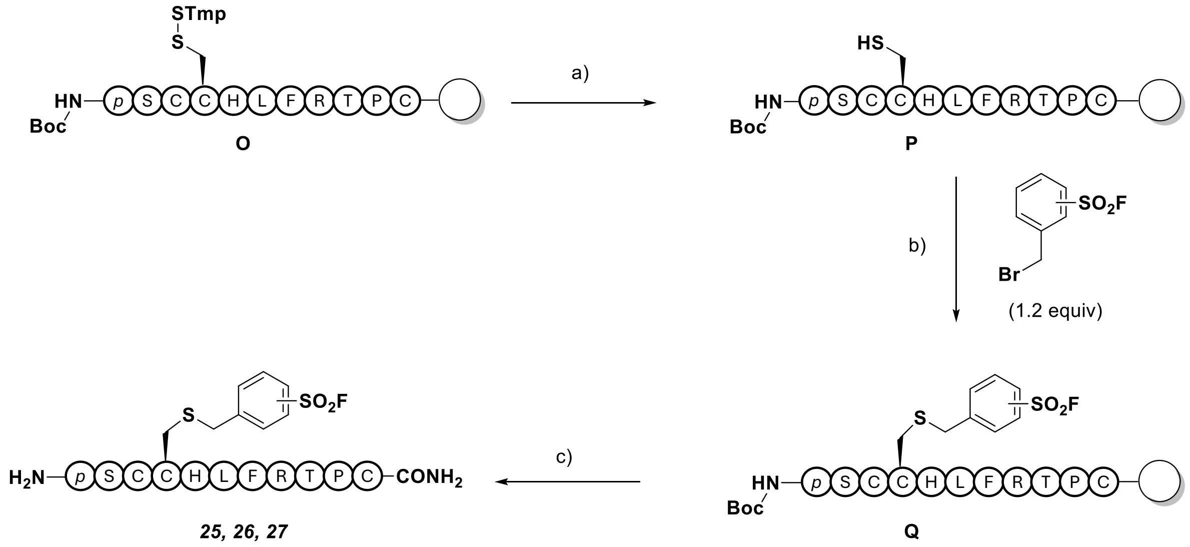

***Supplementary Scheme 6: Solid-phase synthesis of compounds 25-27.*** *a) 5% (v/v) DTT, 0.1 M N-methylmorpholine, DMF, 3 x 5 min, RT. b) (bromomethyl)benzenesulfonyl fluoride (1.2 equiv.), DIPEA (2.5 equiv.), DMF, 1 h, RT. c) TFA (82.5%), $H_2O$ (5%), thioanisole (5%), phenol (5%) and DODt (2.5%), 2 h, RT.*

##### 3.3.7 Synthesis of Diphenylphosphonate 28

2-Chlorotrityl chloride resin (150 mg, loading 1.22 mmol/g) was swollen in DCM for 10 min, followed by washing with DCM (3 x 5 mL). Fmoc-Phe-OH (5 equiv.) and DIPEA (10 equiv.) in DCM was then added and the reaction mixture was agitated for 2 h at RT. After 2 h, MeOH (500 μL) was added and the mixture agitated for a further 30 min at RT. The resin was then washed sequentially with DMF (x 2), DCM (x 2), and DMF (x 2). Removal of the Fmoc group was achieved by treating the resin with a 20% (v/v) solution of 4-methylpiperidine in DMF (3 x 5 mL, 3 min per treatment). The resin was then sequentially washed with DMF (x 2), DCM (x 2), and DMF (x 2).

Fmoc-Leu-OH, Fmoc-His(Trt)-OH and Fmoc-Aminohexanoic acid (Fmoc-Ahx-OH_ were coupled to the resin using standard SPPS procedures as described above. The peptide was capped with fluorescein (3 equiv.), DIC (3 equiv.) and Oxyma (3 equiv.) in 2 mL DMF overnight at RT, washed with DMF (x 2), DCM (x 2), DMF (x 2) and cleaved from the resin by treatment with 2 mL of a 1:3 (v/v) mixture of HFIP:DCM for 2 h at RT. The resultant filtrate was concentrated *in vacuo* followed by purification *via* reversed-phase flash column chromatography on C18 silica gel to afford **Intermediate W** (**Supplementary Scheme 8**).

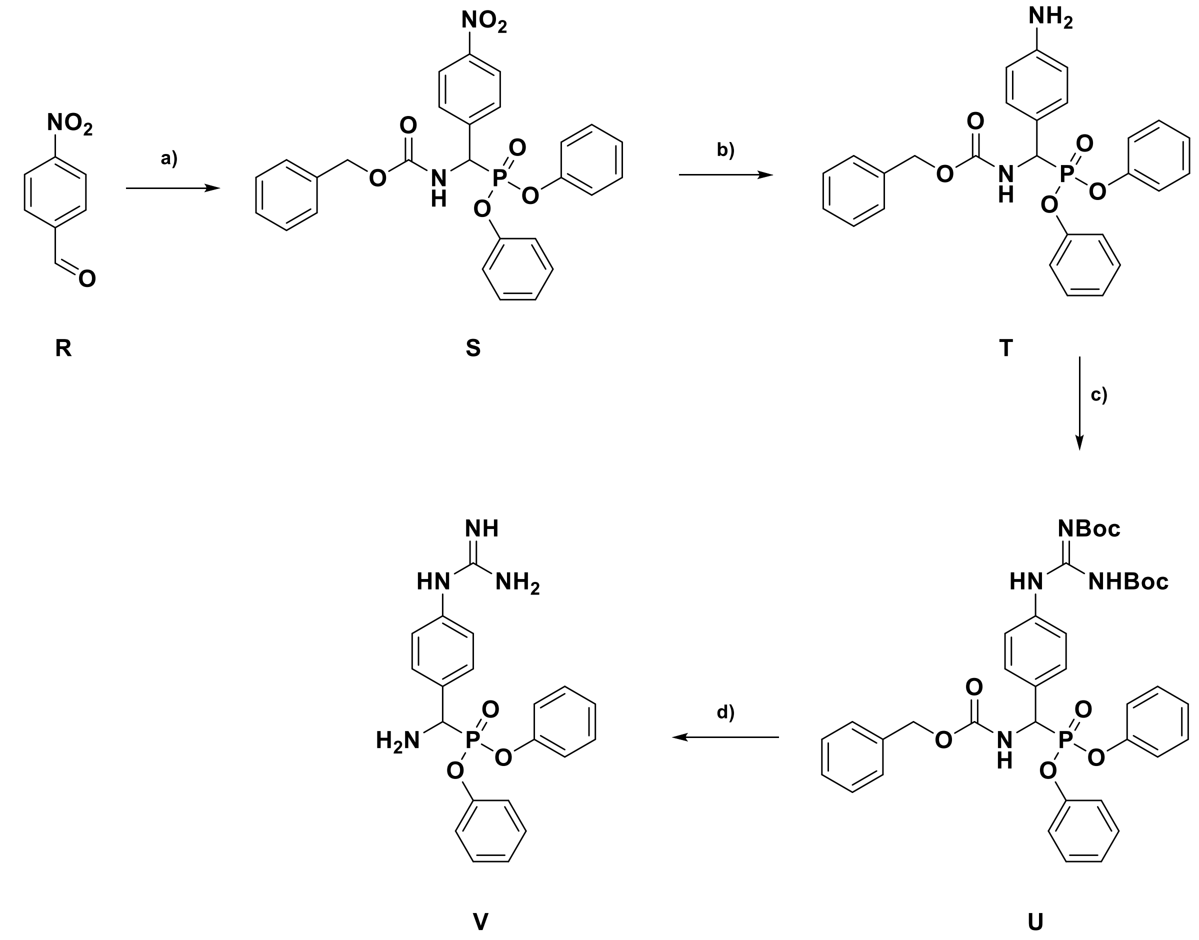

***Supplementary Scheme 7: Synthesis of Diphenylphosphonate warhead.*** *a)* 4-nitrobenzaldehyde, $P(OPh)_3$, benzyl carbamate, AcOH, 90 °C, 2 h. b) Fe, AcOH, 70 °C, 2 h. *c.* *N*,*N'*-bis-Boc-1-guanylpyrazole, $NEt_3$, DCM, RT, 16 h. d) 33% HBr in AcOH, RT, 2 h.

**Benzyl ((diphenoxyphosphoryl)(4-nitrophenyl)methyl)carbamate (Intermediate S)**

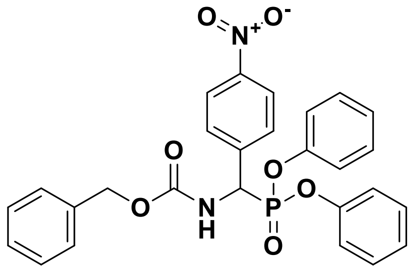

**S**

A solution of 4-nitrobenzaldehyde (**Intermediate R, Supplementary Scheme 7**) (0.91 g, 6.00 mmol, 1.5 equiv.), benzyl carbamate (1.24 g, 4.00 mmol, 1.0 equiv.), and triphenyl phosphite (1.05 mL, 4.00 mmol, 1.0 equiv.) in glacial acetic acid (1.2 mL) was stirred at 90 °C for 2 h. Upon completion (as monitored by TLC), the mixture was concentrated *in vacuo* and recrystalised from MeOH (washed with ice-cold 5 mL of MeOH and $Et_2O$) to afford **Intermediate S** as a white solid (1.16 g, 56%).

**$^1$H NMR** (500 MHz, DMSO-$d_6$) $\delta$ 9.09 (d, $J$ = 10.1 Hz, 1H), 8.27 (d, $J$ = 8.5 Hz, 2H), 7.94 (dd, $J$ = 8.9, 2.2 Hz, 2H), 7.40 – 7.28 (m, 9H), 7.20 (td, $J$ = 7.4, 3.9 Hz, 2H), 7.04 (dd, $J$ = 16.7, 8.0 Hz, 4H), 5.91 – 5.81 (m, 1H), 5.14 (d, $J$ = 12.5 Hz, 1H), 5.07 (d, $J$ = 12.4 Hz, 1H).

**Benzyl (*R*)-((4-aminophenyl)(diphenoxyphosphoryl)methyl)carbamate (Intermediate T)**

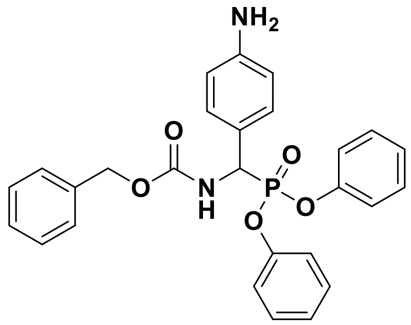

**T**

**Intermediate S** (3.00 g, 6.00 mmol, 1.0 equiv.) and iron powder (3.00 g, 53.7 mmol, 9.0 equiv.) was dissolved in glacial acetic acid (24 mL) and stirred at 70 °C for 2 h. The reaction mixture was concentrated *in vacuo* and the crude residue was suspended in

50 mL EtOAc. The suspension was centrifuged for 5 min at 3000 rpm and the supernatant was collected and concentrated *in vacuo* to afford **Intermediate T** as a brown solid (2.61 g, 5.34 mmol, 89%) which was carried forward without further purification.

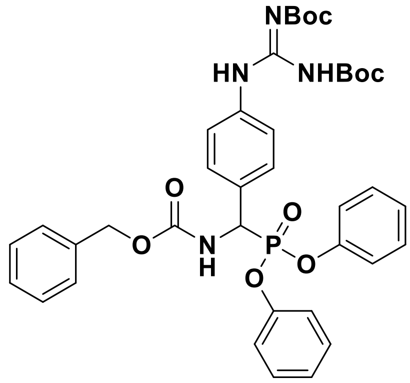

**U**

***ii*: Guanylation.** To a solution of **Intermediate T** (2.61 g, 5.34 mmol, 1.0 equiv.) in DCM (30 mL) was added *N,N'*-bis-Boc-1-guanylpyrazole (1.66 g, 5.34 mmol, 1.0 equiv.) and $Et_3N$ (1.49 mL, 10.70 mmol, 2.0 equiv.) which was allowed to stir at RT overnight. Following completion, the reaction mixture was concentrated *in vacuo*, dissolved in 50 mL EtOAc, and washed sequentially with 20 mL of 1 M HCl, sat. aq. $NaHCO_3$, and brine. The organic layer was dried ($MgSO_4$), filtered, and concentrated *in vacuo*. Purification *via* flash column chromatography on silica gel in 40-60° petroleum ether/EtOAc using a gradient of 10–100% afforded **Intermediate U** as a white solid (1.80 g, 45% over two steps).

**$^1$H NMR** (500 MHz, DMSO-$d_6$) $\delta$ 11.37 (s, 1H), 10.00 (s, 1H), 8.86 (d, $J$ = 10.1 Hz, 1H), 7.57 (q, $J$ = 8.7 Hz, 4H), 7.33 (p, $J$ = 8.1 Hz, 8H), 7.18 (t, $J$ = 7.4 Hz, 2H), 7.03 (d, $J$ = 8.0 Hz, 2H), 6.96 (d, $J$ = 8.0 Hz, 2H), 5.61 – 5.51 (m, 1H), 5.12 (d, $J$ = 12.7 Hz, 1H), 5.04 (d, $J$ = 12.3 Hz, 1H), 1.49 (s, 9H), 1.38 (s, 9H).

**Diphenyl (amino(4-guanidinophenyl)methyl)phosphonate (Intermediate V)**

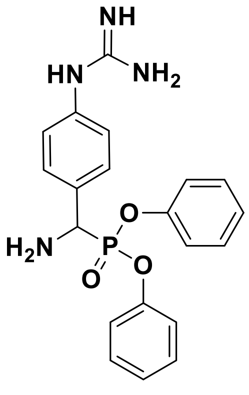

**V**

**Intermediate U** (0.75 g, 1.02 mmol) was treated with a 33% (w/w) solution of HBr in acetic acid (5 mL) and allowed to stir for 2 h at RT. After 2 h, the reaction mixture was concentrated *in vacuo* followed by purification by recrystallization from MeOH/$Et_2O$ to afford **Intermediate V** as a white solid (0.34 g, 83%).

**$^1$H NMR** (500 MHz, DMSO-$d_6$) δ 9.88 (s, 1H), 9.41 (s, 3H), 7.73 (dd, $J$ = 8.7, 2.1 Hz, 2H), 7.55 (s, 4H), 7.38 (dq, $J$ = 15.3, 7.8 Hz, 5H), 7.24 (dt, $J$ = 14.9, 7.4 Hz, 2H), 7.14 (d, $J$ = 7.9 Hz, 2H), 7.02 (d, $J$ = 8.0 Hz, 2H), 5.75 – 5.65 (m, 1H).

#### Diphenyl Phosphonate ABP (28)

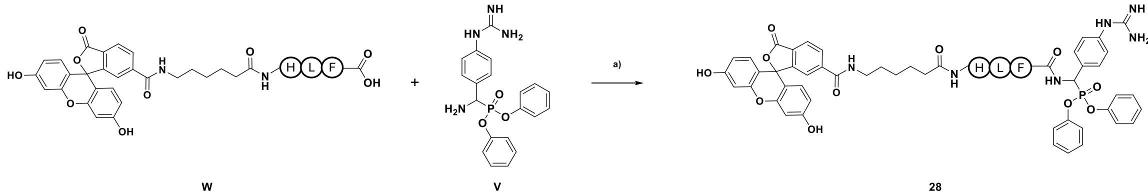

***Supplementary Scheme 8: Synthesis of 28.*** *a) HATU, 2,4,6-trimethylpyridine, 2 h, RT.*

To a solution of **Intermediate W,** (**Supplementary Scheme 8**) (25.0 mg, 29.7 µmol, 1 equiv.) in DMF (1 mL) was added HATU (11.3 mg, 29.7 µmol, 1 equiv.) and 2,4,6-trimethylpyridine (19.6 µL, 148.5 µmol, 5 equiv.). The reaction mixture was stirred for 3 min, then **Intermediate V** (**Supplementary Scheme 8**) (14.2 mg, 35.6 µmol, 1.2 equiv.) was added and the reaction mixture was stirred for 2 h at RT. After completion (as monitored by LC-MS analysis), the reaction mixture was concentrated *in vacuo* and the resulting residue was then treated with a 95:2.5:2.5 solution of TFA/TIS/$H_2O$ (1 mL) and stirred for 2h at RT. The reaction mixture was concentrated *in vacuo* and purification *via* reversed-phase flash column chromatography on $C_{18}$ silica gel afforded **28** as a (2.00 mg, 6%).

### 4 Supplementary Figures

#### 4.1 Volcano Plot of NGS Data

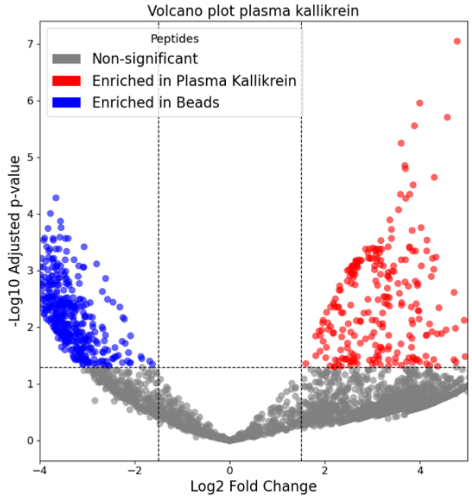

***Supplementary Figure 1: Volcano plot showing enrichment of peptides from phage display against plasma kallikrein relative to a streptavidin bead control.*** *The x-axis represents $\log_2$ fold change in peptide frequency, and the y-axis shows statistical significance (–$\log_{10}$ adjusted p-value). Dashed lines indicate the significance thresholds (adj. $p < 0.05$, $\log_2 FC > 1.5$). Peptides enriched in plasma kallikrein and control samples are highlighted in red and blue, respectively; non-significant peptides are shown in grey.*

#### 4.2 Labelling of Plasma Kallikrein Mutants with FP-TAMRA

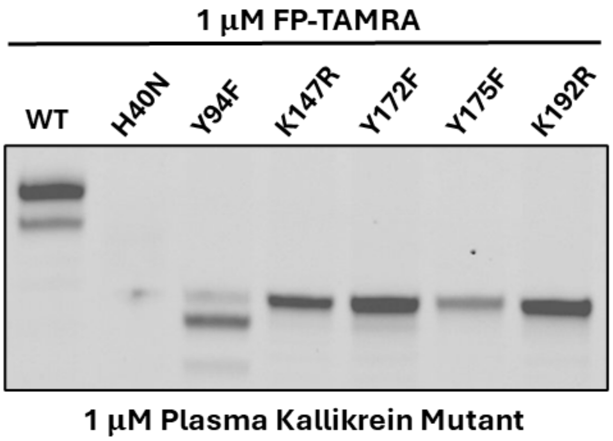

***Supplementary Figure 2: Labelling of plasma kallikrein mutants by FP-TAMRA.*** *Mutants (1 µM) were incubated with the broad-spectrum activity-based probe FP-TAMRA (1 µM) in PBS (pH 7.4) for 60 min at 37 °C. Following SDS-PAGE resolution, covalent labelling was visualised via in-gel TAMRA fluorescence.*

#### 4.3 Inhibition Constant Plots for 10 and 12

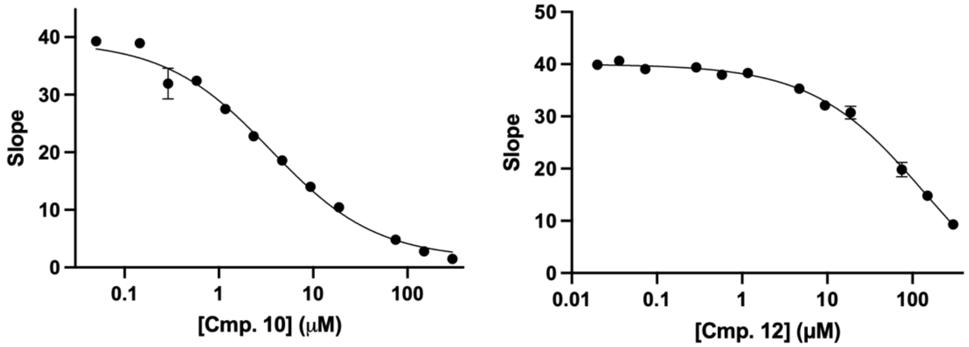

| Compound | $K_i$ (µM) |
|---|---|
| 10 | 3.53 ± 0.7 |
| 12 | >100 |

***Supplementary Figure 3: Determination of inhibition constants (Ki) for compounds 10 and 12.*** *Dose-response curves for the residual enzymatic activity of human plasma kallikrein in the presence of varying concentrations of compound 10 (left) and 12 (right). Enzymatic activity is represented by the slope of the kinetic progress curves. Data points indicate the mean ± SD of three independent replicates. Solid lines represent the non-linear regression fit to a standard dose-response model using GraphPad Prism. The accompanying table summarises the calculated $K_i$ values for each compound.*

#### 4.4 Tryptic-Digest LC-MS/MS

| Peptide Sequence | Modified Sequence | Charges | Assigned Modifications | Protein | Kallikrein Spectral Count | Kallikrein_CmP23 Spectral Count | Kallikrein Intensity | Kallikrein_CmP23 Intensity |
|---|---|---|---|---|---|---|---|---|
| GDTSTIYTNCWVTGWGFSK | GDTSTIYTNC[57.0215]WVTGWGFSK | 2,3 | 10C(57.0215) | Kallikrein | 8 | 4 | 8.80E+08 | 7.10E+08 |
| GDTSTIYTNCWVTGWGFSKEK | GDTSTIYTNC[57.0215]WVTGWGFSKEK | 3 | 10C(57.0215) | Kallikrein | 3 | 3 | 1.00E+07 | 2288659.2 |
| GDTSTIYTNCWVTGWGFSKEKGEIQNILQK | GDTSTIYTNC[57.0215]WVTGWGFSKEKGEIQNILQK | 4 | 10C(57.0215) | Kallikrein | 1 | 1 | 1.69E+07 | 6549460 |
| LVGITSWGEGCAR | LVGITSWGEGC[57.0215]AR | 2,3 | 11C(57.0215) | Kallikrein | 21 | 17 | 1.09E+10 | 1.21E+10 |
| LVGITSWGEGCARR | LVGITSWGEGC[57.0215]ARR | 2,3 | 11C(57.0215) | Kallikrein | 5 | 1 | 9.35E+09 | 1.25E+10 |
| LVGITSWGEGCARREQPGVYTK | LVGITSWGEGC[57.0215]ARREQPGVYTK | 2,3 | 11C(57.0215) | Kallikrein | 3 | 0 | 6.97E+08 | 0 |
| VNIPLVTNEECQK | VNIPLVTNEEC[57.0215]QK | 2,3 | 11C(57.0215) | Kallikrein | 18 | 7 | 1.07E+10 | 1.04E+10 |
| VNIPLVTNEECQKR | VNIPLVTNEEC[57.0215]QKR | 2,3 | 11C(57.0215) | Kallikrein | 30 | 20 | 2.63E+10 | 2.80E+10 |
| **DACKGDSGGPLVCKHNGMWR** | DAC[57.0215]KGDSGGPLVC[57.0215]KHNGM[15.9949]WR | 4 | 13C(57.0215),18M(15.9949),3C(57.0215) | Kallikrein | 0 | 1 | 0 | 1074719.8 |
| **DACKGDSGGPLVCK** | DAC[57.0215]KGDSGGPLVC[57.0215]K | 2,3 | 13C(57.0215),3C(57.0215) | Kallikrein | 8 | 4 | **1.64E+09** | **4.64E+08** |
| **DACKGDSGGPLVCKHNGMWR** | DAC[57.0215]KGDSGGPLVC[57.0215]KHNGMWR | 3,4,5 | 13C(57.0215),3C(57.0215) | Kallikrein | 8 | 6 | 2.69E+07 | 1.66E+07 |
| **DACKGDSGGPLVCK** | DAC[57.0215]K[1556.5999]GDSGGPLVC[57.0215]K | 3,4,5 | 13C(57.0215),3C(57.0215),4K(1556.5999) | Kallikrein | **0** | **4** | 0 | 0 |
| MVCAGYKEGGKDACK | M[15.9949]VC[57.0215]AGYKEGGKDAC[57.0215]K | 3,4 | 14C(57.0215),1M(15.9949),3C(57.0215) | Kallikrein | 3 | 2 | 3083475.5 | 395483.16 |
| MVCAGYKEGGKDACK | MVC[57.0215]AGYKEGGKDAC[57.0215]K | 2,3,4 | 14C(57.0215),3C(57.0215) | Kallikrein | 4 | 2 | 1.91E+07 | 1.10E+07 |
| **EGGKDACKGDSGGPLVCK** | EGGK[1556.5999]DAC[57.0215]KGDSGGPLVC[57.0215]K | 5,6 | 17C(57.0215),4K(1556.5999),7C(57.0215) | Kallikrein | **0** | **7** | 0 | 0 |
| **EGGKDACKGDSGGPLVCK** | EGGKDAC[57.0215]KGDSGGPLVC[57.0215]K | 2,3,4 | 17C(57.0215),7C(57.0215) | Kallikrein | 33 | 17 | **4.41E+09** | **1.22E+08** |
| **EGGKDACKGDSGGPLVCK** | EGGKDAC[57.0215]K[1556.5999]GDSGGPLVC[57.0215]K | 3,7 | 17C(57.0215),7C(57.0215),8K(1556.5999) | Kallikrein | 0 | 3 | 0 | 0 |
| HLCGGSLIGHQWVLTAAHCFDGLPLQDVWR | HLC[57.0215]GGSLIGHQWVLTAAHC[57.0215]FDGLPLQDVWR | 3,4,5 | 19C(57.0215),3C(57.0215) | Kallikrein | 36 | 16 | 7.56E+07 | 9.14E+07 |
| MVCAGYK | M[15.9949]VC[57.0215]AGYK | 2 | 1M(15.9949),3C(57.0215) | Kallikrein | 8 | 6 | 7.97E+07 | 9493074 |
| MVCAGYKEGGK | M[15.9949]VC[57.0215]AGYKEGGK | 2,3 | 1M(15.9949),3C(57.0215) | Kallikrein | 6 | 3 | 1.32E+07 | 999056.2 |
| GEIQNILQKVNIPLVTNEECQK | GEIQNILQKVNIPLVTNEEC[57.0215]QK | 3 | 20C(57.0215) | Kallikrein | 1 | 2 | 5187395 | 2894033 |
| GEIQNILQKVNIPLVTNEECQKR | GEIQNILQKVNIPLVTNEEC[57.0215]QKR | 3,4 | 20C(57.0215) | Kallikrein | 2 | 1 | 1.41E+08 | 3.77E+07 |
| EKGEIQNILQKVNIPLVTNEECQK | EKGEIQNILQKVNIPLVTNEEC[57.0215]QK | 3 | 22C(57.0215) | Kallikrein | 1 | 0 | 2.53E+07 | 1.15E+07 |
| LQAPLEYTEFQKPISLPSKGDTSTIYTNCWVTGWGFSK | LQAPLEYTEFQKPISLPSKGDTSTIYTNC[57.0215]WVTGWGFSK | 3,4,5 | 29C(57.0215) | Kallikrein | 23 | 9 | 2.55E+09 | 1.94E+09 |
| LQAPLEYTEFQKPISLPSKGDTSTIYTNCWVTGWGFSKEK | LQAPLEYTEFQKPISLPSKGDTSTIYTNC[57.0215]WVTGWGFSKEK | 5 | 29C(57.0215) | Kallikrein | 3 | 3 | 1.15E+07 | 2.41E+07 |
| MVCAGYK | MVC[57.0215]AGYK | 2 | 3C(57.0215) | Kallikrein | 11 | 3 | 4.45E+08 | 1.58E+08 |
| MVCAGYKEGGK | MVC[57.0215]AGYKEGGK | 2,3 | 3C(57.0215) | Kallikrein | 5 | 4 | 1.92E+08 | 3.93E+07 |
| VAEYMDWILEK | VAEYM[15.9949]DWILEK | 2 | 5M(15.9949) | Kallikrein | 16 | 11 | 5.10E+08 | 3.73E+08 |
| VAEYMDWILEKTQSSDGK | VAEYM[15.9949]DWILEKTQSSDGK | 2,3 | 5M(15.9949) | Kallikrein | 6 | 1 | 7.12E+07 | 1.97E+07 |
| GDSGGPLVCK | GDSGGPLVC[57.0215]K | 2 | 9C(57.0215) | Kallikrein | 2 | 1 | 1.50E+08 | 4.64E+07 |
| GDSGGPLVCKHNGMWR | GDSGGPLVC[57.0215]KHNGMWR | 2,3 | 9C(57.0215) | Kallikrein | 3 | 1 | 3762560.5 | 1.67E+07 |
| DTPFSQIK | DTPFSQIK | 2 | | Kallikrein | 3 | 0 | 1.98E+09 | 1.36E+09 |
| DTPFSQIKEIIIHQNYK | DTPFSQIKEIIIHQNYK | 2,3,4 | | Kallikrein | 4 | 3 | 1.40E+09 | 8.55E+08 |
| EIIIHQNYK | EIIIHQNYK | 2 | | Kallikrein | 9 | 3 | 8.38E+08 | 7.82E+08 |
| EIIIHQNYKVSEGNHDIALIK | EIIIHQNYKVSEGNHDIALIK | 3,4,5 | | Kallikrein | 4 | 2 | 4.25E+08 | 4.49E+08 |
| EKGEIQNILQK | EKGEIQNILQK | 2,3 | | Kallikrein | 27 | 10 | 2.33E+10 | 2.18E+10 |
| EQPGVYTK | EQPGVYTK | 2 | | Kallikrein | 2 | 0 | 2.28E+08 | 8.02E+07 |
| EQPGVYTKVAEYMDWILEK | EQPGVYTKVAEYMDWILEK | 2,3 | | Kallikrein | 1 | 3 | 2.07E+07 | 1.37E+07 |
| EQPGVYTKVAEYMDWILEKTQSSDGK | EQPGVYTKVAEYMDWILEKTQSSDGK | 3,4 | | Kallikrein | 2 | 2 | 2.06E+07 | 1.90E+07 |
| GEIQNILQK | GEIQNILQK | 2 | | Kallikrein | 2 | 0 | 5.74E+09 | 3.71E+09 |
| IVGGTESSWGEWPWQVSLQVK | IVGGTESSWGEWPWQVSLQVK | 2,3 | | Kallikrein | 13 | 3 | 9.96E+08 | 9.15E+08 |
| IVGGTESSWGEWPWQVSLQVKLTAQR | IVGGTESSWGEWPWQVSLQVKLTAQR | 3 | | Kallikrein | 1 | 2 | 0 | 4355353.5 |
| IYSGILELSDITK | IYSGILELSDITK | 2,3 | | Kallikrein | 12 | 3 | 3.54E+09 | 1.78E+09 |
| IYSGILELSDITKDTPFSQIK | IYSGILELSDITKDTPFSQIK | 2,3,4 | | Kallikrein | 14 | 4 | 1.52E+09 | 5.66E+08 |
| IYSGILELSDITKDTPFSQIKEIIIHQNYK | IYSGILELSDITKDTPFSQIKEIIIHQNYK | 3,4,5 | | Kallikrein | 6 | 3 | 4.74E+08 | 7.77E+08 |
| LQAPLEYTEFQKPISLPSK | LQAPLEYTEFQKPISLPSK | 2,3,4 | | Kallikrein | 7 | 7 | 9.11E+09 | 6.43E+09 |
| REQPGVYTK | REQPGVYTK | 2 | | Kallikrein | 11 | 6 | 2.48E+08 | 5.61E+07 |
| REQPGVYTKVAEYMDWILEK | REQPGVYTKVAEYMDWILEK | 3,4 | | Kallikrein | 2 | 2 | 1.01E+07 | 1.08E+07 |
| VAEYMDWILEK | VAEYMDWILEK | 2,3 | | Kallikrein | 45 | 45 | 9.40E+09 | 1.10E+10 |
| VAEYMDWILEKTQSSDGK | VAEYMDWILEKTQSSDGK | 2,3 | | Kallikrein | 8 | 5 | 1.32E+09 | 9.06E+08 |
| VAEYMDWILEKTQSSDGKAQMQSPAHHHHHH | VAEYMDWILEKTQSSDGKAQMQSPAHHHHHH | 5 | | Kallikrein | 0 | 2 | 0 | 1.30E+07 |
| VSEGNHDIALIK | VSEGNHDIALIK | 2 | | Kallikrein | 4 | 4 | 9.15E+09 | 8.17E+09 |
| VSEGNHDIALIKLQAPLEYTEFQKPISLPSK | VSEGNHDIALIKLQAPLEYTEFQKPISLPSK | 4,5 | | Kallikrein | 3 | 3 | 1.58E+08 | 3.33E+07 |

***Supplementary Figure 4*. *LC–MS/MS identification of plasma kallikrein peptides modified by 23*.** *Peptides containing the expected single-oxidized **23** adduct (+1556.5999 Da) are highlighted in yellow. These modified peptides were detected exclusively in the cmABP-treated sample ('Kallikrein-Cmp23), and all contain Lys192. The exact residue modified by **23** cannot be unambiguously assigned due to complex MS2-spectra.*

| Peptide species | Monoisotopic mass (Da) | z=2 | z=3 | z=4 | z=5 | z=6 | z=7 | Observation |
|---|---|---|---|---|---|---|---|---|
| Apo peptide (2×CAM) | 1833.8349 | 917.9247 | 612.2856 | 459.4660 | 367.7743 | 306.6464 | 262.9837 | Detected in apo and **23**-treated samples (RT = 18.7 min) |
| Oxidized 23-modified peptide (2×CAM) | 3390.4348 | 1696.2247 | 1131.1522 | 848.6160 | 679.0942 | 566.0797 | 485.3551 | Detected only in **23**-treated sample (RT = 30.71 min) |

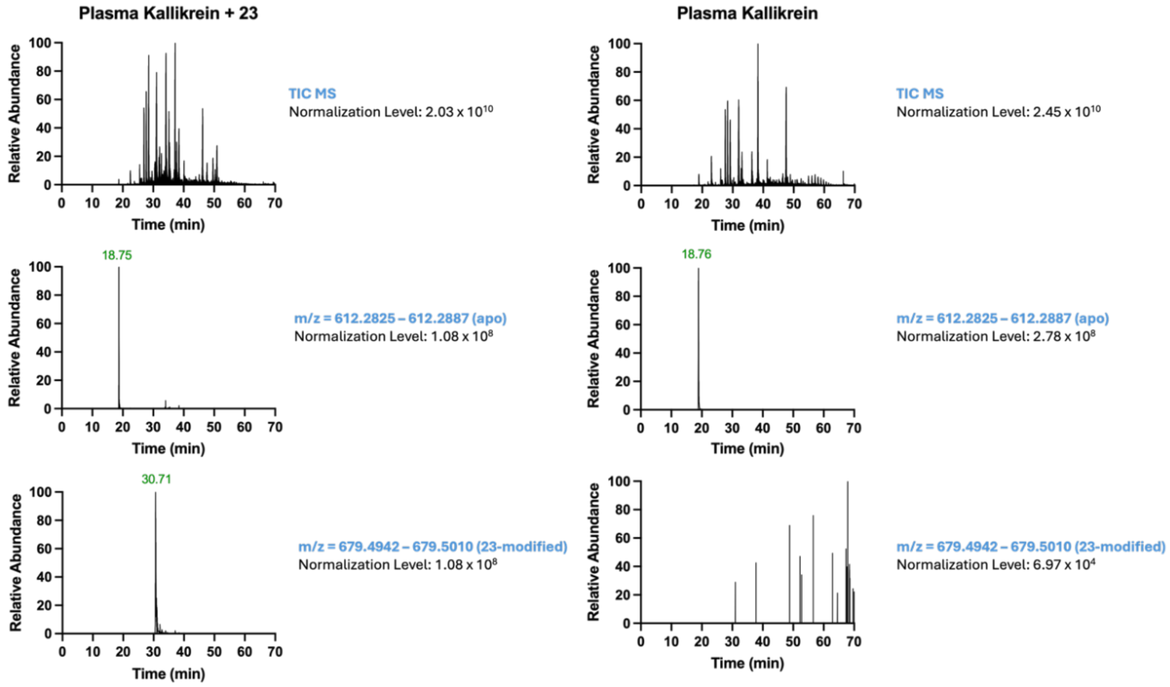

***Supplementary Figure 5: HPLC–MS analysis of apo and 23-modified plasma kallikrein peptide EGGKDACKGDSGGPLVCK using FreeStyle.*** *(**a**) Summary of the expected monoisotopic masses and charge states for unmodified and **23**-modified peptide ions. (**b**) Total ion chromatograms (TICs) and extracted mass spectra for **23**-treated and untreated samples, showing detection of the 3+ ion corresponding to the unmodified peptide in both samples, whereas the 5+ ion corresponding to the oxidized **23**-modified peptide is detected exclusively in the **23**-treated sample.*

#### 4.5 In-gel Fluorescence Profiling Against Serine Protease Panel

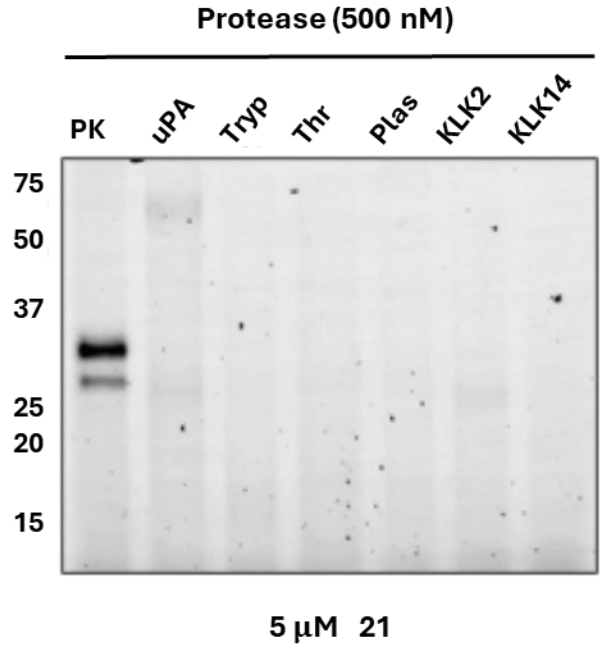

***Supplementary Figure 6: In-gel Fluorescence profiling of 21 specificity across a serine protease panel.*** *Recombinant proteases (500 nM) were incubated with* ***21*** *(5 µM) in PBS (pH 7.4) for 16 h at 37 °C. Following SDS-PAGE resolution, covalent labelling was visualised via in-gel fluorescence following CuAAC conjugation to TAMRA-azide. Protease abbreviations: PK, plasma kallikrein; uPA, urokinase-type plasminogen activator; Tryp, trypsin; Thr, thrombin; Plas, plasmin; KLK2, kallikrein-2; KLK14, kallikrein-14. Molecular weight markers (in kDa) are indicated on the left.*

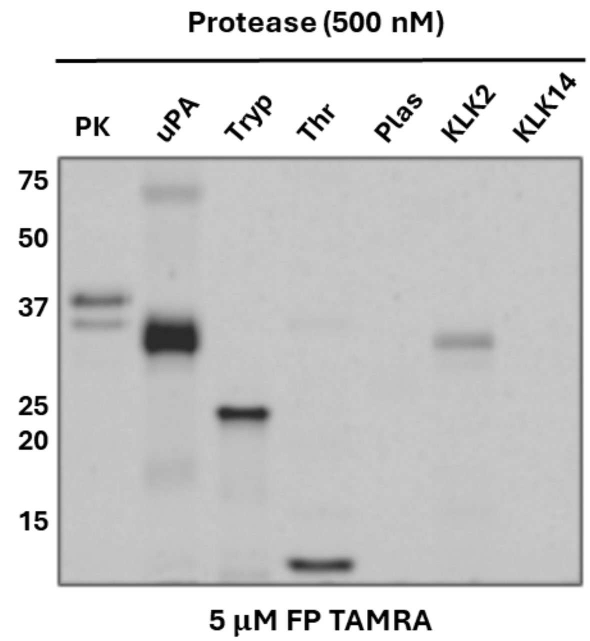

***Supplementary Figure 7: In-gel fluorescence profiling of FP-TAMRA reactivity across a serine protease panel.*** *Recombinant proteases (500 nM) were incubated with the broad-spectrum activity-based probe FP-TAMRA (5 µM) in PBS (pH 7.4) for 60 min at 37 °C. Following SDS-PAGE resolution, covalent labelling was visualised via in-gel TAMRA fluorescence. Protease abbreviations: PK, plasma kallikrein; uPA, urokinase-type plasminogen activator; Tryp, trypsin; Thr, thrombin; Plas, plasmin; KLK2, kallikrein-2; KLK14, kallikrein-14. Molecular weight markers (in kDa) are indicated on the left.*

#### 4.6 PC3 Cell Media Labelling with FP TAMRA

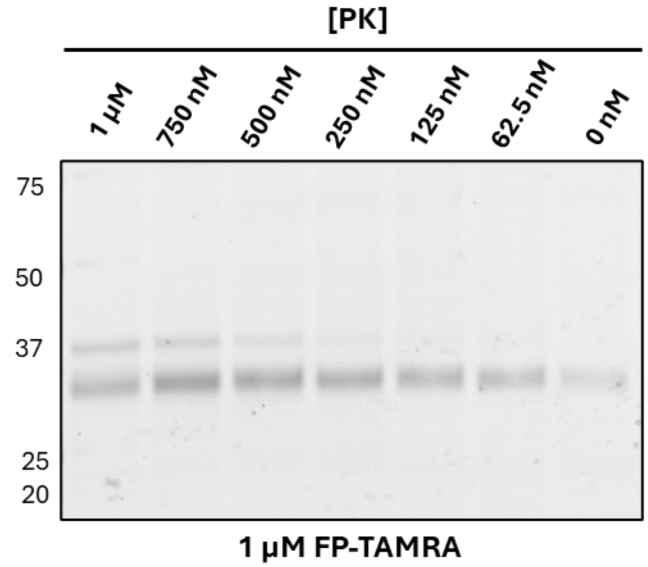

***Supplementary Figure 8*: *Labelling of PC3 cell media with FP-TAMRA.*** *Conditioned Media (10 µg) was incubated with FP-TAMRA (5 µM) for 60 min at 37 °C. Following SDS-PAGE resolution, covalent labelling was visualised via in-gel TAMRA fluorescence.*

### 5 Probe Analytical Data

**Compound 1**

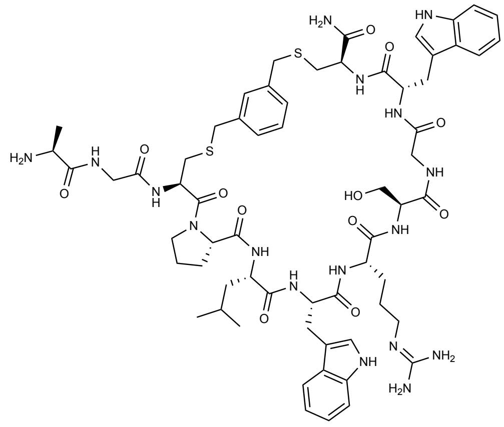

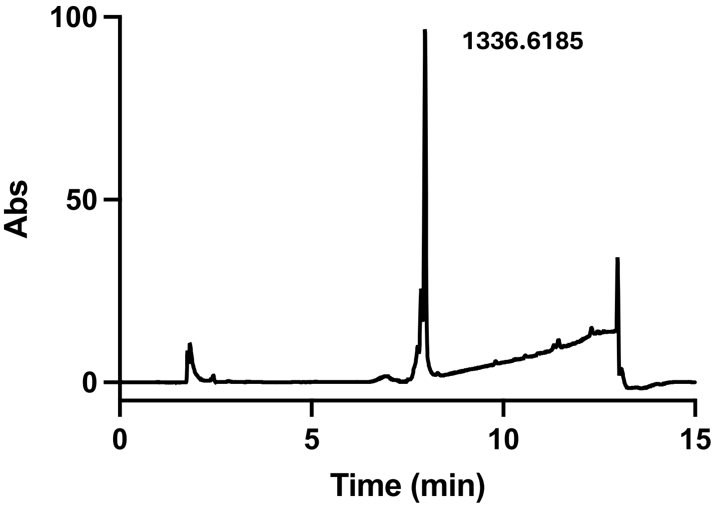

**Chemical formula: $C_{63}H_{85}N_{17}O_{12}S_2^+$**

**Calculated $[M + H]^+$ = 1336.60**

**Observed $[M + H]^+$ = 1336.6185**

**Compound 2**

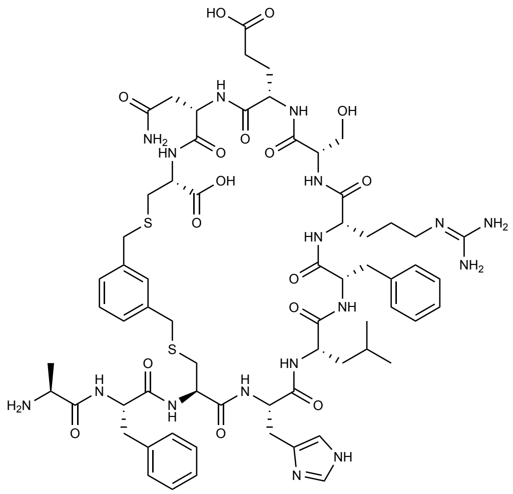

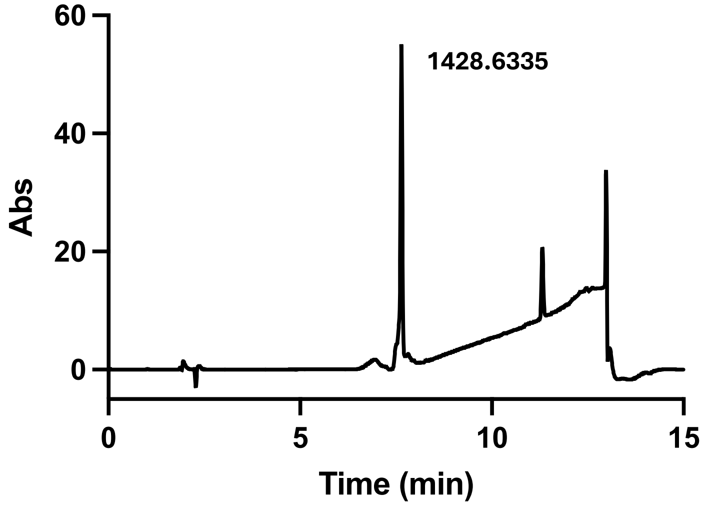

**Chemical formula: $C_{65}H_{89}N_{17}O_{16}S_2^+$**

**Calculated $[M + H]^+$ = 1428.61**

**Observed $[M + H]^+$ = 1428.6335**

**Compound 3**

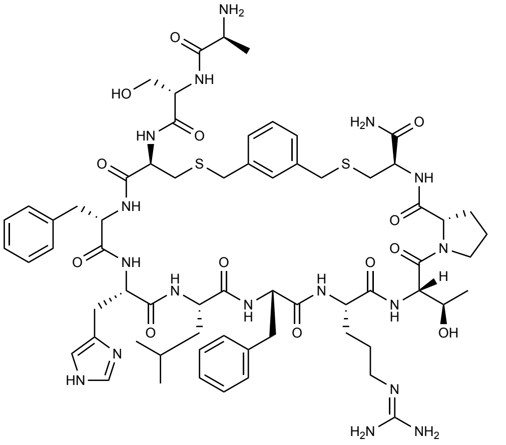

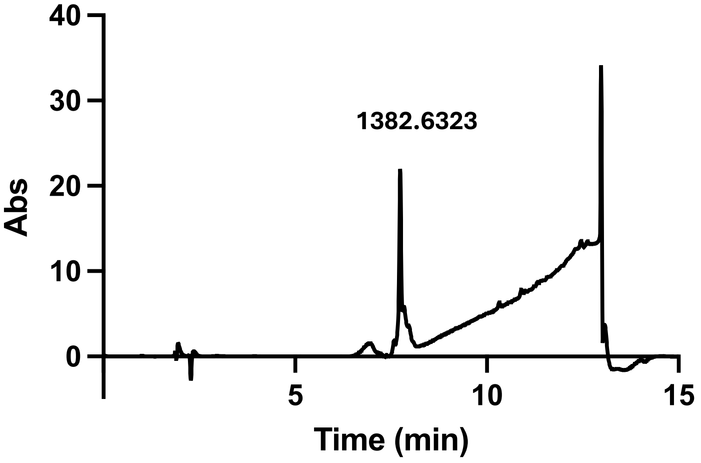

**Chemical formula: $C_{65}H_{91}N_{17}O_{13}S_2^+$**

**Calculated $[M + H]^+$ = 1382.64**

**Observed $[M + H]^+$ = 1382.6323**

**Compound 4**

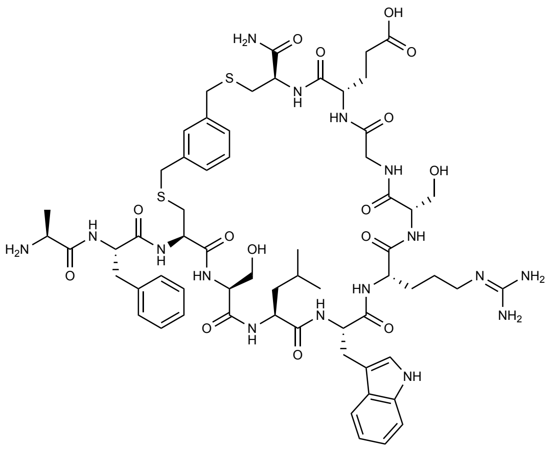

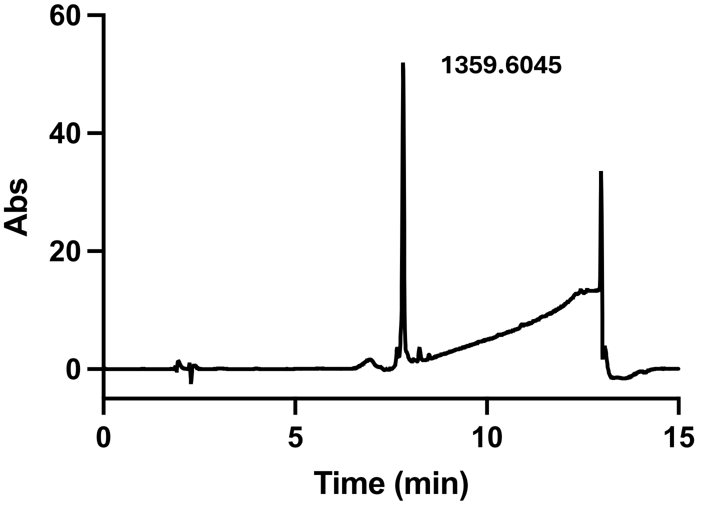

**Chemical formula: $C_{62}H_{85}N_{17}O_{12}S_2^+$**

**Calculated $[M + H]^+$ = 1359.59**

**Observed $[M + H]^+$ = 1359.6045**

**Compound 5**

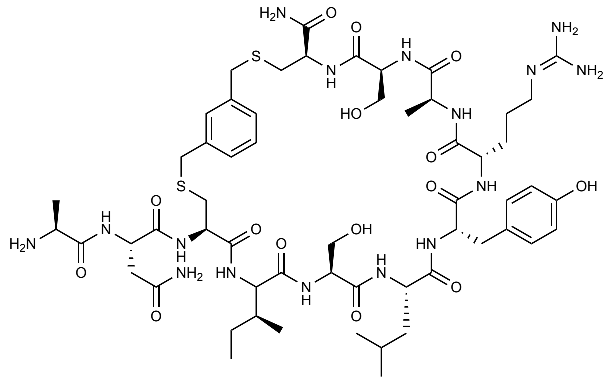

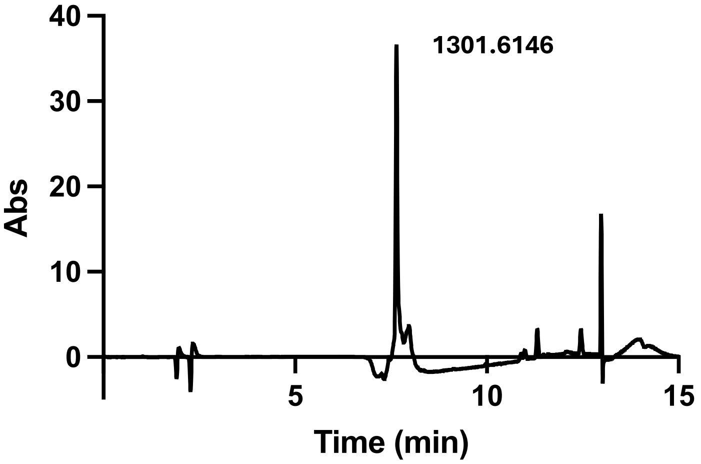

**Chemical formula: $C_{57}H_{88}N_{16}O_{15}S_2^+$**

**Calculated $[M + H]^+$ = 1301.55**

**Observed $[M + H]^+$ = 1301.6146**

**Compound 6**

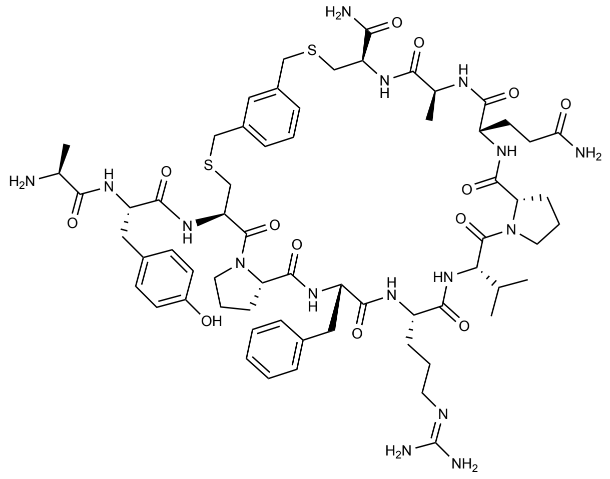

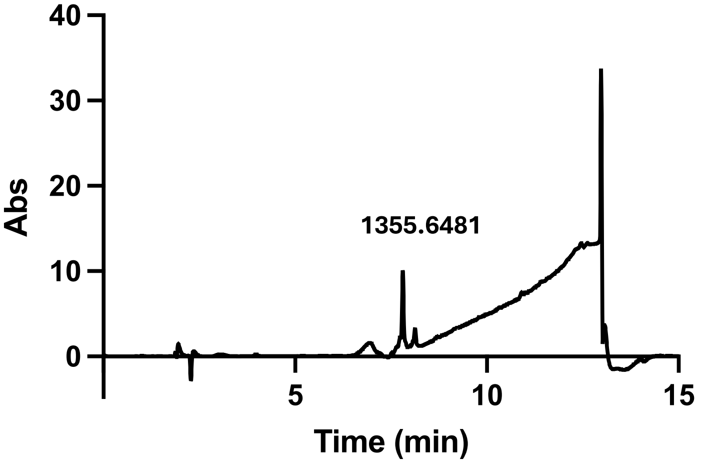

**Chemical formula: $C_{64}H_{90}N_{16}O_{13}S_2^+$**

**Calculated $[M + H]^+$ = 1355.64**

**Observed $[M + H]^+$ = 1355.6481**

**Compound 7**

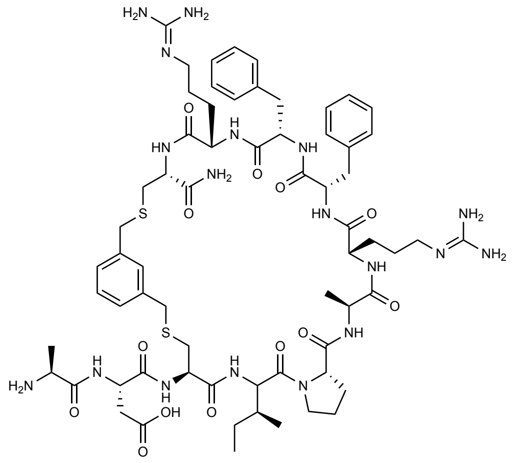

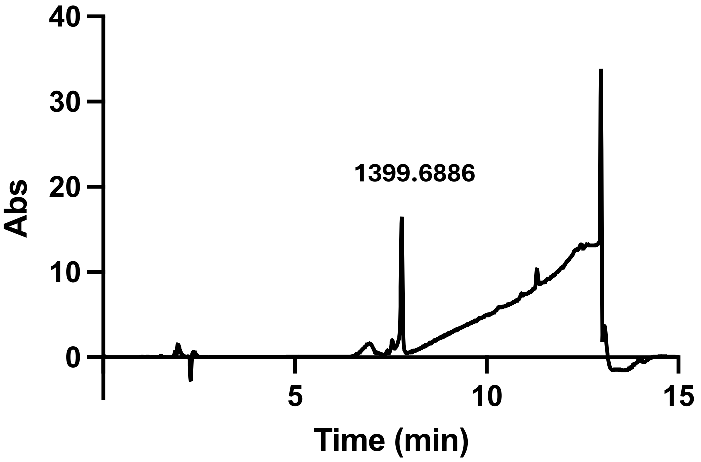

**Chemical formula: $C_{65}H_{94}N_{18}O_{13}S_2^+$**

**Calculated $[M + H]^+$ = 1399.70**

**Observed $[M + H]^+$ = 1399.6886**

**Compound 8**

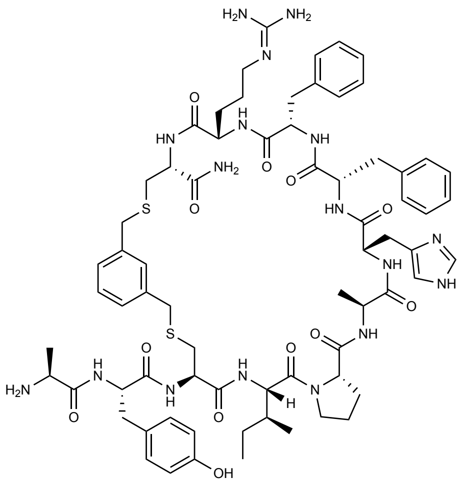

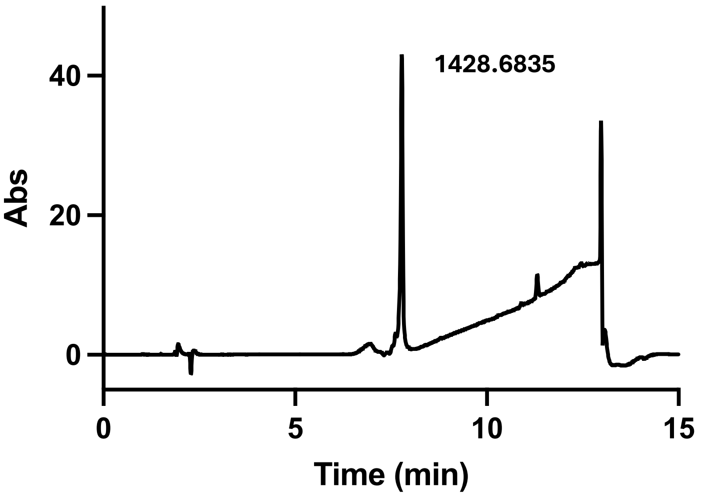

**Chemical formula: $C_{70}H_{93}N_{17}O_{12}S_2^+$**

**Calculated $[M + H]^+$ = 1428.74**

**Observed $[M + H]^+$ = 1428.6835**

**Compound 9**

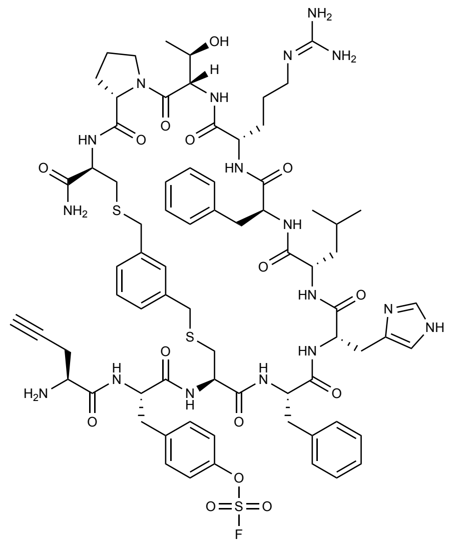

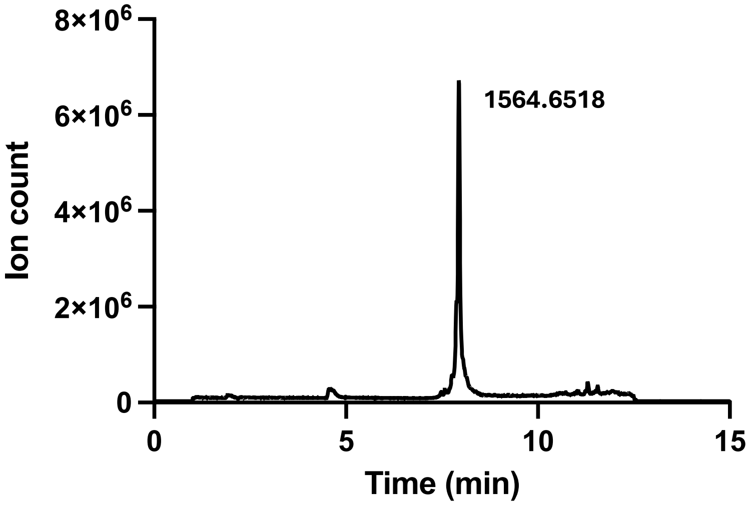

**Chemical formula: $C_{73}H_{94}FN_{17}O_{15}S_3^+$**

**Calculated $[M + H]^+$ = 1564.84**

**Observed $[M + H]^+$ = 1564.6518**

**Compound 10**

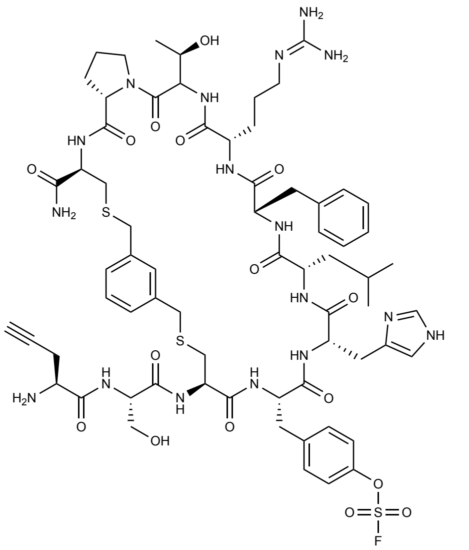

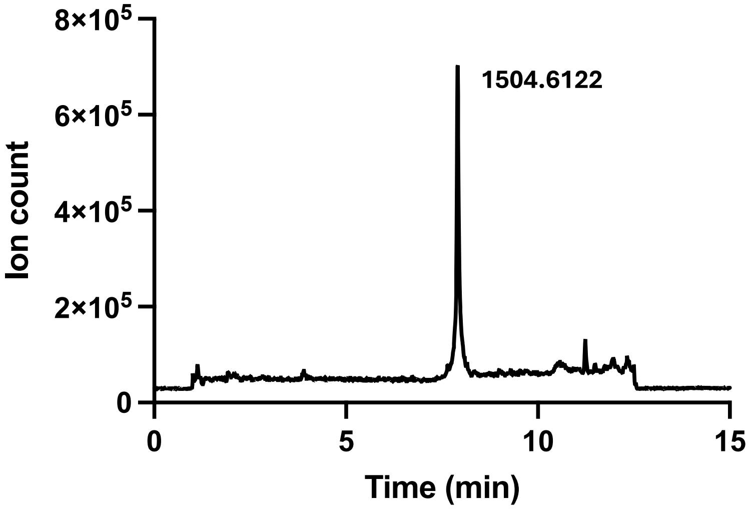

**Chemical formula: $C_{67}H_{90}FN_{17}O_{16}S_3^+$**

**Calculated $[M + H]^+$ = 1504.74**

**Observed $[M + H]^+$ = 1504.6122**

**Compound 11**

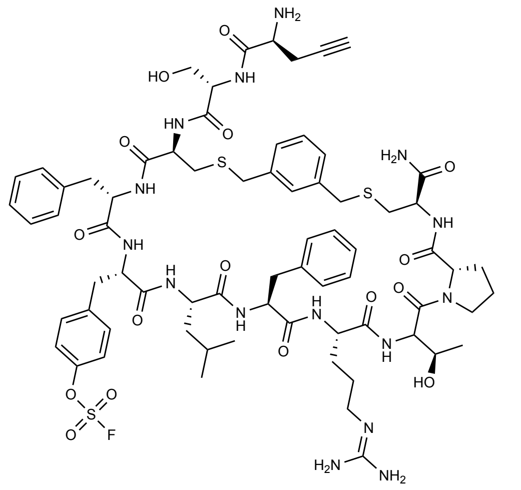

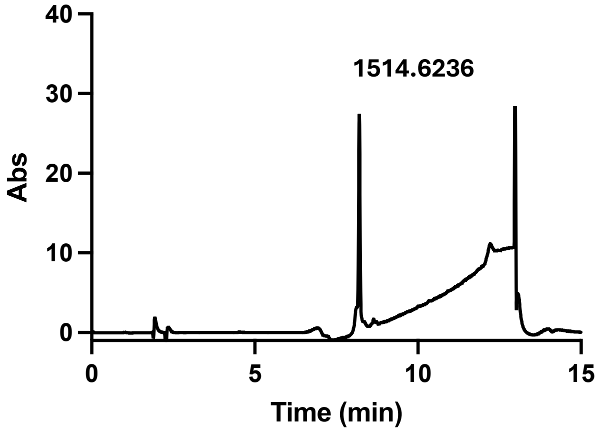

**Chemical formula: $C_{70}H_{92}FN_{15}O_{16}S_3^+$**

**Calculated $[M + H]^+$ = 1514.77**

**Observed $[M + H]^+$ = 1514.6236**

**Compound 12**

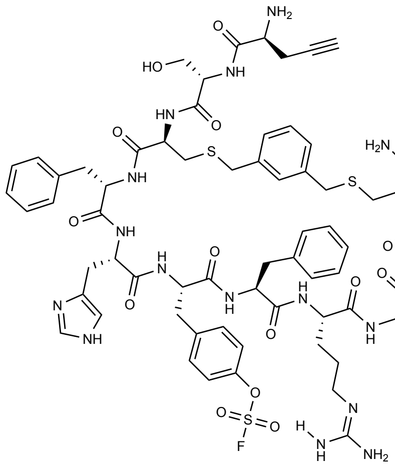

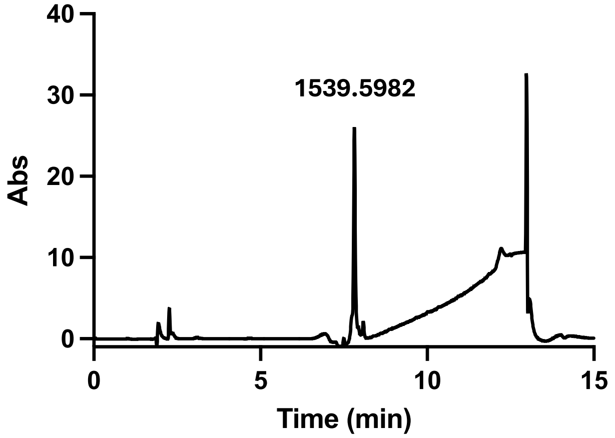

**Chemical formula: $C_{70}H_{88}FN_{17}O_{16}S_3^+$**

**Calculated $[M + H]^+$ = 1539.76**

**Observed $[M + H]^+$ = 1539.5982**

**Compound 13**

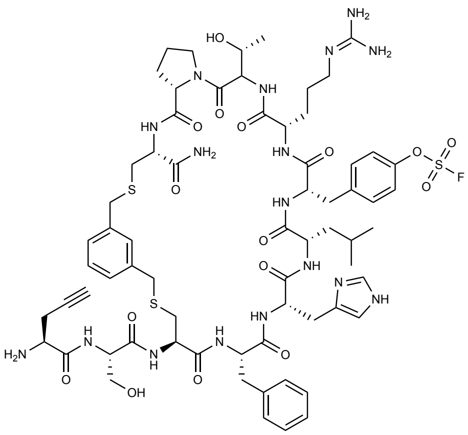

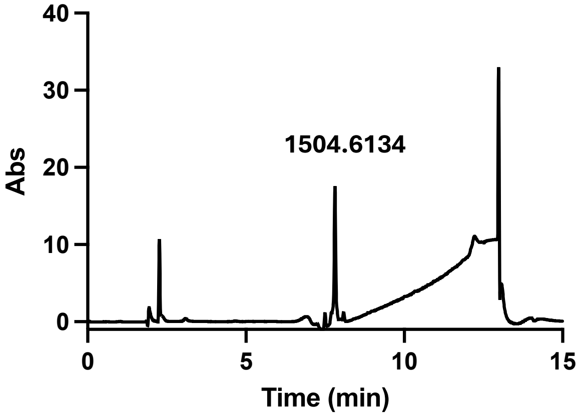

**Chemical formula: $C_{67}H_{90}FN_{17}O_{16}S_3^+$**

**Calculated $[M + H]^+$ = 1504.74**

**Observed $[M + H]^+$ = 1504.6134**

**Compound 14**

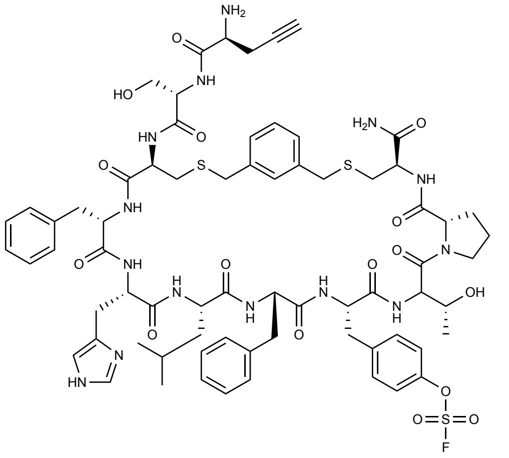

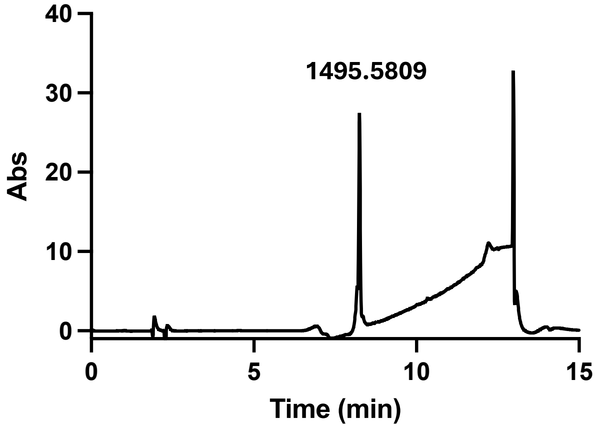

**Chemical formula: $C_{70}H_{87}FN_{14}O_{16}S_3^+$**

**Calculated $[M + H]^+$ = 1495.73**

**Observed $[M + H]^+$ = 1495.5809**

**Compound 15**

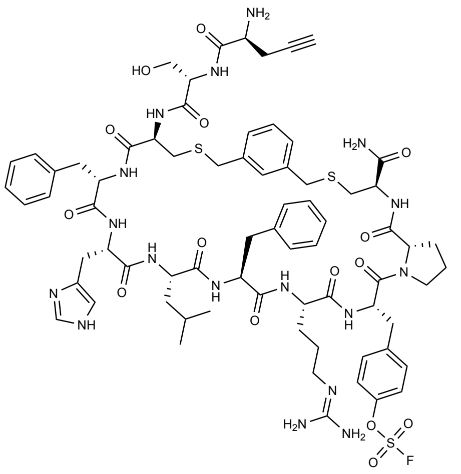

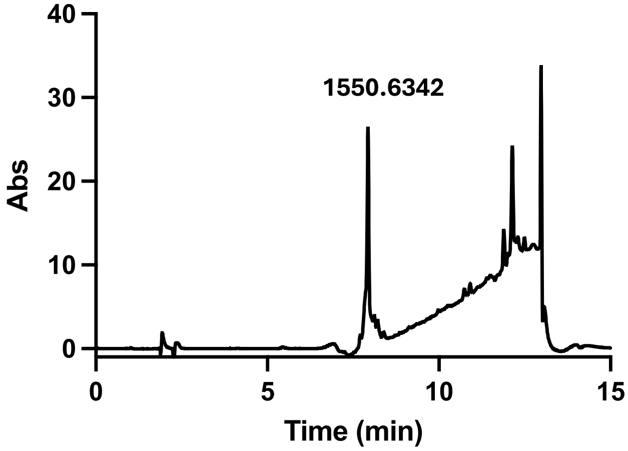

**Chemical formula: $C_{72}H_{92}FN_{17}O_{15}S_3^+$**

**Calculated $[M + H]^+$ = 1550.81**

**Observed $[M + H]^+$ = 1550.6342**

**Compound 16**

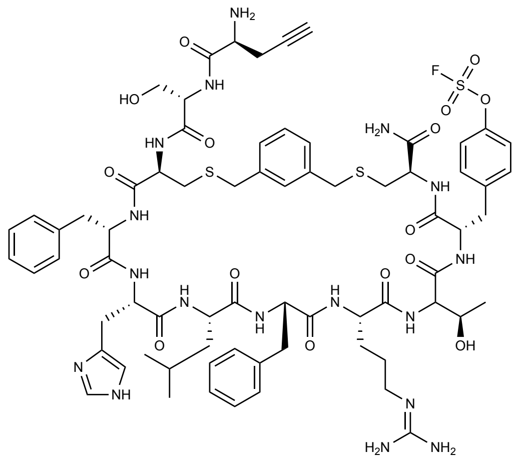

**1554.6287**

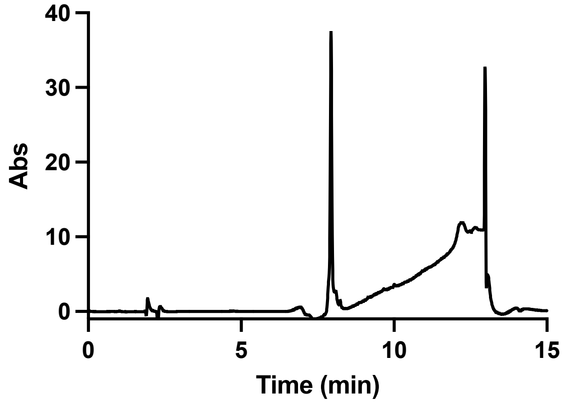

**Chemical formula: $C_{71}H_{92}FN_{17}O_{16}S_3^+$**

**Calculated $[M + H]^+$ = 1554.80**

**Observed $[M + H]^+$ = 1554.6287**

**Compound 17**

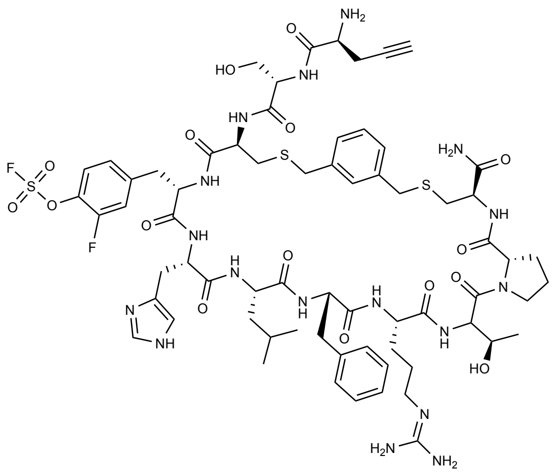

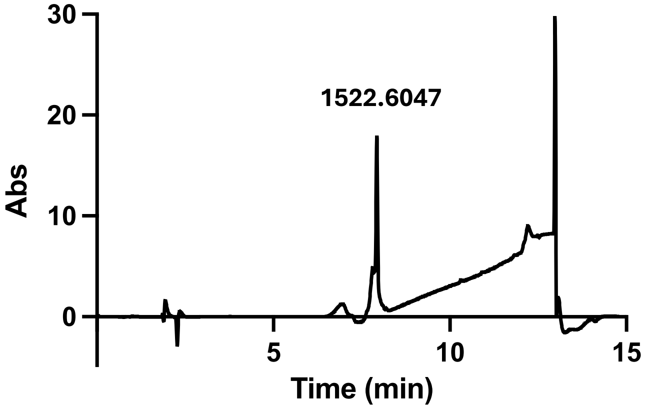

**Chemical formula: $C_{67}H_{89}F_2N_{17}O_{16}S_3^+$**

**Calculated $[M + H]^+$ = 1522.73**

**Observed $[M + H]^+$ = 1522.6047**

**Compound 18**

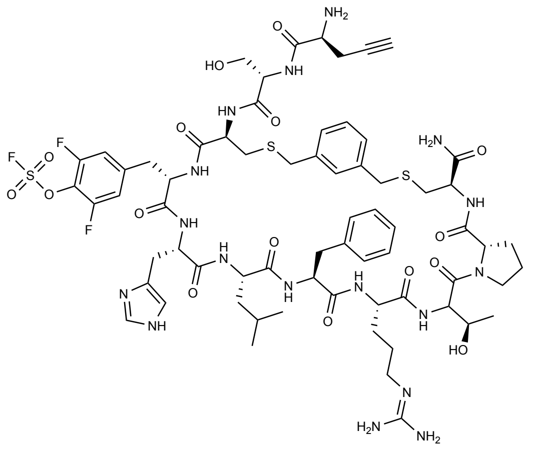

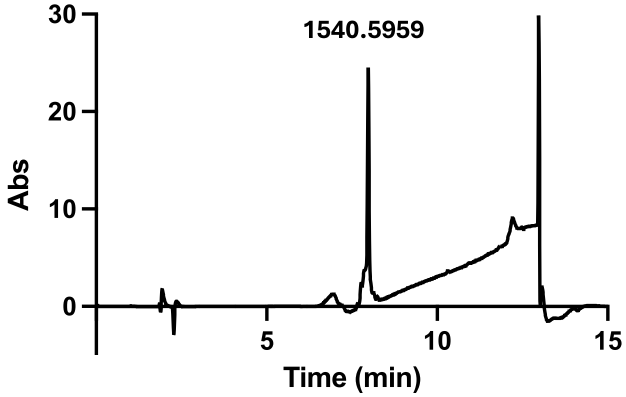

**Chemical formula: $C_{67}H_{88}F_3N_{17}O_{16}S_3^+$**

**Calculated $[M + H]^+$ = 1540.72**

**Observed $[M + H]^+$ = 1540.5959**

**Compound 19**

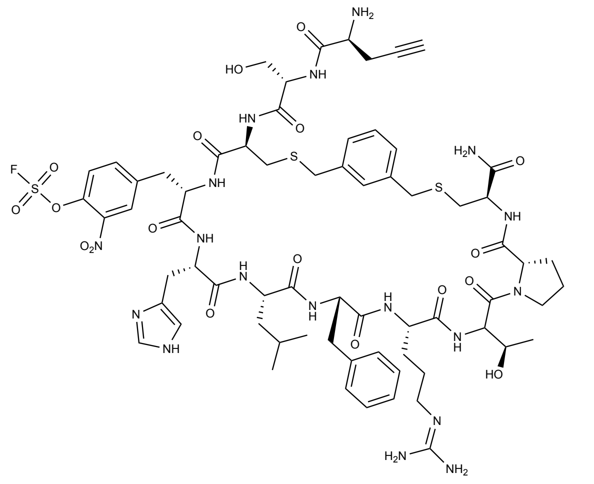

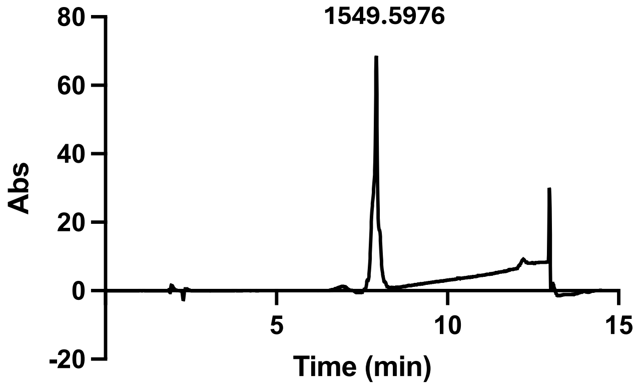

**Chemical formula: $C_{67}H_{89}FN_{18}O_{18}S_3^+$**

**Calculated $[M + H]^+$ = 1549.74**

**Observed $[M + H]^+$ = 1549.5976**

**Compound 20**

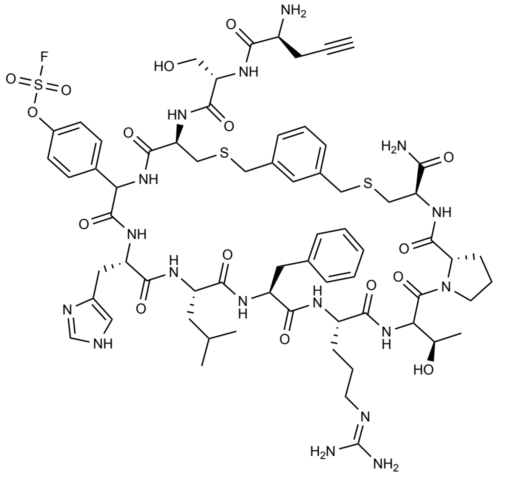

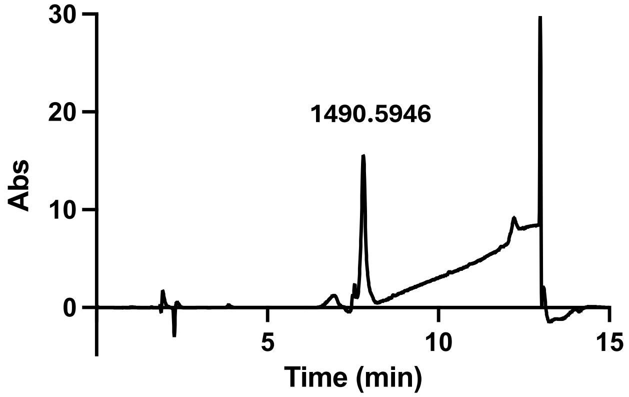

**Chemical formula: $C_{66}H_{88}FN_{17}O_{16}S_3^+$**

**Calculated $[M + H]^+$ = 1490.71**

**Observed $[M + H]^+$ = 1490.5946**

**Compound 21**

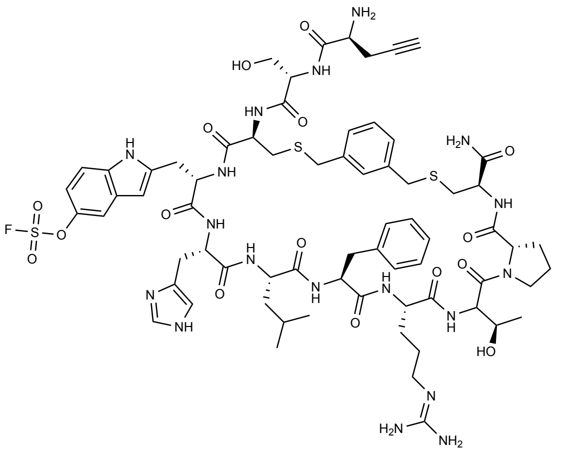

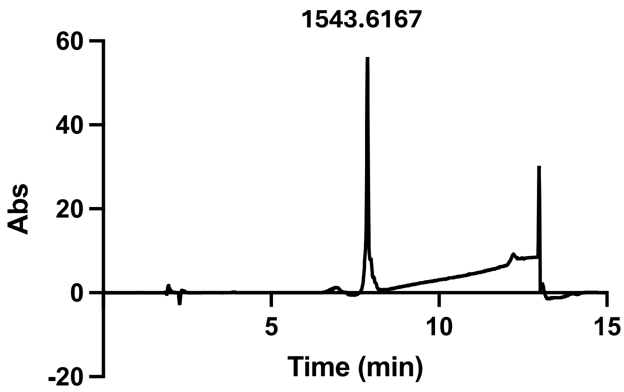

**Chemical formula: $C_{69}H_{91}FN_{18}O_{16}S_3^+$**

**Calculated $[M + H]^+$ = 1543.78**

**Observed $[M + H]^+$ = 1543.6167**

**Compound 22**

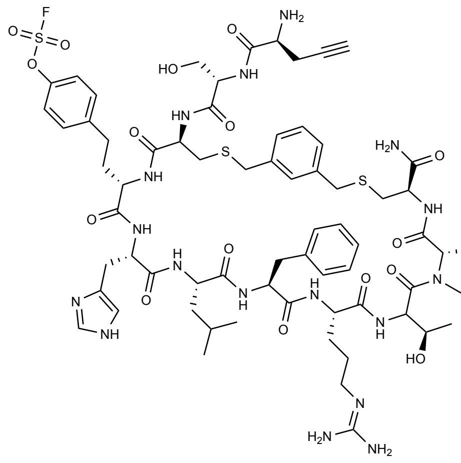

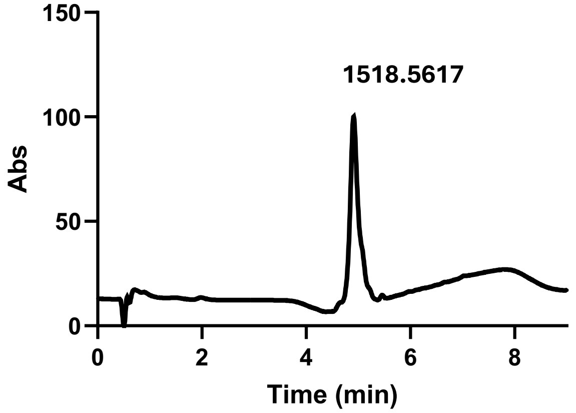

**Calculated [M + 2H]$^{2+}$ = 1518.56**

**Observed [M + 2H]$^{2+}$ = 1518.5617**

**Compound 23**

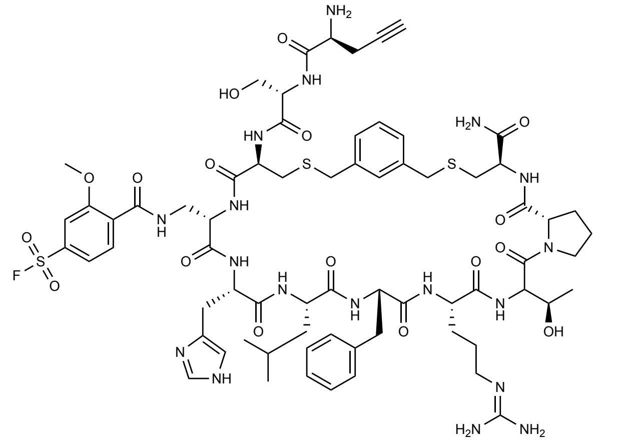

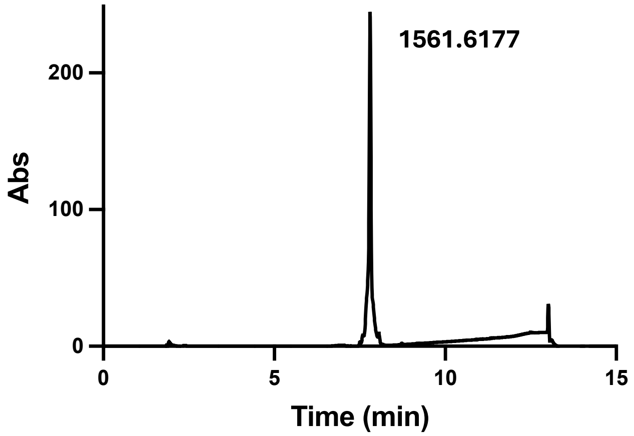

**Chemical formula: $C_{69}H_{93}FN_{18}O_{17}S_3^+$**

**Calculated $[M + H]^+$ = 1561.79**

**Observed $[M + H]^+$ = 1561.6177**

**Compound 24**

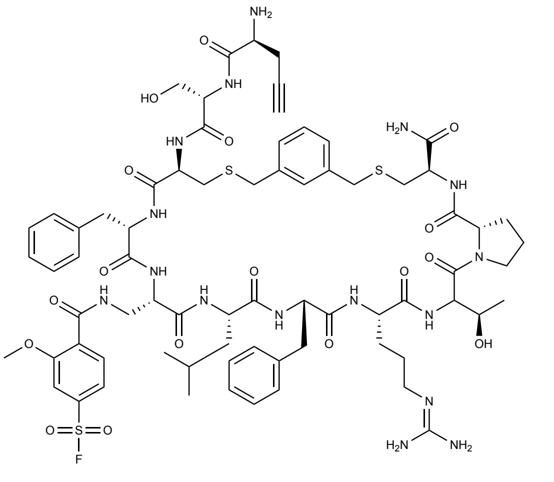

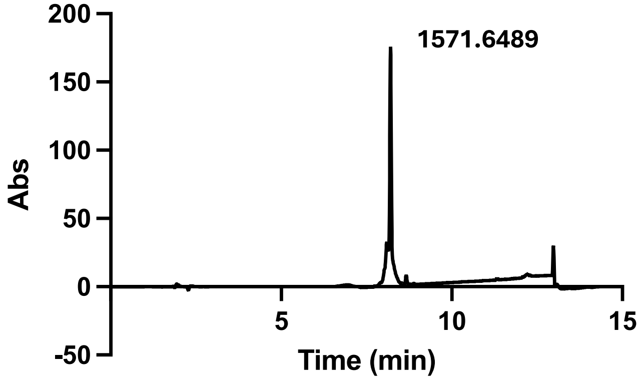

**Chemical formula: $C_{72}H_{95}FN_{16}O_{17}S_3^+$**

**Calculated $[M + H]^+$ = 1571.83**

**Observed $[M + H]^+$ = 1571.6489**

**Compound 25**

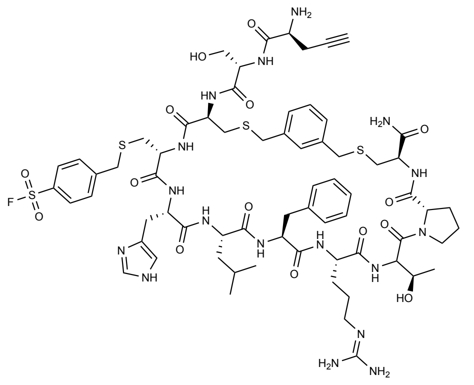

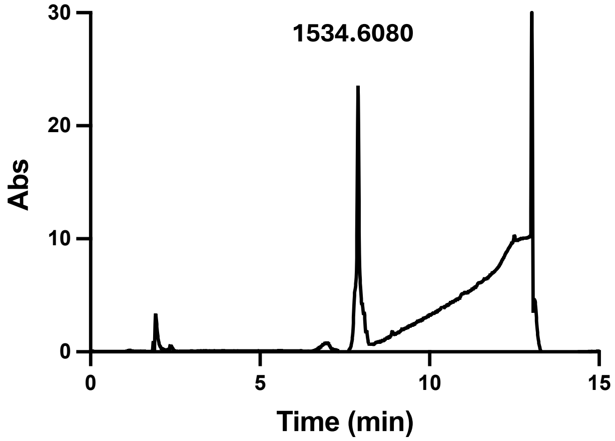

**Chemical formula: $C_{68}H_{92}FN_{17}O_{15}S_4^+$**

**Calculated $[M + H]^+$ = 1534.83**

**Observed $[M + H]^+$ = 1534.6080**

**Compound 26**

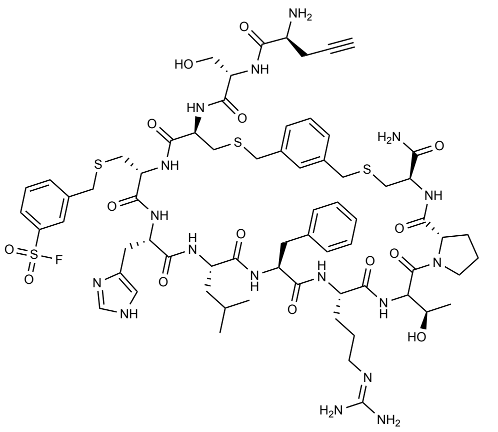

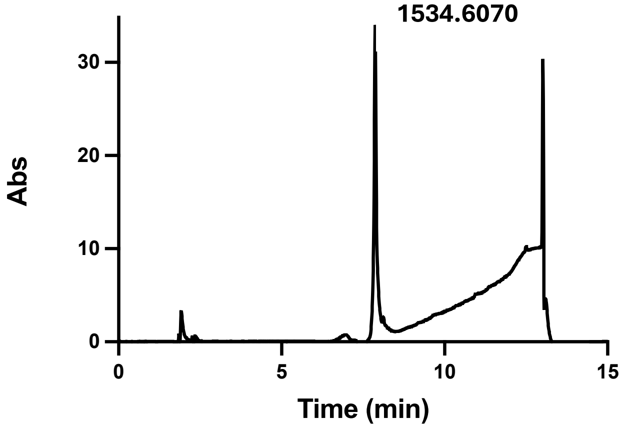

**Chemical formula: $C_{68}H_{92}FN_{17}O_{15}S_4^+$**

**Calculated $[M + H]^+$ = 1534.83**

**Observed $[M + H]^+$ = 1534.6070**

**Compound 27**

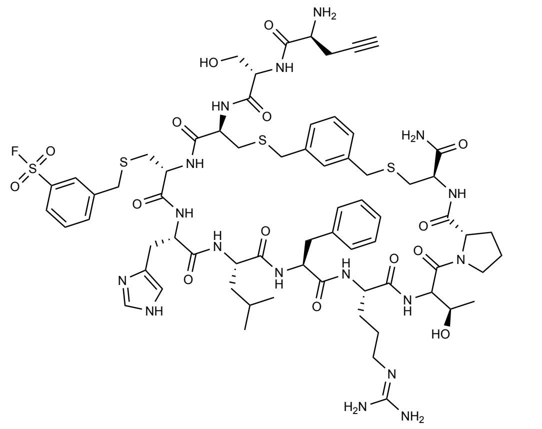

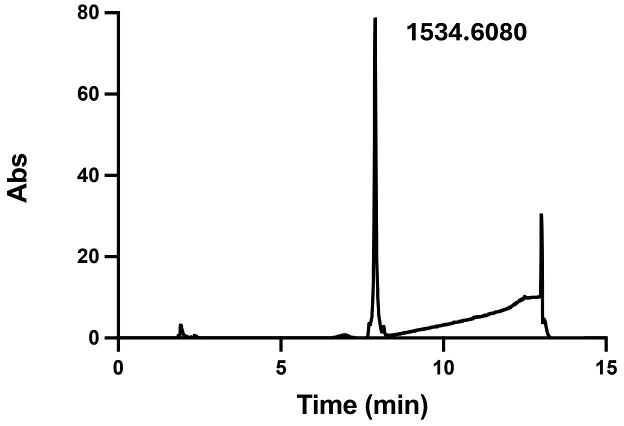

**Chemical formula: $C_{68}H_{92}FN_{17}O_{15}S_4^+$**

**Calculated $[M + H]^+$ = 1534.83**

**Observed $[M + H]^+$ = 1534.6080**

**Compound 28**

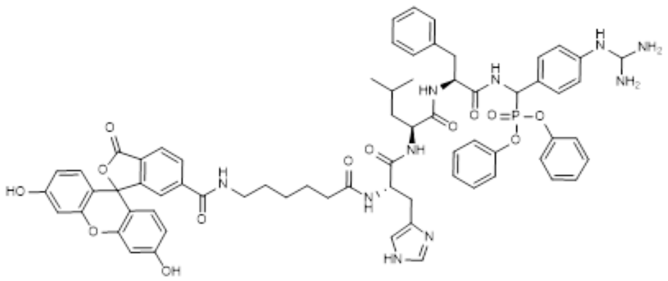

**Chemical Formula: $C_{67}H_{69}N_{10}O_{11}P$**

**Calculated $[M + 2H]^{2+}$ = 633.24**

**Observed $[M + 2H]^{2+}$ = 633.2466 // 633.2409**

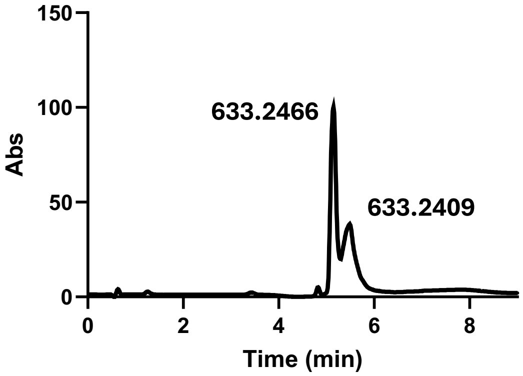

**Compound 29**

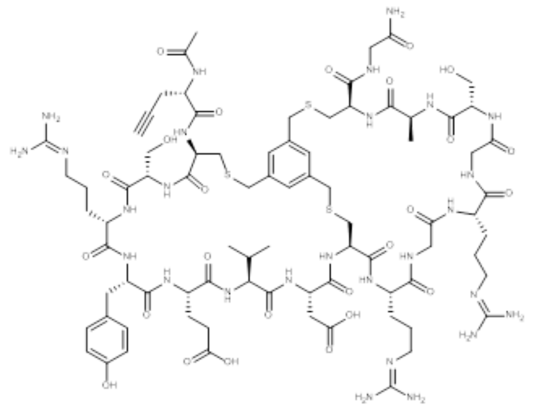

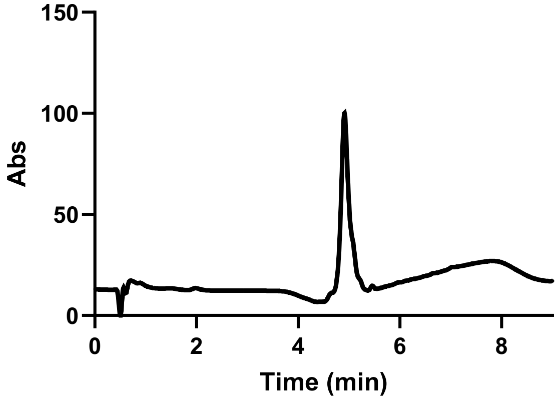

**Calculated $[M + 2H]^{2+}$ = 985.41**

**Observed $[M + 2H]^{2+}$ = 985.4191**

**Compound 30**

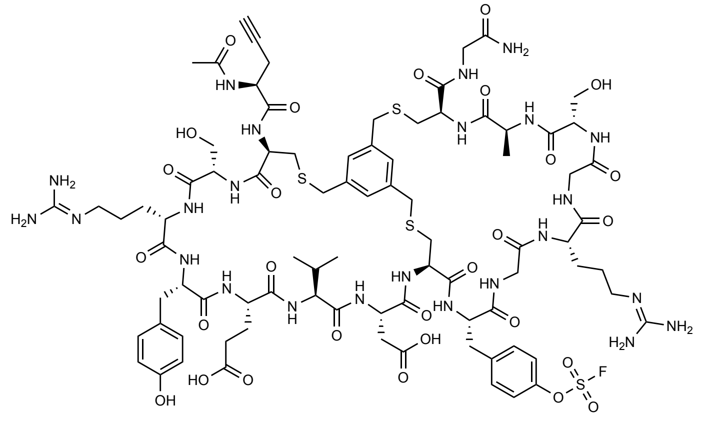

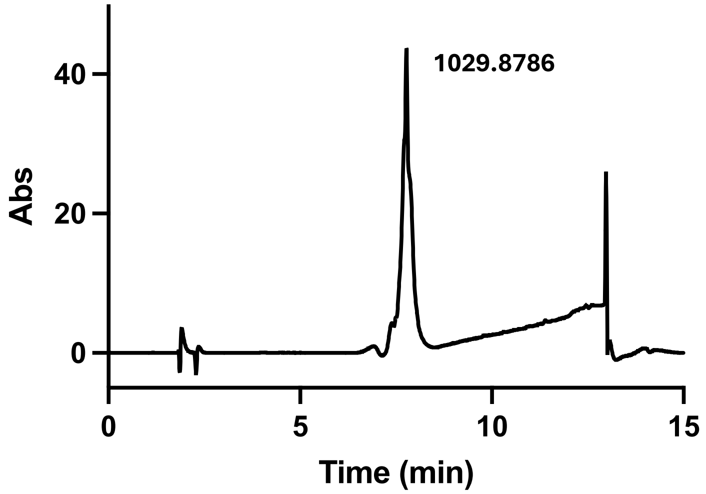

**Chemical formula: $C_{84}H_{117}FN_{24}O_{28}S_4^{2+}$**

**Calculated $[M + 2H]^{2+}$ = 1029.36**

**Observed $[M + 2H]^{2+}$ = 1029.8786**

**Compound 31**

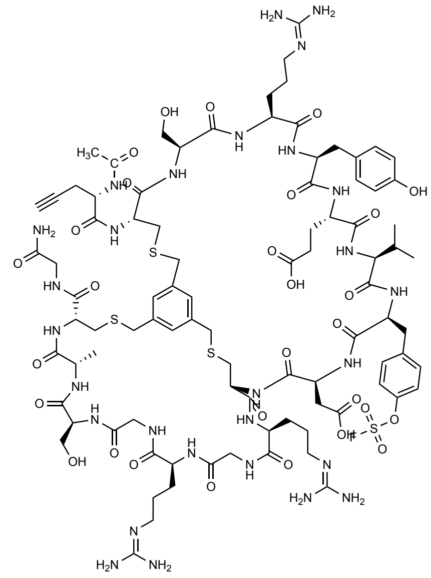

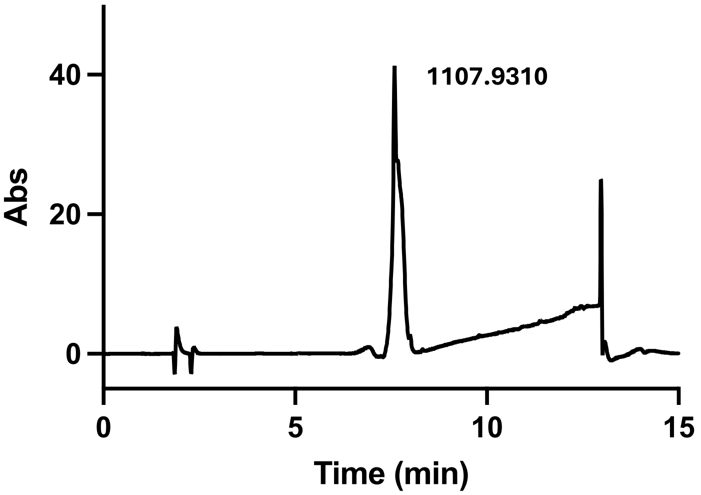

**Chemical formula: $C_{90}H_{129}FN_{28}O_{29}S_4^{2+}$**

**Calculated $[M + 2H]^{2+}$ = 1107.41**

**Observed $[M + 2H]^{2+}$ = 1107.9310**

**Compound 32**

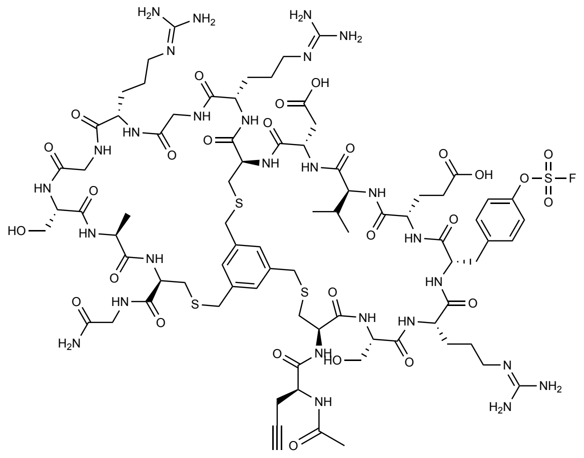

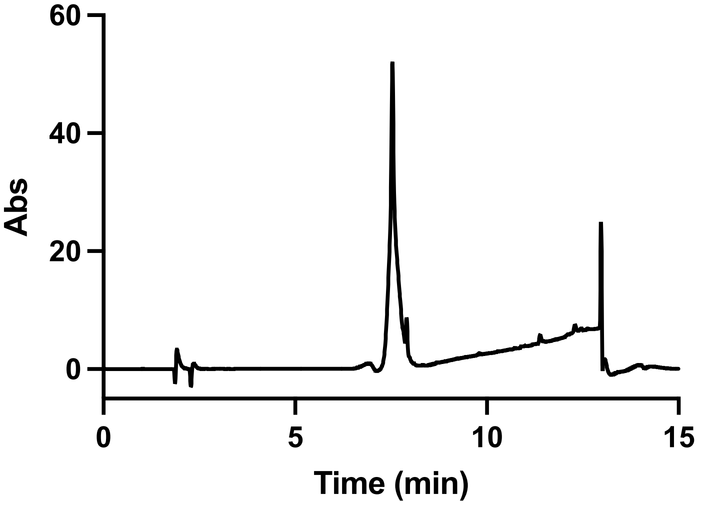

**$C_{81}H_{120}FN_{27}O_{27}S_4$**

**Calculated [M + 2H]$^{+2}$ 1026.39**

**Observed [M + 2H]$^{2+}$ = 1026.3946**

**Compound 33**

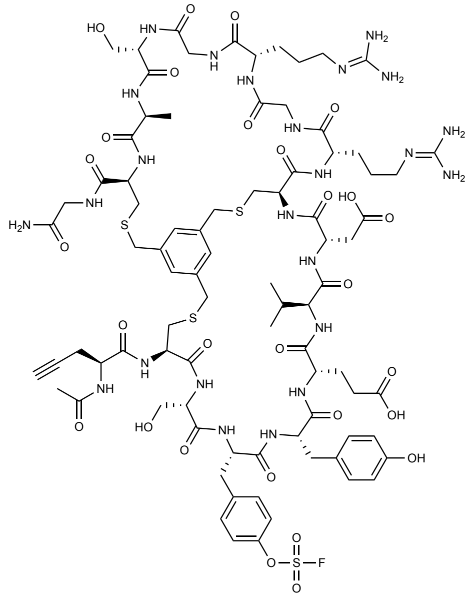

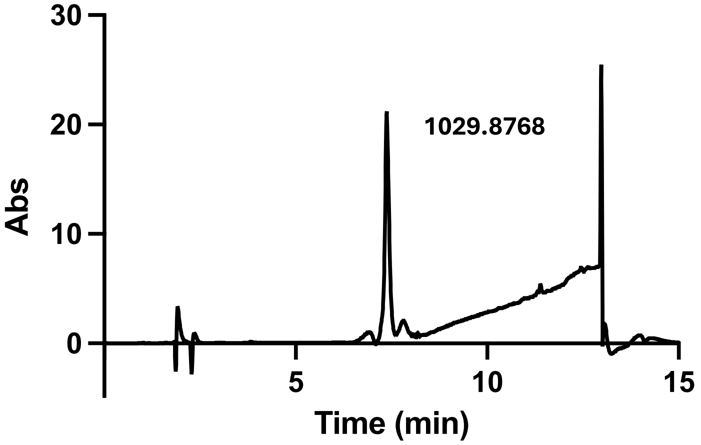

**Chemical formula: $C_{84}H_{117}FN_{24}O_{28}S_4^{2+}$**

**Calculated $[M + 2H]^{2+}$ = 1029.36**

**Observed $[M + 2H]^{2+}$ = 1029.8768**

### 6 References

1. S. Chen, S. Lovell, S. Lee, M. Fellner, P. D. Mace and M. Bogyo, Identification of Highly Selective Covalent Inhibitors by Phage Display, *Nat. Biotechnol.*, 2021, *39*, 490–498
2. V. Baeriswyl, H. Rapley, L. Pollaro, C. Stace, D. Teufel, E. Walker, S. Chen, G. Winter, J. Tite and C. Heinis, Bicyclic Peptides with Optimized Ring Size Inhibit Human Plasma Kallikrein and Its Orthologues while Sparing Paralogous Proteases, *ChemMedChem*, 2012, ***7***, 1173–1176.
